# Topovectorial mechanisms control the juxtamembrane proteolytic processing of Nrf1 to remove its N-terminal polypeptides during maturation of the CNC-bZIP factor

**DOI:** 10.1101/289785

**Authors:** Yuancai Xiang, Josefin Halin, Zhuo Fan, Shaofan Hu, Meng Wang, Lu Qiu, Zhengwen Zhang, Peter Mattjus, Yiguo Zhang

**Affiliations:** The Laboratory of Cell Biochemistry and Topogenetic Regulation, College of Bioengineering and Faculty of Sciences, Chongqing University, No. 174 Shazheng Street, Shapingba District, Chongqing 400044, China; Department of Biochemistry, Faculty of Science and Engineering, Åbo Akademi University, Artillerigatan 6A, III, BioCity, FI-20520 Turku, Finland; Institute of Neuroscience and Psychology, School of Life Sciences, University of Glasgow, 42 Western Common Road, G22 5PQ, Glasgow, Scotland, United Kingdom

**Keywords:** Nrf1, topovectorial regulation, regulated juxtamembrane proteolysis, ubiquitination, topobiology

## Abstract

The topobiological behaviour of Nrf1 dictates its post-translational modification and its ability to transactivate target genes. Here, we have elucidated that topovectorial mechanisms control the juxtamembrane processing of Nrf1 on the cyto/nucleoplasmic side of endoplasmic reticulum (ER), whereupon it is cleaved and degraded to remove various lengths of its N-terminal domain (NTD, also refold into a UBL module) and acidic domain-1 (AD1) to yield multiple isoforms. Notably, an N-terminal ∼12.5-kDa polypeptide of Nrf1 arises from selective cleavage at an NHB2-adjoining region within NTD, whilst other longer UBL-containing isoforms may arise from proteolytic processing of the protein within AD1 around PEST1 and Neh2L degrons. The susceptibility of Nrf1 to proteolysis is determined by dynamic repositioning of potential UBL-adjacent degrons and cleavage sites from the ER lumen through p97-driven retrotranslocation and -independent pathways into the cyto/nucleoplasm. These repositioned degrons and cleavage sites within NTD and AD1 of Nrf1 are coming into their *bona fide* functionality, thereby enabling it to be selectively processed by cytosolic DDI-1/2 proteases and also degraded *via* 26S proteasomes. The resultant proteolytic processing of Nrf1 gives rise to a mature ∼85-kDa CNC-bZIP transcription factor, which regulates transcriptional expression of cognate target genes. Furthermore, putative ubiquitination of Nrf1 is not a prerequisite necessary for involvement of p97 in the client processing. Overall, the regulated juxtamembrane proteolysis (RJP) of Nrf1, though occurring in close proximity to the ER, is distinctive from the mechanism that regulates the intramembrane proteolytic (RIP) processing of ATF6 and SREBP1.

## INTRODUCTION

It is known that nuclear factor-erythroid 2 (NF-E2)-related factor (Nrf1, also called Nfe2l1) belongs to the family of cap’n’collar (CNC) basic-region leucine zipper (bZIP) transcription factors. This family is involved in the regulation of metabolic, antioxidant, detoxification and cytoprotective genes against a vast variety of stresses (e.g. oxidants, xenobiotics, nutrients), pathophysiological stimuli (e.g. inflammation and aging) and other biological cues (e.g. metabolites, inducers and inhibitors) [1]. As far as we know to date, the CNC-bZIP family comprises the founding *Drosophila* Cnc protein, the *Caenorhabditis elegans* Skn-1, the vertebrate activator NF-E2 p45 subunit and related factors Nrf1, including its long TCF11 and short LCR-F (also called Nrf1β), Nrf2 and Nrf3, as well as transcription repressors Bach1 and Bach2 [2–6]. Amongst them, Nrf1 and Nrf2 are two important CNC-bZIP proteins in particular mammals. Nrf1 exerts the unique biological functions, but its loss of function results in specific pathophysiological phenotypes, which are distinctive from those of Nrf2 [1]. Such distinctions between Nrf1 and Nrf2 are principally determined by significant differences in both the primary structures of their gene products and their functional subcellular locations, where they are regulated through different mechanisms in distinct processes. Furthermore, it is also worth emphasizing a point that Nrf1, TCF11, Nrf3, CncC and Skn-1, but not Nrf2, are identified to form an NHB1 (N-terminal homology box 1)-CNC subfamily of membrane-bound transcription factors [7]. They have been determined to fold into a similar membrane-topology being integrated within and around the endoplasmic reticulum (ER) before being sorted out of the ER into the nucleus, in which the mature active isoform of CNC-bZIP transcription factor is enabled to bind its cognate gene promoters [7–10]. Of note, selective processing of Nrf1 (and its long TCF11) *via* various mechanisms yields multiple isoforms of between 140-kDa and 25-kDa, which exert different and even opposing biological activities. They coordinately regulate both basal and inducible expression of distinct subsets of antioxidant and/or electrophile-response elements (ARE/EpRE)-driven genes, that are responsible for homeostasis, development and cytoprotection against a variety of cellular stresses. However, detailed mechanism(s) by which isoforms of the CNC-bZIP protein are generated in distinct topovectorial processes remains elusive to date.

This being the case, it is of crucial importance to understand the intramolecular and subcellular basis for Nrf1 regulation. There is no doubt, as it has been shown for the topobiological model of Nrf1 [1], that the post-synthetic processing of the CNC-bZIP protein into various lengths of multiple polypeptide isoforms, e.g. Nrf1α, Nrf1β, Nrf1γ and Nrf1δ [11], with different or opposing functional potentials, is dictated by distinct topovectorial processes. This can allow Nrf1 (and its derivates) to fold into a proper topology within and around the ER membranes, in order to dynamically move in and out of the ER luminal side into extra-ER cyto/nucleoplasmic compartments, before being translocated to the nucleus and then gaining an access to target genes [1]. At first, upon translation of Nrf1, its NHB1 signal sequence (aa 1-30, Figure 1A) enables it to be anchored within ER membranes [12]. Subsequently, the *cis*-topovectorial process enables its DNA-binding CNC-bZIP domains to be positioned on the cyto/nucloplasmic side of membranes, whilst its transactivation domains (TADs, including AD1, AD2, NST & SR regions) are transiently translocated into the ER lumen. The NST domain of Nrf1 is N-glycosylated therein to yield an inactive glycoprotein of 120-kDa (estimated by NuPAGE) [13, 14]. Thereafter, when required for induction by biological cues, portions of the luminal-located TAD elements are dynamically retrotranslocated into extra-luminal subcellular compartments, where Nrf1 is deglycosylated to yield an active form of 95-kDa [7, 15]. The *trans*-topovectorial process monitors the selective proteolytic processing of Nrf1 so as to yield several membrane-released isoforms of between 85-kDa and 25-kDa by Hrd1-p97-mediated ubiquitin proteasome-dependent ERAD pathways [16–19]. However, no convincing evidence has been presented supporting the selective processing of Nrf1 by 26S proteasome-limited proteolysis, particularly in the intracellular ‘bounce-back’ response to low concentrations of proteasomal inhibitors [16, 20]. Therefore, this is a hot subject of debate with sensible doubtful points reported previously [18, 21, 22].

**Figure 1.**
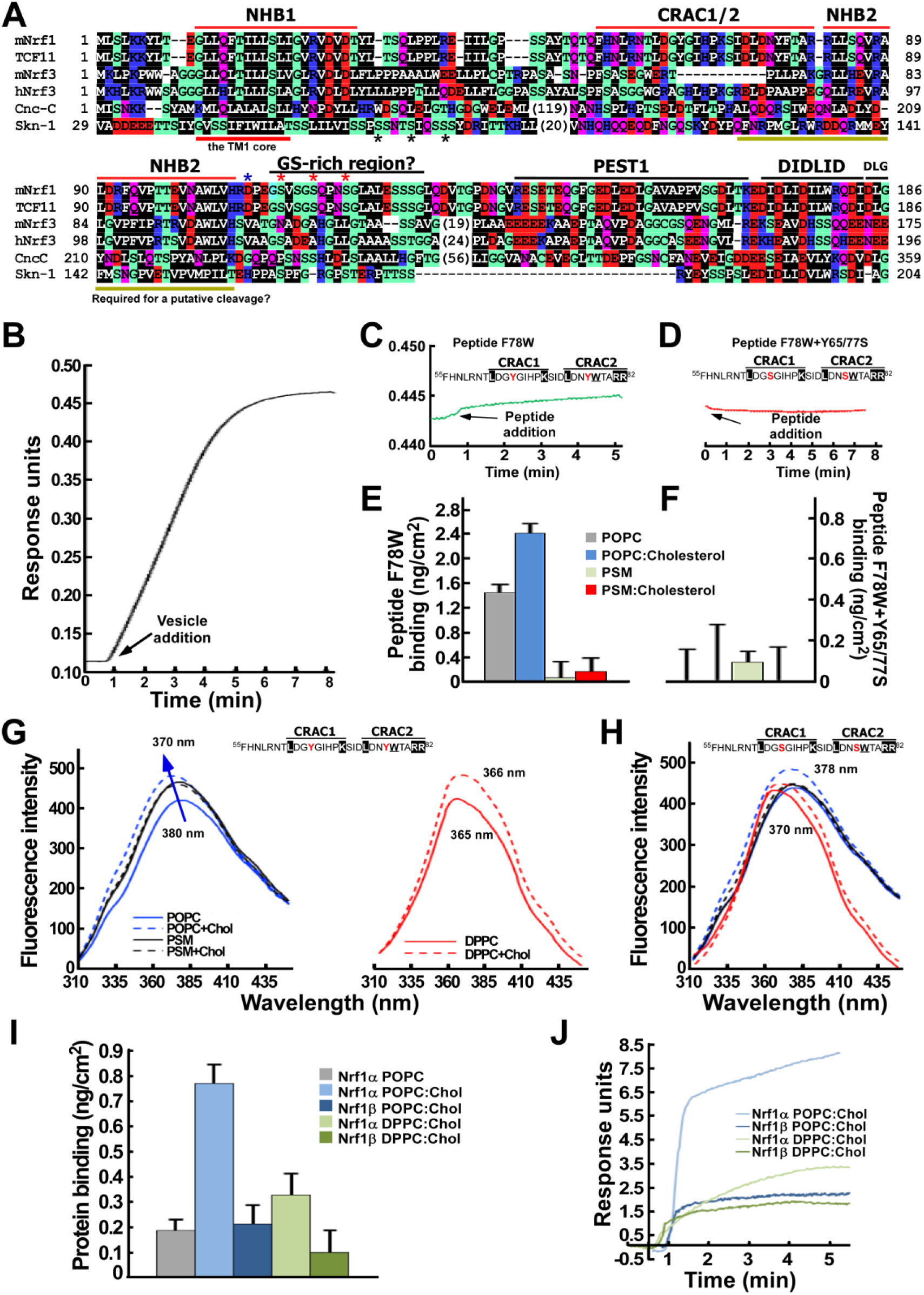
SPR analysis of cholesterol effects on binding of the Nrf1’s CRAC peptide to phospholipid vesicles. (**A**) Multiple sequence alignment of the N-terminal portions of indicated CNC-bZIP members, in which several major motifs are located, along with a putative cleavage site underlined (*yellow*). (**B**) A typical SPR sensogram showing the POPC:cholesterol (85:15 mol%) vesicle binding (*black trace*) to the sensor chip. (**C**) Addition of the CRAC^F78W^ (^55^FHNLRNTL DGYGIHPKSIDLDNY*<u>W</u>*TARR^82^) peptide to the sensor bound with the POPC:cholesterol vesicles causes an increase in the response units (i.e. mass increase) indicative of peptide binding (*green trace*). (**D**) Addition of the CRAC^F78W+Y65/77S^ (^55^FHNL RNTLDG*<u>S</u>*GIHPKSIDLDN*<u>SW</u>*TARR^82^) peptide causes no changes in the response units (*red trace*). Representative traces are shown from more than five independent experiments for each peptide. (**E**) The mass unit values for binding of the CRAC^F78W^ peptide to POPC or PSM vesicles containing zero or 15 mol% cholesterol, and (**F**) for the CRAC^F78W+Y65/77S^ peptide binding (note the scale difference between these two graphs) (mean ± S.D, n = 5). (**G**) The CRAC peptide normalized tryptophan fluorescence emission spectra in the presence of phospholipid vesicles with or without cholesterol. The *left panel* shows the tryptophan emission (λ_exc_ 295 nm) scans for the CRAC^F78W^ peptide with POPC vesicle added (solid blue scan), with emission maxima at 380 nm. The emission maximum blue-shifts with 10 nm to 370 nm when cholesterol containing POPC vesicles (25% cholesterol) were added (*dashed blue scan*). The emission maxima for the CRAC^F78W^ peptide did not shift and was at 376 nm both for PSM vesicle added (*solid black scan*) and the PSM:cholesterol vesicle added (*dashed black scan*). The *right panel* shows the emission scans and maxima for the CRAC^F78W^ peptide after addition of DPPC (365 nm, *solid red scan*) or DPPC:cholesterol (75:25, 366 nm, *dashed red scan*) vesicles. (**H**) The panel shows the tryptophan emission (λ_exc_ 295 nm) scans for the CRAC peptide with POPC vesicles added (*solid blue scan*) with an emission maxima remain at 378 nm, and the emission maxima with PSM vesicles added (*solid black scan*) at 378 nm. By contrast, the emission maxima remains at 378 nm when cholesterol containing POPC vesicles (25% cholesterol) are added (*dashed blue scan*), as well as after cholesterol containing PSM vesicles addition (*dashed black scan*). The red scans shows the emission maxima (370 nm) for the CRAC^F78W+Y65/77S^ peptide after addition of DPPC (*solid red scan*) or DPPC:cholesterol (75:25, *dashed red scan*) vesicles. The peptide concentration was defined at 2 μmol/L and the lipid concentration was 0.5 mmol/L. A representative scan is shown of at least 3 independent experiments. (**I**) Binding of Nrf1α and its short Nrf1β (lacking the N-terminal 298 aa of Nrf1α) to phosphatidylcholine (POPC or DPPC) vesicles with or without cholesterol (15 mol%) was measured with SPR. The SPR response was converted to mass using the approximation applied for proteins (1 RU = 0.0001° = 0.1 ng/cm^2^) according to the BioNavis protocol (mean ± S.D, n = 4). Error bars indicate SEM of more than three independent experiments. (**J**) Representative SPR sensogram showing the binding of the full-length Nrf1α or its shorter isoform Nrf1β to lipid vesicles composed of either POPC:cholesterol (85:15 mol fraction) or DPPC:cholesterol (85:15 mol fraction) used to calculate the data as described above.

Notably, no available evidence has been provided showing that a sufficient blockage of the proteasome activity leads to significant attenuation or complete abolishment in the proteolytic processing of intact Nrf1/TCF11. Their full-length glycoprotein and deglycoprotein of between 140-kDa and 120-kDa estimated on routine SDS-PAGE gels, are subjected again to separation by LDS-NuPAGE to migrate between 120-kDa and 95-kDa, respectively. However, this objective fact is manifest to contradict a recent report by Sha and Goldberg [18]. The authors assumed that potential proteasome-limited protealytic processing of Nrf1/TCF11 with a molecular mass of ∼100-kDa (resolved by SDS-PAGE gels) to give rise to a cleaved ∼75-kDa protein, seemed to be abolished by high doses of proteasomal inhibitors. The wrongly-estimated mass of ∼75-kDa protein was corrected to be approximately 90-95 kDa by same authors [22]. Importantly, it should also be noted that no evidence has been hitherto reported that the cytoplamic non-membranous 26S proteasome complex is integrated in close proximity of ER membranes. Rather, a similar process of both Site-1 and Site-2 proteases (i.e. SIP and S2P) has been accepted as a prerequisite for the regulated intramembrane proteolysis (RIP) accounting for ATF6 and SREBP1 [23]. More recently, a proteasome-independent mechanism involving DDI-1/2 has been shown to control the proteolytic processing of Nrf1 and Skn-1 [21, 24, 25], but the underlying mechanism remains obscure.

It is of critical importance to note that the selective post-translational processing of ectopic Nrf1 protein to yield cleaved polypeptides should be influenced by its net charged N-terminal and C-terminal epitopes in distinct topovectorial processes. According to the positive-inside rule and net charge differences across transmembrane (TM) regions [26–28], the strong negative charged 3×Flag sequence M<u>D</u>Y*K*<u>D</u>H<u>D</u>G<u>D</u>Y*K*<u>D</u>H <u>DID</u>Y*K*<u>DDDD</u>*K* [25, 29], two relative lower negative HA-tag (a nonapeptide YPY<u>D</u>VP<u>D</u>YA) [22, 24] and Myc-tag (a decapeptide <u>E</u>Q*K*LIS<u>EED</u>L) [30] are theoretically deduced to misdirect an improper membrane-topology of Nrf1, leading to an error-prone or even inverted orientation misfolded within ER membranes, as shown by [29]. As a matter of objective fact, accumulating evidence obtained from our group [12–14] and other workers on other membrane-proteins [28, 31] has revealed that the membrane-topology of Nrf1 is determined predominantly by its NHB1-associated TM1 peptide (aa 7-26). The TM1 is properly folded in the correct orientation of N_cyto_/C_lum_ (i.e. its N-terminal end faces the cytoplasmic side of ER membranes, whilst its C-terminal end faces the luminal side). Subsequently, the TM1-connecting CRAC1/2 (cholesterol-recognition/amino acid consensus 1/2, aa 55-82) and NHB2 peptides (81-106, Figure 1A), alongside with the TAD elements of Nrf1, are co-translationally translocated into the ER lumen, in which they are protected by membranes against cytosolic proteolysis [7, 14, 19]. For this reason, it is therefore postulated that some ER-resident portions of Nrf1 could be allowed for the possible proteolytic processing only after dynamic repositioning of them from the luminal side of membranes into extra-ER compartments. Therefore, these topovectorial processes of Nrf1 are inferred to dictate the selective proteolysis of the protein through its N-terminal domain [NTD, aa 1-124) and acidic domain-1 (AD1, aa 125-298). This is supported by the fact that most of NTD may also refold into a putative membrane-regulated ubiquitin-like (UBL) module, as described in DDI-1/2 and other UBL proteins (Figure S1A) [32, 33]. The UBL-adjacent AD1 is identified to possess additional three potential degrons, i.e. PEST1, DIDLID/DLG-adjoining Neh2L and CPD, besides its Neh5L subdomain essential for transactivation of Nrf1 [19]. If these putative degrons will be repartitioned and repositioned into the extra-ER side of membranes, they may target this CNC-bZIP protein to be selectively processed into distinct proteoforms.

To provide a better understanding of topovectorial mechanisms controlling selective proteolytic processing of Nrf1 to remove distinct lengths of within its putative UBL-adjoining regions, we have here determined: (i) whether the N-terminal CRAC1/2 peptide of Nrf1 situated between NHB1 and NHB2 within NTD enables the protein to bind the cholesterol-enriched membranes; (ii) which peptides within its N-terminal 298-aa region (N298, consisting of both NTD and AD1) are intrinsically required for distinct topovectorial processes, that monitor its juxtamembrane proteolysis on the extra-ER side of the membranes; (iii) which peptides within the N-terminal portions of Nrf1 have an ability to serve as a *bona fide* cleavage site for putative proteases (i.e. DDI-1/2 and 26S proteasomes) in order to remove different lengths of NTD or N298, when required for its topovectorial processing; and (iv) how p97-driven retrotranslocation and -independent pathways are involved in selective proteolytic processing of Nrf1 by cytosolic DDI-1 proteases and 26S proteasomes, so as to generate a mature CNC-bZIP factor contributing to transcriptional regulation of target genes.

## RESULTS

### The CRAC peptide within the N-terminal UBL-containing domain of Nrf1 enables the protein to directly bind the cholesterol-enriched membrane microdomains

Within NTD of mouse Nrf1 (mNrf1, Figure 1A), its amino acids 1 to 114, covering three major motifs NHB1 (essential for the ER-targeting of Nrf1 within membranes), CRAC1/2 and NHB2 (of which its C-terminal portion serves as an ER luminal-resident anchor[14]) were aligned with similar regions of 29 known UBL proteins, including DDI-1/2 (Figure S1A), as explained by [25, 32, 33]. Further bioinformatic analysis of NTD has identified that it shares structurally conserved homology with other UBL domains (Figure S1B), except for a unique TM1 α-helix (α_TM1_, which is folded by most of the NHB1 signal peptide and additional α_2_ (similar to equivalent of SB132 only, which is folded by an N-terminal part of the CRAC1/2-adjoining peptide of Nrf1) (Figure S1A). It is thus postulated that these two variable membrane-associated α_TM1_ and α_2_ peptides of Nrf1 may dictate dynamic refolding of its N-terminal region into a putative UBL module only after being released from the membranes into the cyto/nucleoplasmic compartments. Thereby, the UBL module of Nrf1 should be regulated by its membrane-topology, albeit it remains to be further studied in structural biology of the protein.

To provide an in-depth insight of CRAC-associated cholesterol-rich microdomains allowing for translocatition of Nrf1 across membrane lipid bilayers, the peptide-lipid binding experiments were firstly performed by employing surface plasmon resonance (SPR) spectroscopy, with immobilized vesicles on a hydrophobically modified dextran layer attached to a gold sensor chip. As a result, the difference in the binding of between two CRAC1/2 peptides was calculated, based on the level of the response units of the SPR sensogram at the end of the peptide injection (Figure 1, B to D). The binding of CRAC^F78W^ peptide to the POPC (1-palmitoyl-2-oleoyl-sn-glycero-3-phosphocholine) vesicles is significantly stronger in its binding to POPC:cholesterol (at a ratio of 85:15 (mol%)) vesicles (Figure 1E). By contrast, the binding of CRAC^F78W^ to other vesicles of PSM (N-palmitoyl-D-erythro-sphingosylphosphorylcholine) or PSM:cholesterol (at a ratio of 85:15) was negligible. Further examination also demonstrates that another mutant peptide CRAC^F78W+Y65/77S^ did not bind significantly to any vesicle types (Figure 1F).

Within Nrf1, its N-terminal CRAC1/2 sequence ^55^FHNLRNT*L*DG*Y*GIHP*K*SID*L*DN*Y***<u>W</u>**TARR^82^ (i.e. CRAC^F78W^, in which a tryptophan at position 78 was underlined, instead of phenylalanine in wild-type full-length Nrf1α) was employed to analyze the intrinsic emission properties of the peptide in the presence of membranes. Subsequently, the intrinsic tryptophan emission blue-shifts (10 nm) of the Nrf1 CRAC^F78W^ peptide were observed in the presence of cholesterol-containing POPC vesicles (Figure 1G). The parallel experiments were also conducted with additional point-mutant peptide ^55^FHNLRNTLDG**<u>S</u>**GIHPKSIDLDN**<u>S</u>**WTARR^82^ (i.e. CRAC^F78W+Y65/77S^, in which two underlined serines at position 65 and 77 substitute original tyrosines in order to influence its sensitivity of binding towards cholesterol). The results revealed that the tryptophan emission of CRAC^F78W+Y65/77S^ did not shift in the presence of either PSM or DPPC (1, 2-dipalmitoyl-sn-glycero-3-phosphocholine) vesicles containing cholesterol (Figure 1H). The difference in the tryptophan emission of between CRAC^F78W^ and CRAC^F78W+Y65/77S^ suggests that the tryptophan environment, which becomes more hydrophobic in the presence of cholesterol-containing POPC vesicles, is indicative of the peptide-membrane binding. This is consistent with the notion that the central Y65 residue of the CRAC motif plays a vital role in the cholesterol-binding, albeit other residues outside of this motif can also influence both the CRAC peptide-membrane interaction and its functionality [34]. In addition, no shift in the tryptophan emission was found when vesicles were composed of more tightly packed membranes, such as PSM and DPPC (Figure 1G).

Next, binding of the full-length Nrf1α or its shorter Nrf1β (lacking the N-terminal 298 aa of Nrf1α) to lipid vesicles was further analyzed by SPR spectroscopy. The results showed that intact Nrf1α strongly bound to vesicles composed of POPC:cholesterol, rather than DPPC:cholesterol, at the same ratio of 85:15 (mol%) (Figure 1I). By sharp contrast, a parallel experiment revealed no significant binding of the N-terminally-truncated Nrf1β to the same types of vesicles (Figure 1I,J). Together, the N298 portion of Nrf1α enables the intact full-length protein to be tightly tethered to the cholesterol-enriched microdomain of biomembranes through its CRAC sequence. The polypeptide-membrane interaction is hence inferable to be implicated in dynamic repartitioning of Nrf1 across cholesterol-enriched membranes, in order to partially move in and out of the ER lumen.

### Distinct peptides within the N298 portion of Nrf1 have antonymous effects on its AD1-mediated transactivation activity to regulate Gal4-target reporter gene expression

Clearly, the proper topology of Nrf1 is dominantly determined by its NHB1-adjoining TM1 peptide that adopts the orientation of N_cyto_/C_lum_ spanning the ER membranes [7, 14, 35]. The topovectorial folding of Nrf1 is, according to the positive-inside rule[26–28], enhanced by attachment of the net positively charged DNA-binding domain of the Gal4 transcription factor (i.e. Gal4D of 16.8 kDa, with a ratio of KR to ED = 26:20) to the N-terminal end of its N298 portion (to yield a fusion protein Gal4D-N298) (Figure 2A, *left panel*). In fact, our experimental evidence that has been provided previously [11, 13, 15] reveals that the basic N-terminal Gal4D epitope of Gal4D-N298 is partitioned on the cyto/nucleoplasmic side of membranes, whereas most of its C-terminal N298 portion is buried in the ER lumen. Such topological organization of Gal4D-N298 posits two questions of: i) whether the TM1-associated peptides (e.g. NHB1, CRAC and NHB2) and others within N298 were proteolytically processed, leading to disassociation of Gal4D from the C-terminal AD1 portion (acting as an essential TAD in Nrf1) and hence, if doing so completely, Gal4-target reporter gene expression is unable to be transactivated by AD1 in the context of Gal4D-N298 factor; ii) whether the AD1-adjoining regions are otherwise trapped in the ER lumen or retrotranslocated into the extra-luminal side of membranes. As if the luminal-resident AD1 element is solidly sequestered by membranes, it is also unable to trigger transactivation of the extra-luminal Gal4-target reporter gene by the Gal4D-N298 factor. Nevertheless, the reporter gene expression is *de facto* activated, though insufficiently, by Gal4D-N298 (Figure 2A, *row 1*). This implies that portions of AD1-adjoing regions in N298 are dynamically retrotranslocated across membranes and dislocated into the extra-ER cyto/nucleoplasmic compartments, allowing for an efficient access to transcriptional machineries before transactivating target reporter gene. The assumption is also supported by the data of Gal4*/UAS-luc* reporter assays showing a lower transactivation activity (∼100-fold changes) of intact Gal4D-N298 (i.e. Gal4D/Nrf1^1-298^), but this was accompanied by a maximum activity to ∼930-fold changes of Gal4-target reporter gene mediated by a membrane-free Gal4D/Nrf1^120-298^ (lacking the NTD of Nrf1 but retaining the entire AD1, *row 10*). This demonstrates that AD1-mediated transactivation is negatively regulated by NTD through association with the ER membranes, as consistent with previous reports [12, 35, 36]. Further examinations of four AD1-truncated mutants Gal4D/Nrf1^170-298^ (lacking the N-terminal PEST1-containing one-third region of AD1, *row 11*), Gal4D/Nrf1^188-298^ (lacking both the PEST1-containing region and the DIDLID/DLG element, *row 12*), Gal4D/Nrf1^170-242^ (containing the Neh2L subdomain alone, *row 15*) and Gal4D/Nrf1^280-298^ (only containing the Neh5L subdomain, *row 14*) indicated that the DIDLID/DLG element of Neh2L, besides Neh5L, contributes to AD1-mediated transactivation in the Gal4D-N298 fusion context.

**Figure 2.**
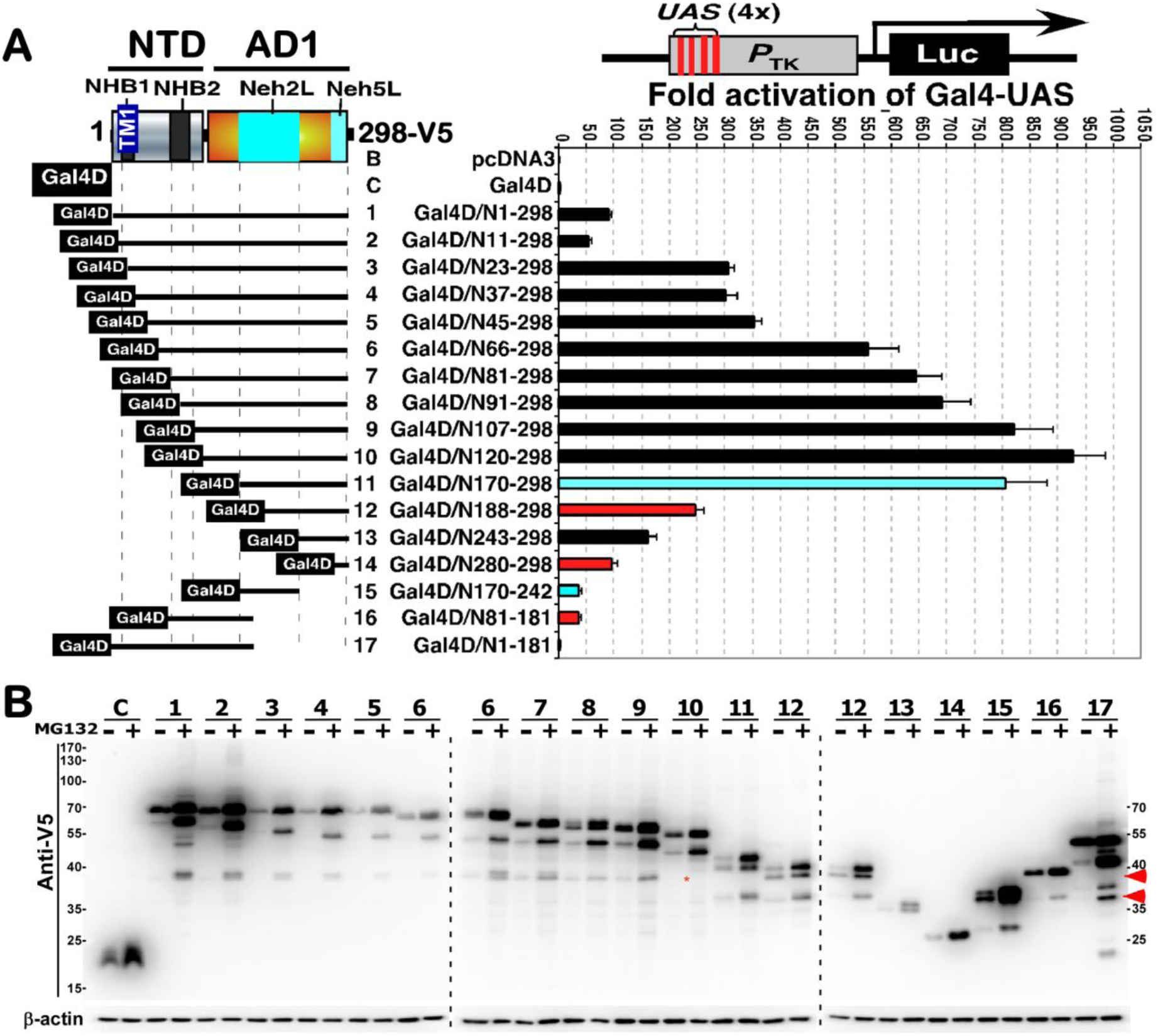
Distinct regions within the N298 portion of Nrf1 in Gal4 fusion proteins have antonymous effects on either its transactivation activity or its post-synthetic processing into multiple proteoforms. (**A**) Schematic of a series of Gal4D fusion proteins with N298 or its truncated mutants (*left panel*), with locations of NHB1, NHB2, Neh2L and Neh5L being indicated. The *right panel* shows the Gal4/UAS-driven transcriptional activity mediated by Gal4D/N298 chimeric factors or its mutants. The luciferase reporter activity was measured in lysates of COS-1 cells that had been transfected with each of expression constructs for Gal4D/N298 and its mutants, together with *P_TK_-UAS/Gal4*-*Luc* and *β-gal* plasmids. The resulting data were calculated as a fold change (mean ± S.D, n = 9) relative to the control Gal4D level after normalization to the β-gal value. (**B**) The transfected COS-1 cells were treated with MG132 (+ 5 µmol/L) or not (−) for 4 h and then lysized in a denature buffer. Then, these protein extracts were resolved by SDS-PAGE containing 10% polyacrylamide and visualized by immunoblotting with anti-V5 antibody. The single letters *B* and *C* indicate a basal background and another control levels, obtained from those cells that had been transfected with an empty pcDNA vector or only Gal4D-expression construct, plus the indicated reporter genes, respectively.

To explore significant effects of NHB1, NHB2 and CRAC, along with putative UBL-adjoining peptides, on the transactivation function of AD1, a series of expression constructs were created by fusing Gal4D at the C-terminus to various lengths of progressively N-terminally-truncated mutants of N298. As anticipated, AD1-mediated transactivation is inhibited by adjacent UBL-containing NTD, because deletion of the NTD from within Gal4D-N298 (to yield Gal4D/Nrf1^120-298^) resulted in a maximal ∼9-time increase in transactivation of Gal4*/UAS-luc* reporter gene, when compared with that obtained from intact Gal4D/Nrf1^1-298^ (Figure 2A, *rows 10 vs 1*). Surprisingly, transfection of the construct that directed expression of the fusion protein lacking residues 1-10 from the NTD, but with the entire AD1 being retained in Gal4D/Nrf1^11-298^, was less able to transactivate *UAS*-driven gene expression than the intact Gal4D/Nrf1^1-298^ factor (*rows 1 vs 2*). This suggests that the N-terminal 10 residues (i.e. the n-region of ER-targeting peptide) of Nrf1 might antagonize negative regulation of AD1 by NTD or contribute to topovectorial regulation of AD1-mediated transactivation. By comparison of Gal4D/Nrf1^11-298^, Gal4D/Nrf1^23-298^ and Gal4D/Nrf1^37-298^ (*cf. rows 2-4*), the results revealed that the N-terminal residues 11-22 (covering the ER-targeting h-region), but not residues 23-36 (covering the ER-targeting c-region) within and around the TM1-containing NHB1 sequence contribute essentially to the negative regulation by NTD. Further comparisons of other mutants between Gal4D/Nrf1^37-298^ and Gal4D/Nrf1^120-298^, along with those intermediary deletions (*cf. rows 5-10*), indicated that residues 37-119 (covering the CRAC1/2 and NHB2 sequences within putative UBL) also contribute substantially to the negative regulation of AD1 by NTD. However, it should be noted that such striking changes in their transactivation activity did not appear to be attributed to differences in total levels of the above-examined fusion proteins (Figure 2B, *cf. lanes 1-10*). Overall, these data demonstrate that the core h-region (aa 11-22) of TM1-adjoining NHB1 peptide, and the whole NHB2 (aa 81-106) peptide, as well as the CRAC1/2-adjoining sequence between NHB1 and NHB2, are essential for negative stepwise regulation of AD1 by NTD. This conclusion is further supported by additional data obtained from both Gal4D/Nrf1^81-181^ and Gal4D/Nrf1^1-181^, showing that the N-terminal residues 1-80 of Nrf1 (only covering the NHB1 and CRAC1/2-adjoining peptides) completely prevented *UAS-Luc* reporter gene transactivation by an N-terminal one-third portion of AD1 (Figure 2A, *cf. rows 16 with 17*).

It is important to note that the first N-terminal 50-aa region of Gal4D is essential for its DNA-binding to the *UAS-Luc* reporter gene, whilst AD1-mediated transactivation is determined by the last C-terminal Neh5L tail of AD1, besides its DIDLID/DLG element, within the N298 portion [11, 15, 19]. The fact demonstrates that only after the ER luminal-resident portions of AD1 fused in the context of Gal4D-N298 factor is dynamically repartitioned and then dislocated into extra-ER cyto/nucleopasmic compartments, they may be allowed for transactivation of Gal4-target reporter gene. Conversely, upon truncation of the N-terminal Gal4D or C-terminal TAD elements from intact fusion protein Gal4D-N298, it will have lost its constructive transactivation activity. Therefore, a large majority of multiple degraded polypeptides arising from the Gal4D-N298 processing should be theoretically non-functional, no matter whether they have been recovered or not in the ER membrane factions [11, 15].

### The N298 portion of Nrf1 directs the processing of Gal4 fusion protein to yield multiple polypeptides

Western blotting of proteins, separated by SDS-PAGE gels containing 10% polyacrylamide and then visualized with antibodies against their C-terminally-tagged V5 epitope (Figures 2B and S2), showed that Gal4D/Nrf1^243-298^ gave rise to two weak bands migrated close to 35-kDa (Figure 2B, *lane 13*), and Gal4D/Nrf1^280-298^ was represented by a single weak band of 25-kDa (*lane 14*). Yet, their proteins were obviously accumulated by proteasomal inhibitor MG132. Together with the previous data [19, 37], these indicate that Neh5L-adjoining Cdc4-phosphodegron (CPD) of Nrf1 may target Gal4-fusion proteins to proteasome-mediated degradation pathway. Intriguingly, an additional minor polypeptide band of ∼26-kDa that seems similar to Gal4D/Nrf1^280-298^ was also resolved by electrophoresis of Gal4D/Nrf1^170-242^ (*lanes 15 vs 14*), as was accompanied by two major closer polypeptides migrated to ∼36/37-kDa. It is hence deduced that the unstable ∼26-kDa polypeptide is yielded possibly through Neh2L-triggered proteolytic processing of the fusion protein within its Gal4D epitope and its yield was promoted by MG132. Similar proteolytic processing event also appeared to emerge in additional two cases of Gal4D/Nrf1^170-298^ and Gal4D/Nrf1^188-298^ (*lanes 11 & 12*). This is supported by the finding that expression of their fusion proteins also gave rise to an extra minor polypeptide of ∼36-kDa, which migrated close to the molecular mass of Gal4D/Nrf1^243-298^, besides two major bands of between 39-kDa and 42-kDa juxtaposed.

Further examination revealed that Neh2L-adjoining residues 81-181 of Nrf1 (covering its NHB2, the PEST1 degron and DIDLID/DLG sequences) are also involved in putative proteolytic processing of Gal4D/Nrf1^81-181^ to yield an additional cleaved ∼36-kDa polypeptide, besides a major polypeptide of ∼39-kDa (Figures 2B and S2, *lane 16*). By contrast, a multiple polypeptide ladder of between 55-kDa and 20-kDa was clearly exhibited following protein resolution by SDS-PAGE of total lysates of COS-1 cells expressing Gal4D/Nrf1^1-181^ (containing both putative UBL and PEST1-adjoining DIDLID/DLG sequences, *lane 17*). This, together with the previous-reported data [15], suggests that putative UBL and PEST1 degrons may be allowed for the proteolytic processing of Gal4D/Nrf1^1-181^ to yield multiple polypeptides; small forms of ∼20/23-kDa were postulated to arise from the proteolysis of fused Gal4D *per se*. This notion is further supported by additional patterns of multiple polypeptide ladders composed of between 70-kDa and 25-kD, which occurred in the cases of Gal4D/Nrf1^1-298^ and Gal4D/Nrf1^11-298^ (*cf. lanes 1 & 2 with 17*). Furthermore, all other cases also gave rise to similar but nuanced multiple peptide ladders with two to three major electrophoretic bands, and their occurrence appeared to be unaffected by sequential deletions of the N-terminal 11- to 187-aa from within the N298 portion of the fusion proteins (*cf. lanes 2 to 12*). Collectively, these indicate that Neh2L and/or CPD degrons may also target AD1-retaining fusion proteins to proteolytic degradation.

In each case of progressively increasing deletions to yield different lengths of the N-terminal portion of N298, subsequent proteolytic processing of these resulting fusion proteins within extra-Gal4D portions is also inferred to give rise to short polypeptides between 39-kDa and 25-kDa. Interestingly, the ∼39-kDa polypeptide was prevented by both mutants Gal4D/Nrf1^120-298^ (lacking putative UBL) and Gal4D/Nrf1^170-298^ (lacking both UBL and PEST1), and remained to be silent even in the presence of MG132 (Figure 2B, *lanes 10 & 11*). This finding implies that a putative proteolytic cleavage event to yield the 39-kDa polypeptide may occur in the C-terminal border of UBL overlapping with PEST1. In addition, variable abundances of the above processed polypeptides were determined in different experimental settings (Figures 2B and S2). Amongst them, the fusion proteins along with their derivates particularly from TM1-disrupted mutants (*lanes 3 to 16*) are unstable due to the fact that they are unprotected by membranes and thus rapidly proteolytically degraded by proteasomes [11, 15, 19]. Of note, it cannot be ruled out that another possibility that many of the processed polypeptides (i.e. more than ∼39 kDa in *lanes 1 to 9*, ∼36 kDa in *lanes 11 & 12*, and ∼26 kDa in *lane 15*) may also have undergone some modifications, such as ubiquitination, after the proteolytic processing of within Nrf1-derived sequence.

### Various lengths of N-terminal polypeptides arise from the putative proteolytic processing of Nrf1 fusion proteins within its NTD and N298 portions, even in the presence of proteasomal inhibitors

To determine whether the above-mentioned degrons direct the selective proteolytic processing of Nrf1 *per se* or its fusion portions through its NTD alone or plus AD1 within N298, we constructed a series of chimeric proteins. In the schematic diagrams (Figure 3A), Nrf1, its NTD and N298 was respectively sandwiched in the middle place, and flanked C-terminally by enhanced green fluorescent protein (eGFP) and N-terminally by net zero, neutral or slightly positive charged tags, such as V5, VSVg or StrepII (*left panel*). Notably, an extra positive residue arginine (R) was inserted in between each tag and the N-terminal end of Nrf1, such that the proper topology of the protein will be maintained due to its correct folding within and around ER membranes, as abided by the positive-inside rule of topology [26–28]. The resultant sandwiched proteins V5-Nrf1-eGFP (Figure 3,B to D), as well as VSVg-Nrf1-eGFP and StrepII-Nrf1-eGFP (Figure S3) were resolved by SDS-PAGE gels containing 10% polyacrylamide. Then, Western blotting results showed that their electrophoretic bands representing these fusion proteins and derivates were weakly visualized by antibodies against each of their indicated tags V5, VSVg, StrepII or GFP, but strongly immunoblotted with specific antibodies recognized the C-terminal residues 291-741 of mNrf1 (i.e. Nrf1β). This suggests that a fraction of the chimeric protein is proteolytically processed leading to removal of diverse lengths of its N-terminal and/or C-terminal portions from Nrf1, leading to the production of multiple processed polypeptides.

**Figure 3.**
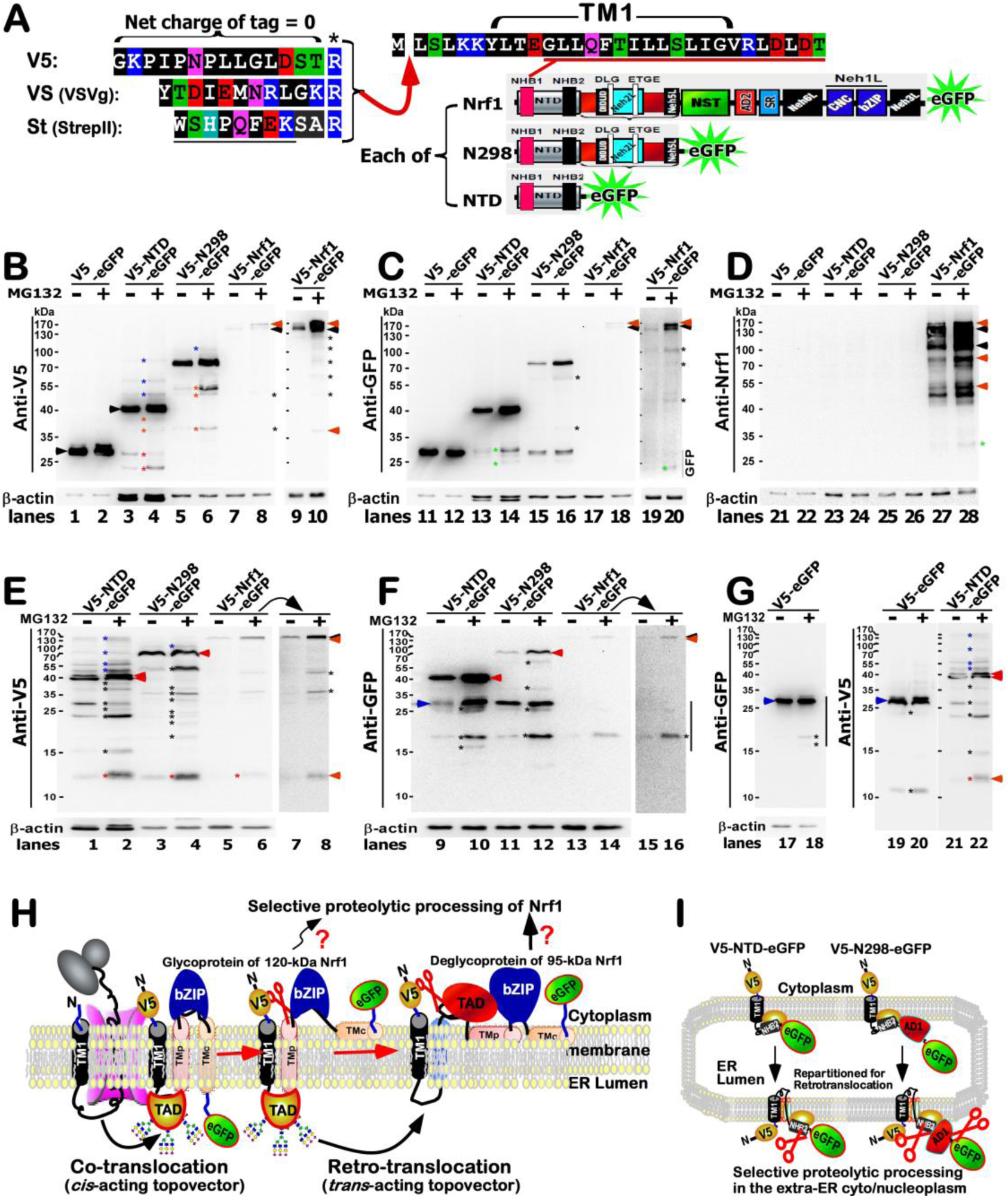
The N-terminal ∼12.5-kDa and other longer polypeptides of Nrf1 arising from the post-synthetic processing of its doubly-tagged chimeric proteins even in the presence of proteasomal inhibitors. (**A**) Schematic of the full-length Nrf1-, its NTD- and N298-fusion proteins with different tags, in which each tag of V5, VSVg and StrepII (with net zero charges) was attached to the N-terminus of Nrf1 or its truncated portions, and their C-terminal ends were further fused with eGFP. (**B** to **D**) The expression constructs for the doubly-tagged fusion proteins were transfected into COS-1 cells and allowed for a 16-h recovery before additional 4-h treatment with MG132 (5 µmol/L), following by SDS-PAGE containing 10% polyacrylamide and Western blotting with distinct antibodies against V5 (***B***), GFP (***C***), Nrf1 (***D***). In addition, the contrasting images of V5-Nrf1-eGFP (*lanes* 9 & 10, 19 & 20) were cropped from its original (*lanes* 7 & 8, 17 & 18). (**E** to **G**) COS-1 cells were transfected with either V5-Nrf1-eGFP or truncated mutants and allowed for a 16-h recovery, before being treated with MG132 (+ 5 µmol/L) or not (−) for 4 h. Total lysates were resolved by SDS-PAGE containing 15% polyacrymide and then visualized by Western blotting with antibodies against V5 (*E*) or GFP (***F***). Additional samples of V5-eGFP (***G***) were also examined in the parallel experiments. (**H**) A model is proposed, to give an explicit explanation of distinct topovectorial processes of Nrf1 (in the context of V5-Nrf1-eGFP) that dynamically moves in and out of the ER, with its post-synthetic processing into distinct isoforms. (**I**) Shows another proposed model of putative proteolytic processing of V5-NTD-eGFP and V5-N298-eGFP.

Further examination of V5-NTD-eGFP (Figure 3,B to D), VSVg-N298-eGFP and StrepII-N298-eGFP (Figure S3) revealed that a major full-length protein of ∼41-kDa, along with additional four minor processed polypeptides of between ∼39-kDa and 23-kDa, was detected by immunoblotting with antibodies against V5 or other tags. The intact ∼41-kDa major protein, plus other two minor processed polypeptides of between 28-kDa and 25-kDa, was also visualized by anti-GFP antibodies (Figure 3C). Similar immunoblotting of V5-N298-eGFP (Figure 3,B to D), along with VSVg-N298-eGFP and StrepII-N298-eGFP (Figure S3), indicated a major ∼80-kDa full-length protein that was recognized coincidently by distinct antibodies against GFP and V5 (or other tags). This was accompanied by additional three minor processed polypeptides of ∼55-, 50- and 38-kDa, that were visualized by their N-terminal tags-specific antibodies (Figure 3B), together with other four minor processed polypeptides of between ∼63-kDa and 25-kDa, that were examined by their C-terminal GFP-recognized antibodies (Figure 3C). Interestingly, a ∼36/37-kDa N-terminally-tagged polypeptide were generated from in all three sandwiched proteins of V5-NTD-eGFP, V5-N298-eGFP and V5-Nrf1-eGFP (Figure 3B). Collectively, these data suggest that putative proteolytic processing of Nrf1-sandwiched proteins to yield several small cleaved polypeptides may occur within its NTD and/or AD1. These two domains of Nrf1 might also target the GFP epitope of fusion proteins to the proteasomal degradation pathway, because abundances of their derived polypeptides were obviously enhanced by treatment with proteasomal inhibitor MG132. In addition, expression of V5-NTD-eGFP and V5-N298-eGFP also gave rise to several forms of higher molecular weight than their original full-length proteins (Figure 3B), implying that they may be ubiquinated through fused NTD and AD1 regions of Nrf1.

Surprisingly, re-examination of V5-NTD-eGFP, V5-N298-eGFP and V5-Nrf1-eGFP by SDS-PAGE containing 15% polyacrylamide, followed by Western blotting with distinct antibodies, clearly revealed the presence of a major N-terminally V5-tagged polypeptide of ∼12.5-kDa (containing most of putative UBL), arising from all three fusion proteins (Figure 3E). However, another N-terminal UBL-containing polypeptide of ∼15-kDa was detected only in the cells expressing V5-NTD-eGFP, but not seen in other two cases of V5-N298-eGFP and V5-Nrf1-eGFP (Figure 3E). It is thus reasoned that the proteolytic processing of Nrf1 by an endopeptidase or endoproteinase leads to removal of its N-terminal polypeptide of ∼12.5-kD and ∼15-kDa from the UBL-adjoining NTD and AD1 regions. In addition, the putative processing of V5-NTD-eGFP gave rise to an N-terminally-tagged polypeptide ladder composed of between ∼140-kDa and 15-kDa, besides its intact full-length ∼41-kDa protein (Figure 3E). This was also accompanied by several C-terminally GFP-tagged polypeptides between 27-kDa and ∼16-kDa (Figure 3F). An additional similar, but different, polypeptide ladder was also observed following separation of total lysates of cells expressing V5-N298-eGFP (Figure 3,E and F), but not V5-eGFP only (Figure 3G). Collectively, it is deduced from these multiple polypeptides arising from the processing of V5-NTD-eGFP and V5-N298-eGFP, that both are allowed for the progressive proteolytic cleavage and/or degradation of within NTD and AD1 (Figure 3,H and I), but this event may also occur within the GFP fused.

Next experiments showed that yield of N-terminally-tagged ∼12.5-kDa (and ∼15-kDa somewhere) polypeptides from V5-NTD-eGFP and V5-N298-eGFP was enhanced by treatment with proteasomal inhibitors MG132, ALLN and bortezomib (BTZ), but not calpain inhibitor-II (CII), calpeptin (CP), ER stressors tunicamycin (TU) or thapsigargin (TG) (Figure 4, A and D). Also, a dose-dependent enhancement in the abundance of the N-terminal ∼12.5-kDa polypeptide was also observed following treatment of cells with MG132 (at 0.5-10 μmol/L) or BTZ (at 0.01-10 μmol/L) (Figure 3,G and J). Even though cells had been treated with higher concentrations of MG132 and BTZ, no significant inhibition of the ∼12.5-kDa polypeptide and other proteoforms was found. Overall, the N-terminal ∼12.5-kDa and longer polypeptides are not only degraded by 26S proteasomes, but may also be yielded from the selective proteolytic cleavage of Nrf1 fusion proteins by other proteasomal inhibitor-insensitive protease-mediated pathways.

**Figure 4.**
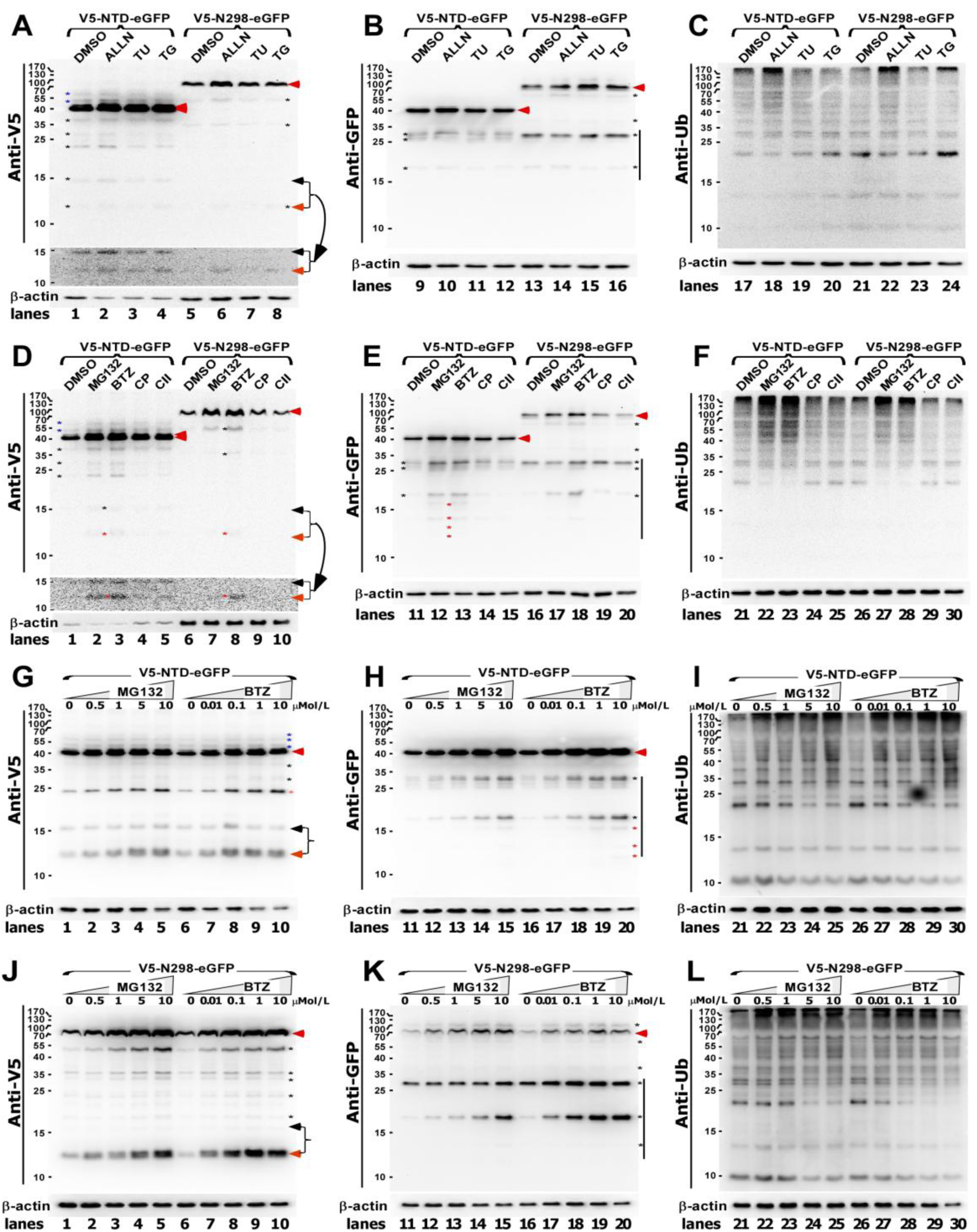
An increase in the N-terminal ∼12.5-kDa polypeptide of Nrf1 arising from the processing of V5-NTD-eGFP and V5-N298-eGFP following treatment with proteasomal inhibitors but not other chemicals used herein. (**A** to **F)** COS-1 cells were transfected with either V5-NTD-eGFP or V5-N298-eGFP and then treated with DMSO, ALLN (5 µg/ml), TU (1 µg/ml), TG (1 µmol/L), MG132 (5 µmol/L), BTZ (1 µmol/L), CP (5 µg/ml) or CII (5 µg/ml) for 4 h, before being harvested in denatured lysis buffer. Total lysates were then subjected to protein isolation by SDS-PAGE containing 15% polyacrymide, followed by immunobloting with antibodies against V5 (***A, D***), GFP (***B, E***) or Ub (***C, F***). **(G** to **I)** Different concentrations of proteasome inhibitors MG132 and BTZ were allowed for 4-h incubation with COS-1 cells that had been transfected with V5-NTD-eGFP or V5-N298-eGFP. The cells were harvested in denatured lysis buffer and protein extracts were examined as described above.

### Distinctive effects of CRAC-adjoining NHB1 and NHB2 peptides on the proteolytic processing of Nrf1 fusion protein to yield its N-terminal ∼12.5-kDa polypeptide

To clarify which peptides within the N-terminal UBL-adjoining regions of Nrf1 are intrinsically involved in putative proteolytic processing of its chimeric proteins to produce the N-terminal 12.5-kDa polypeptide, their mutagenesis mapping were experimented by a series of expression constructs for internal deletion mutants from within NTD and AD1 (Figure 5A). As anticipated, deletion of an NHB2-adjoning peptide from the N298 portion of chimeric protein (to yield a mutant V5-N298^Δ107-124^-eGFP) diminished the N-terminally processed 12.5-kDa polypeptide to a lesser extent than that arising from the intact protein (Figure 5B, *lanes 10 vs 2*, and also Figure S4). By contrast, the N-terminal 12.5-kDa polypeptide was superseded by each of other relative weak proteoforms faster-migrated to: (i) ∼11.5-kDa, arising from either V5-N298^Δ31-50^-eGFP (lacking an NHB1-associated 22-aa sequence, *lane 4*) or V5-N298^Δ55-80^-eGFP (lacking the 26-aa CRAC1/2 peptide, *lane 6*); (ii) ∼11-kDa, yield from V5-N298^Δ80-106^-eGFP (lacking the NHB2 peptide of 27-aa, *lane 8*); and (iii) ∼12-kDa, raised from V5-N298^Δ125-170^-eGFP (lacking the PEST1-adjoning 46-aa sequence of AD1, *lane 12*) (Figure 5B). These data together indicate a potential cleavage site within the C-terminal border of NHB2. Intriguingly, the full-length ∼80-kDa V5-N298-eGFP and major ∼55-kDa processed polypeptide were unaffected by removal of CRAC1/2- or NHB2-adjoning peptides (*lanes 1 to 10*); but both disappeared so as to be replaced by two shorter polypeptides of ∼65-kDa and ∼39-kDa in the case of V5-N298^Δ125-170^-eGFP (*lane 12*). In addition, other N-terminally-derived polypeptides between ∼36-kDa and ∼14-kDa were partially prevented, dependent on deletion mutants of CRAC1/2- or NHB2-adjoning peptides (from the putative UBL), but not the PEST1-adjoning sequence.

**Figure 5.**
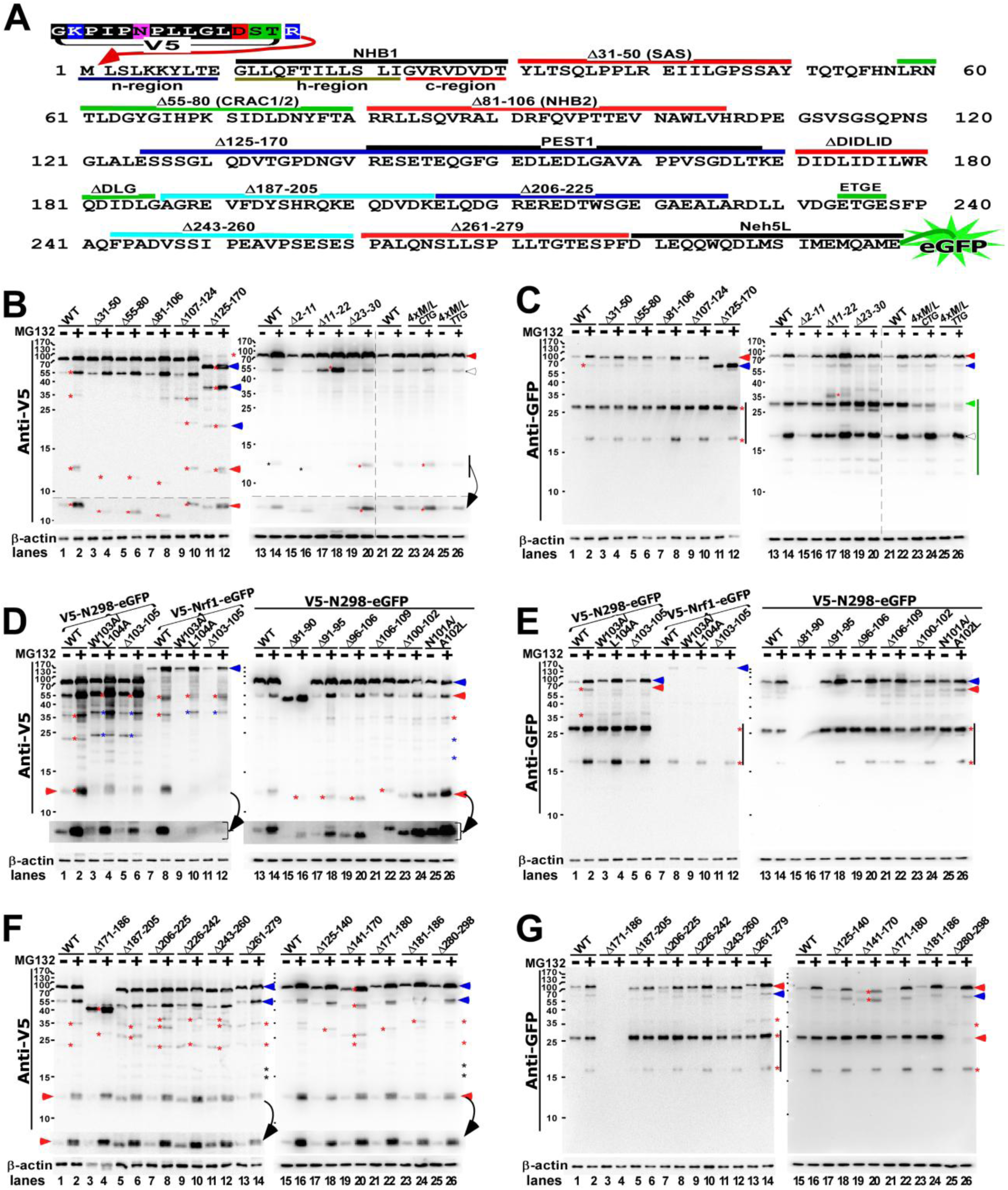
CRAC-adjoining NHB1 and NHB2 peptides of Nrf1 have distinct roles in monitoring the processing of its fusion proteins to yield its N-terminal ∼12.5-kDa and other longer polypeptides. (**A**) Schematic of amino acid residues covering the N298 portion of Nrf1 in different fusion contexts of V5-N298-eGFP and its deletion mutants, with several motifs as indicated. (**B** to **G**) A series of expression constructs for V5-N298-eGFP and its mutants (2 µg of DNA, each) were transfected into COS-1 cells for 7 h and allowed for a 16-h recovery in fresh media before being treated for 4 h with MG132 (+ 5 µmol/L) or not (−), prior to being harvested in denatured lysis buffer. The total lysates were separated by SDS-PAGE containing 15% polyacrylamide and then visualized by Western blotting with indicated antibodies against V5 (***B, D, F***) or GFP (***C, E, G***), as described for the legend of Figure 4.

Next, we examined whether both the NHB1 signal peptide and TM1-adjoining regions have distinct effects on the proteolytic processing of Nrf1 fusion protein to yield the N-terminal ∼12.5-kDa polypeptide. Surprisingly, the N-terminally V5-tagged ∼12.5-kDa polypeptide was almost completely abolished by the mutant V5-N298^Δ11-22^-eGFP (Figure 5B, *lane 18*, which lacks the core h-region of TM1 integrated within membranes). Interestingly, disappearance of the N-terminal ∼12.5-kDa polypeptide seemed to be accompanied by a marked increase in the processed ∼55-kDa polypeptide rather than the ∼80-kDa full-length protein (*cf. lanes 18 with 14*). By further comparison with wild-type protein, deletion mutant of the n-region of ER-targeting peptide (to yield V5-N298^Δ2-10^-eGFP) did not cause an equivalent N-terminal polypeptide to be slightly faster migrated to ∼12-kDa, and also obviously reduced abundances of the ∼12-kDa and ∼55-kDa processed polypeptides, and its intact ∼80-kDa proteins (*lanes 16 vs 14*). Conversely, an apparent increase in abundance of the N-terminal ∼12.5-kDa polypeptide with unaltered mobility was also observed (*lanes 20 vs 14*), upon loss of the c-region immediately to TM1-anchoring peptide (to yield V5-N298^Δ23-30^-eGFP), with no changes in intact ∼80-kDa protein and derived ∼55-kDa polypeptide. Collectively, these data demonstrate that the ER-anchored NHB1 peptide dictates the proteolytic processing of V5-N298-eGFP to yield the N-terminal ∼12.5-kDa and other polypeptide. The process is finely monitored by distinct TM1-adjoining regions, as consistent with the notion that they predominantly determine the proper topology of Nrf1 with correct orientation integrated within and around ER membranes [1, 7, 14, 19].

### Production of the N-terminal ∼12.5-kDa polypeptide from the putative proteolytic processing within the NHB2-adjoining regions of Nrf1 fusion protein

To gain an in-depth insight into which residues within NHB2 control the proteolytic processing of Nrf1 to yield its N-terminal ∼12.5-kDa polypeptide, the putative effects were here determined by mutagenesis mapping of NHB2 to create both short-length deletions and point-mutants from V5-N298-eGFP and V5-Nrf1-eGFP. As anticipated, three short deletions of residues 81-90, 91-95, and 96-106 resulted in an obviously faster mobility of their N-terminally-derived polypeptides to be located at ∼12-kDa during electrophoresis (Figure 5D, *cf. lanes 15-20 with 14*). Both basal abundance and MG132-stimulated accumulation of the ∼12-kDa mutant polypeptides were, to a greater or lesser extent, diminished by V5-N298^Δ81-90^-eGFP, V5-N298^Δ91-95^-eGFP and V5-N298^Δ96-106^-eGFP. Similar faster mobility of the N-terminal ∼12-kDa mutant polypeptide was, though its abundance was significantly increased, displayed upon deletion of just three residues 100-102 (to yield V5-N298^Δ100-102^-eGFP; Figure 5D, *lanes 24 vs 14*) or point mutation (to yield V5-N298^N 101A/A102L^-eGFP; *cf. lanes 26 with 14*). However, no changes in the electrophoretic location of the N-terminal ∼12.5-kDa polypeptide were observed in the case of V5-N298^Δ106-109^-eGFP (lacking only four residues on the C-terminal border of NHB2, *lane 22*), albeit its abundance was reduced by this deletion mutant. Amongst the above-described mutants, only V5-N298^Δ81-90^-eGFP exhibited an unusual pattern of fewer electrophoretic bands (Figure 5,D and E, *lanes 15,16*). Of note, disappearance of the ∼80-kDa full-length V5-N298^Δ81-90^-eGFP, its N-terminally-derived ∼55-kDa and 12.5-kDa polypeptides, along with all its C-terminally GFP-tagged forms was accompanied by the emergence of additional two N-terminal polypeptides of ∼50-kDa (strongly) and ∼12-kDa (very weakly).

The above-described data indicate that distinct NHB2-adjoining peptides of Nrf1 may make differential and even opposing contributions to either its protein stabilization or its proteolytic processing to generate the N-terminal ∼12.5-kDa polypeptide. To confirm this hypothesis, we examined four additional deletion mutants immediately on the C-terminal border of NHB2 from V5-Nrf1-eGFP and V5-N298-eGFP. The results showed that the N-terminal ∼12.5-kDa polypeptide arising from the processing of N298 was significantly diminished, but not abolished, by two mutants V5-N298^Δ103-105^-eGFP and V5-N298^W103A/L104A^-eGFP (Figure 5D*, lanes 3-6 vs 2*). However, similar N-terminal ∼12.5-kDa polypeptide arising from the full-length Nrf1-fusion protein appeared to be almost completely abolished by V5-Nrf1^Δ103-105^-eGFP and V5-Nrf1^W103A/L104A^-eGFP (*cf. lanes 9-12 with 8*). Unexpectedly, such diminishment and even abolishment of the N-terminally-derived 12.5-kDa polypeptide was not accompanied by additional changes in the electrophoretic mobility of the full-length proteins and their abundances, but with an exception of three major processed polypeptides migrated slowly to between 25-kDa and 55-kDa (*lanes 3-6 vs 2*). Such slower mobility of other two major processed polypeptide of between 35-kDa and 55-kDa was also observed after electrophoresis of V5-Nrf1^Δ103-105^-eGFP and V5-Nrf1^W103A/L104A^-eGFP. Collectively, it is inferable that Nrf1- and its N298-fusion proteins are dynamically targeted for selective progressive proteolytic processing of within its NHB2-adjoining regions, in order to yield various lengths of its N-terminally (and C-terminally)-derived polypeptides.

Notably, no changes in the electrophoretic motility of the N-terminally-derived 12.5-kDa polypeptide of Nrf1 were found following various internal deletions of between its residues 125-298 from AD1 of V5-N298-eGFP (Figure 5F). However, those short-lengthened mutants of between residues 141-260 resulted in obvious variations of the mobility of both the ∼80-kDa full-length protein and its processed ∼55-kDa polypeptide (Figure 5,F and G, *lanes 3-12, 19, and 20*). Intriguingly, no similar protein bands close to ∼80-kDa and ∼55-kDa were resolved by electrophoresis of cell lysates transfected with V5-N298^Δ171-186^-eGFP (lacking the DIDLID/DLG element within Neh2L degron of AD1) (Figure 5, F and G, *lanes 3, 4*), but the appearance was accompanied by the presence of other two closer ∼40/50-kDa polypeptides. This should be owing to a loss of the C-terminally-tagged GFP from the V5-N298^Δ171-186^-eGFP fusion protein, which may be targeted for the non-proteasomal proteolytic degradation to disappear (Figure 5G, *lanes 3, 4*). Intriguingly, such sharp changes in these two cases of V5-N298^Δ171-186^-eGFP and V5-N298^Δ125-170^-eGFP (Figure 5,B and C, *lane 12*) were not recurred in additional cases with shorter deletions of between residues 125-186 (Figure 5, F and G, *lanes 17-24*), albeit abundance of the N-terminal 12.5-kDa polypeptide was decreased to a greater or lesser extent. Furthermore, generation of the N-terminal 12.5-kDa polypeptide was decreased by deletion mutants of within Neh5L-adjoining residues 243-298 (*lanes 14, 26*), but slightly increased by other mutants of within residues 187-242 within Neh2L degon (*lanes 6 to 12*). Together with our previous data [11, 19], these results indicate that the PEST1-adjoining DIDLID/DLG element and Neh5L, but not Neh2L, within AD1 are also required for the protein stabilization and/or proper topological folding.

Moreover, to exclude an effect of potential in-frame translation of Nrf1 on its post-synthetic processing, we created two Neh5L-containing mutants of possible internal translation start *ATG* codons (at methionines 289, 292, 294 and 297) into two different leucine codons, to yield V5-N298^4xM/L(*CTG*)^-eGFP and V5-N298^4xM/L(*TTG*)^-eGFP (Figure 5, A and B). The results showed that abundances of both N-terminally-derived ∼12.5-kDa and ∼55-kDa polypeptides were not decreased but marginally increased (Figure 5B, *lanes 23-26*). However, the C-terminal GFP-containing ∼27-kDa, but not its processed ∼18-kDa, polypeptides were significantly prevented and even completely abolished by these two mutants V5-N298^4xM/L(*CTG*)^-eGFP and V5-N298^4xM/L(*TTG*)^-eGFP, respectively (Figure 5C, *lanes 23-26*). Overall, this suggests the putative proteolytic processing of V5-N298-eGFP to yield its N-terminal polypeptides may also be monitored by Neh5L-adjoining residues.

### A role of p97 for the retrotranslocation and otherwise in selective proteolytic processing of Nrf1-fusion protein to remove its N-terminal ∼12.5-kDa and other polypeptides

As shown in Figure 6, inhibition of p97-driven retrotranslocation by NMS-873 (at 1∼10 μmol/L) resulted in a dose-dependent increase in abundances of the intact ∼80-kDa V5-N298-eGFP and its derived N-terminal polypeptides of ∼12.5-kDa and 55-kDa (Figure 6A, *lanes 5-8*). In the meantime, no obvious changes in the C-terminal ∼18-kDa and ∼27-kDa polypeptides arising from V5-N298-eGFP were determined following treatment of the p97 inhibitor (Figure 6B, *lanes 5-8*). By comparison with V5-N298-eGFP, no striking alterations in the full-length ∼41-kDa V5-NTD-eGFP and its processed N-terminal polypeptides of ∼12.5-kDa, but not ∼23-kDa, were examined following treatment of cells with p97 inhibitor NMS-873 at same doses as described above (*lanes 1 to 4*). These suggest that a dominant effect of p97 on the retrotranslocation of V5-N298-eGFP, distinctive from V5-NTD-eGFP, may be a premise for selective proteolytic processing of N298, rather than NTD, in the fusion contexts.

**Figure 6.**
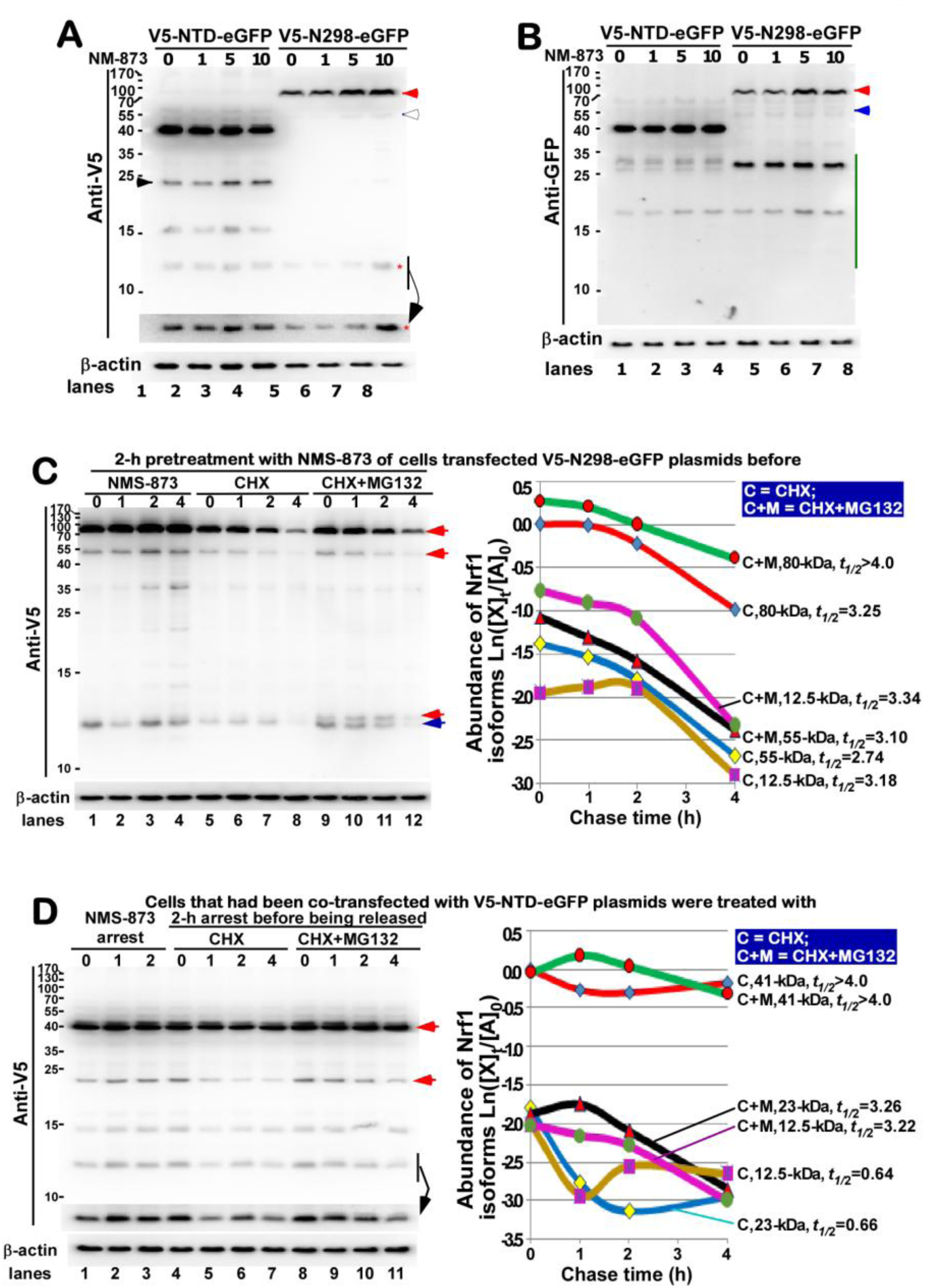
The proteolytic processing of Nrf1-fusion proteins to yield its N-terminal ∼12.5-kDa and other polypeptides is determined by p97-driven retrotranslocation and -independent pathways. (**A, B**) Different concentrations (0-10 µmol/L) of p97-specific inhibitor NMS-873 were incubated for 4 h with COS-1 cells that had been transfected with either V5-NTD-eGFP or V5-N298-eGFP, followed by Western blotting with distinct antibodies against V5 (*A*) or GFP (*B*). (**C, D**) COS-1 cells expressing V5-N298-eGFP (*C*) or V5-NTD-eGFP (*D*) were treated with either NMS-873 (10 µmol/L), CHX (50 µg/ml) alone or plus MG132 (5 µmol/L) for indicated lengths of time, before being harvested. Equal amounts of proteins in total lysates were separated by SDS-PAGE containing 15% polyacrylamide, and visualized by Western blotting with V5 antibody. β-actin served as the protein-loading control. The intensity of three major polypeptide bands *arrowed* on each gels was quantified and shown graphically in the *right panels*, with distinct half-lives estimated.

To clarify distinct roles of p97 for the retrotranslocation of V5-N298-eGFP and V5-NTD-eGFP in their proteolytic processing by cytosolic proteasomes, herein we performed the pulse-chase experiments of cells that had been pretreated with NMS-873, before addition of 50 μg/ml cycloheximide (CHX, which inhibits biosynthesis of nascent proteins) alone or plus MG132. As expected, NMS-873 inhibition of p97-mediated repositioning of V5-N298-eGFP into extra-ER compartments led to an incremental accumulation of the fusion protein and its derivate polypeptides between 80-kDa and 12.5-kDa (Figure 6C, *lanes 1-4*). Subsequently, the recovery of p97 from its inhibitor, at the same time when newly-synthesized proteins were blocked by CHX, rendered portion of the existing ER-resident protein V5-N298-eGFP to be dynamically dislocated into the extra-luminal cytoplasmic side of membranes, in which it was allowed for the successive proteolytic digestion by 26S proteasomes and/or cleavage by other proteases through potential degrons and cleavage sites within N298. Just so a recovery of p97 from its inhibition by NMS-873 led to the release of the existing V5-N298-eGFP, subsequent turnover of its full-length ∼80-kDa protein, together with its derived N-terminal polypeptides of ∼55-kDa and ∼12.5-kDa (Figure 6C, *lanes 5-8*), was determined with their distinct half-lives estimated to be 3.25, 2.74 and 3.18 h after treatment with CHX, respectively (*right graphic*). By contrast, addition of MG132 (at 5 μmol/L) differentially prolonged the half-lives of the ∼80-kDa V5-N298-eGFP, as well as its N-terminal ∼55-kDa and ∼12.5-kDa polypeptides to >4.0, 3.10 and 3.34 h following CHX treatment, respectively (Figure 6C, *lanes 9-12*, *left graphic*). More curiously, the yield of two closer polypeptides of ∼12.5-kDa was augmented by this proteasomal inhibitor (*double arrows*). These data, together with the results for the ‘bounce-back’ response to proteasome-limited inhibition [16, 18, 20], demonstrate that the partial proteolytic processing of V5-N298-eGFP by 26S proteasomes gives rise to its N-terminal polypeptides of particularly ∼12.5-kDa.

By comparison with V5-N298-eGFP, the 41-kDa full-length V5-NTD-eGFP was slightly arrested by NMS-873 (Figure 6D, *lanes 1-3*), but its stability was almost unaffected by the release of p97 from its inhibitor or addition of the proteasomal inhibitor MG132 (*lanes 4-11 vs 1-3, right graphic*). However, it is interesting that an apparent arrest of the N-terminal ∼23-kDa and ∼12.5-kDa isofoms from V5-NTD-eGFP was made by pretreatment of NMS-873, but an increase in their turnover occurred after release of p97 inhibitor (Figure 6D, *lanes 4-7 vs 1-3*). The increased turnover of the ∼23-kDa and ∼12.5-kDa polypeptides was determined with similar shorter half-lives calculated to be 0.66 and 0.64 h after CHX treatment, respectively (*right graph*). Conversely, abundances of the N-terminal ∼23-kDa and ∼12.5-kDa polypeptides of V5-NTD-eGFP were markedly promoted by MG132, such that their half-lives were also significantly prolonged to 3.26 and 3.22 h after CHX treatment, respectively (*lanes 8-11, right graph*). Collectively, it is inferable that distinctions in dynamic repartitioning and repositioning of V5-NTD-eGFP from V5-N298-eGFP, as well as their processed polypeptides, around and within membranes may be attributable to the presence of AD1 or its absence, beyond both TM1-associated NHB1 peptide (anchored within membranes) and CRAC-adjoining NHB2 peptide (anchored in the ER lumen or flipped across membranes).

### Distinct effects of the p97-driven retrotranslocation inhibitor on selective proteolytic processing of between V5-N298-eGFP and V5-NTD-eGFP by cytosolic DDI-1 protease and/or proteasomes

Upon repositioning of the luminal-resident portion of Nrf1-fusion protein into the extra-ER side of membranes, it becomes susceptible to selective proteolytic cleavage and degradation mediated mediated by cytosolic proteases [18, 24, 25]. Since no blockage of Nrf1-derived ∼12.5-kDa N-terminal polypeptide by proteasomal inhibitors (Figure4), we next determined whether DNA damage-inducible protein-1 and -2 (i.e. DDI-1 and DDI-2) have an ability to act as functional proteases involved in selective proteolytic processing of Nrf1. The results showed that abundance of the N-terminal ∼12.5-kDa polypeptide arising from V5-N298-eGFP was substantially increased by co-expression with DDI-1^3xFlag^ or DDI-2^3xFlag^ (Figure 7A, *lanes 2,6*). The ∼12.5-kDa polypeptide was further augmented by MG132 (*lanes 4,6*), despite no promotion of DDI-1 and DDI-2 (*middle panels*). By contrast, no remarkable changes in the ∼80-kDa full-length V5-N298-eGFP and processed ∼55-kDa polypeptide were observed. Conversely, knockdown of DDI-1 and DDI-2 by siRNA-targeting (Figure 7B) caused an obvious decrease in yield of the Nrf1-derived ∼12.5-kDa polypeptide (Figure 7C). Further effective inhibition of the polypeptide resulted from double knockdown of DDI-1 and DDI-2 (*lane 4*), implying an overlapping function of both DDI-1 and DDI-2 proteases in the proteolytic cleavage of Nrf1. In addition, yield of the N-terminal polypeptides was not completely abolished by double knockdown of DDI-1 and DDI-2, indicating that other proteases may also be involved in the proteolytic processing of Nrf1.

**Figure 7.**
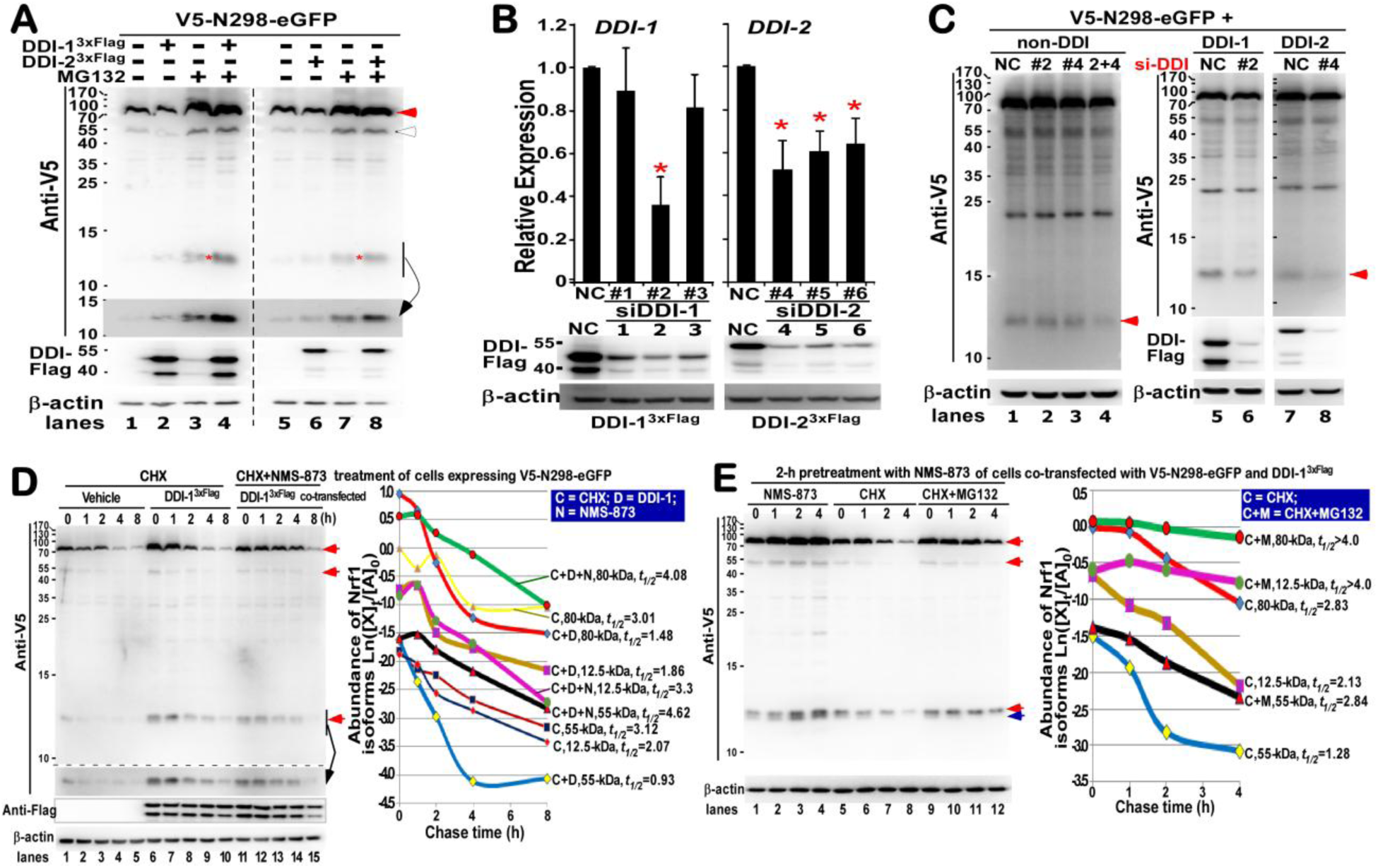
A role of p97 for the retrotranslocation of Nrf1 in selective proteolytic processing of the client proteins by cytosolic DDI-1/2 protease and proteasomes to remove its N-terminal polypeptides. (**A**) COS-1 cells that had been co-transfected with an expression construct for V5-N298-eGFP (1 µg of DNA) alone or in a combination with either DDI-1^3xFlag^ or DDI-2^3xFlag^ (1 µg of DNA) were treated with MG132 (+ 5 µmol/L) or not (−) for 4 hbefore being harvested in denatured lysis buffer. The total lysates were then examined by Western blotting with distinct antibodies against V5 or Flag. (**B**) The interference efficiency of siRNAs against *DDI-1* or *DDI-2* was determined as shown graphically. HEK293 cells were co-transfected with an expression construct for DDI-1^3xFlag^ or DDI-2^3xFlag^ with specific siRNAs (Table S1), before being examinated by RT-PCT and Western blotting with Flag antibody. (**C**) The expression plasmid for V5-N298-eGFP plus DDI-1^3xFlag^ or DDI-2^3xFlag^, together with siRNAs were co-transfected into COS-1 cells. The lysates were analyzed by SDS-PAGE containing 15% polyacrylamide and immunoblotting with V5 or Flag antibodies. (**D,E**) COS-1 cells expressing V5-N298-eGFP alone or plus DDI-1^3xFlag^ were treated with either NMS-873 (10 µmol/L), CHX (50 µg/ml) alone or plus MG132 (5 µmol/L) for indicated lengths of time, before being harvested. Equal amounts of proteins in total lysates were separated by SDS-PAGE containing 15% polyacrylamide, and visualized by Western blotting with V5 antibody. DDI-1 was identified by its Flag antibody (***A*** to ***D***), whilst β-actin served as the protein-loading control. The intensity of three major polypeptide bands *arrowed* on each gels was quantified as shown graphically in the *right panels*, with distinct *t_1/2_* values estimated.

To determined whether the selective proteolytic processing of V5-N298-eGFP by cytosolic DDI-1 protease was monitored by p97-driven retrotranslocation or -independent pathways. the pulse chase experiments were carried out. Western blotting showed a time-dependent decrease in the existing intact ∼80-kDa V5-N298-eGFP, along with its N-terminally-derived polypeptides of ∼55-kDa and ∼12.5-kDa, following only CHX treatment of cells transfected without DDI-1 (Figure 7D, *lanes 1-5*). Their protein turnover was determined with distinct half-lives estimated to be ∼3.01, 3.12 or 2.07 h after CHX treatment, respectively (*right graph*). Intriguingly, co-expression of DDI-1^3xFlag^ led to apparent increases in abundances of the intact ∼80-kDa V5-N298-eGFP and its processed ∼55-kDa and ∼12.5-kDa polypeptides (*lanes 6-10*). As such, their turnover was accelerated by this protease, because their half-lives were significantly shortened to 1.48, 0.93 or 1.86 h after CHX treatment, respectively (*right graph*). By sharp contrast, the DDI-mediated proteolytic turnover of V5-N298-eGFP and its two N-terminal polypeptides was markedly alleviated by NMS-873 (*lanes 11-15*), leading to the prolongation of their half-lives to 4.08, 4.62 or 3.30 h after co-treatment with CHX, respectively (*right graph*). This finding demonstrates that rapid proteolytic destruction of V5-N298-eGFP and its derived polypeptides by DDI-1 is significantly prevented, but not completely abolished, by inhibition of p97-driven retrotranslocation into the extra-ER side of membranes.

In order to elucidate whether DDI-1 alone or plus proteasomes mediate the selective proteolytic processing of V5-N298-eGFP released by p97-trigged retrotranslocation into the extra-ER cyto/nucleoplasmic side, experimental cells had been allowed for co-expression of this protease before being treated as indicated chemicals (Figure 7E). The results revealed that NMS-873-arrested accumulation of V5-N298-eGFP and its derivate polypeptides was almost unaltered by co-expression of DDI-1 (*lanes 1-4*). This finding suggests that dynamic repositioning of V5-N298-eGFP is blocked by the p97 inhibitor, such that it is tempo-spatially sequestered from the DDI-1 protease on two distinct sides of ER membranes. After recovery of p97 from its inhibition by NMS-873, the existing ∼80-kDa V5-N298-eGFP was released into the cyto/nucleoplasmic side, whereupon it was allowed for rapid proteolytic processing by DDI-1 alone or plus other proteases (*lanes 5-8*). The resulting half-life of ∼80-kDa V5-N298-eGFP was shortened to 2.83 h following CHX treatment (*right graph*), as compared with the absence of DDI-1 (as described in Figure 6C, with *t_1/2_* =3.28 h). Conversely, the stability of intact V5-N298-eGFP was increased, with a prolonged half-life, by addition of MG132, even in the presence of DDI-1 (Figure 7E, *lanes 9-12*).

It is rather peculiar and unique that, even though most of the full-length ∼80-kDa V5-N298-eGFP was arrested by p97 inhibitor and hence accumulated on the ER luminal side of membranes, this was still accompanied by a time-dependent increase in its N-terminal polypeptides of ∼55-kDa and ∼12.5-kDa (Figure 7E, *lanes 1-4*). This implies that some potential cleavage sites of Nrf1 may be partially repositioned through p97-independent pathways into the extra-luminal side, in which they could contribute as a local target for the juxtamembrane proteolysis by DDI-1 and other proteases, in order to yield several membrane-protected N-terminal polypeptides (as proposed in Figure3 H & I). Once released, the N-terminal polypeptides were dislocated by p97-driven retrotranslocation, enabling for their destruction mediated by DDI-1. Thus, the half-lives of both the ∼55-kDa and ∼12.5-kDa polypeptides were shortened to 1.28 and 2.13 h after CHX treatment, respectively (Figure 7E, *lanes 5-8, right graph*), as compared with those in the absence of DDI-1 (as described in Figure 6C*, with t_1/2_* = 2.74 and 3.18 h, respectively). However, a marked enhancement in stability of the ∼55-kDa and ∼12.5-kDa polypeptides of V5-N298-eGFP by MG132 resulted in an extension of both half-lives to 2.84 h and more than 4.0 h, respectively (Figure 7E, *lanes 9-12, right graph*).

By comparison with V5-N298-eGFP, additional parallel experiments of V5-NTD-eGFP (Figure S5) revealed that its full-length ∼41-kDa protein was modestly accumulated by NMS-873 in the presence of DDI-1, but its stability was almost unaffected by co-expression of DDI-1, and also unaltered even following release of p97 from its inhibitor NMS-873 or addition of proteasomal inhibitor MG132. The half-life of 41-kDa V5-NTD-eGFP (holding an unchanged basal abundance within 4 h) was estimated to be over 8 h following treatment with CHX (*right graphs*). By contrast, its two N-terminal polypeptides of ∼23-kDa and ∼12.5-kDa were accumulated by arrest of p97 inhibitor (Figure S5B, *lanes 1-3*). However, co-expression of DDI-1 caused an obvious reduction in the ∼23-kDa abundance, even in the treatment with NMS-873 (Figure S5A, *lanes 6-15*). As such, the turnover of the ∼23-kDa polypeptide (derived from the ∼41-kDa V5-NTD-eGFP) was slightly mitigated by DDI-1 alone or plus the p97 inhibitor, which was determined with a half-life extended from 1.40 h to 5.20 or 7.64 h, respectively (*right graph*). Rather, co-expression of DDI-1 led to a significant promotion of the N-terminal ∼12.5-kDa polypeptide arising from the processing of intact V5-NTD-eGFP and its derived ∼23-kDa polypeptide, even in the presence of NMS-873 (Figure S5A). The stability of the ∼12.5-kDa polypeptide was thus enhanced, with a half-life prolonged from 1.06 h to over 8.0 h following treatment with CHX (*right graph*). All together, these data demonstrate that the presence of AD1 within V5-N298-eGFP facilitates its retrotranslocation by p97-fueled pathway into extra-ER compartments, allowing for the proteolytic processing of Nrf1-fusion protein by DDI-1 and proteasomes to remove its N-terminal polypeptides. Conversely, the absence of AD1 from V5-NTD-eGFP enables it to be mostly retained in the ER lumen and hence protected by membranes. In addition, a small fraction of V5-NTD-eGFP and its N-terminal polypeptides (retaining the CRAC motif that may flip across the cholesterol-rich membranes) is hence postulated to gain an accessibility to the extra-ER juxtamembrane proteolysis mediated by cytosolic proteases.

### Putative ubiquitination of Nrf1 is not a prerequisite necessary for involvement of p97 in the selective proteolytic processing of the client protein by cytosolic DDI-1 protease and/or proteasomes

A previous report showed that ubiquitination of Nrf1 occurred prior to, and was also essential for, its proteolytic processing to yield several cleaved polypeptides [18]. To test this, all six lysines (as potential ubiquitin acceptors) of Nrf1 within NTD (containing three lysines at positions 5, 6 and 70) and AD1 (containing other three lysines at positions 169, 199 and 205) were here mutated into another basic arginines, so as to create V5-N298^6×K/R^-eGFP (Figure 8A). In fact, the resultant K/R mutant did not appear to interfere with DDI-mediated proteolytic processing of the protein to generate an equivalent to the Nrf1-derived ∼12.5-kDa polypeptide (*middle panels*), but its abundance was markedly enhanced by MG132 (*lanes 4,8*). However, no marked changes in abundances of both intact ∼80-kDa V5-N298^6×K/R^-eGFP and its N-terminal ∼55-kDa polypeptide resulted from co-expression of DDI-1/2.

**Figure 8.**
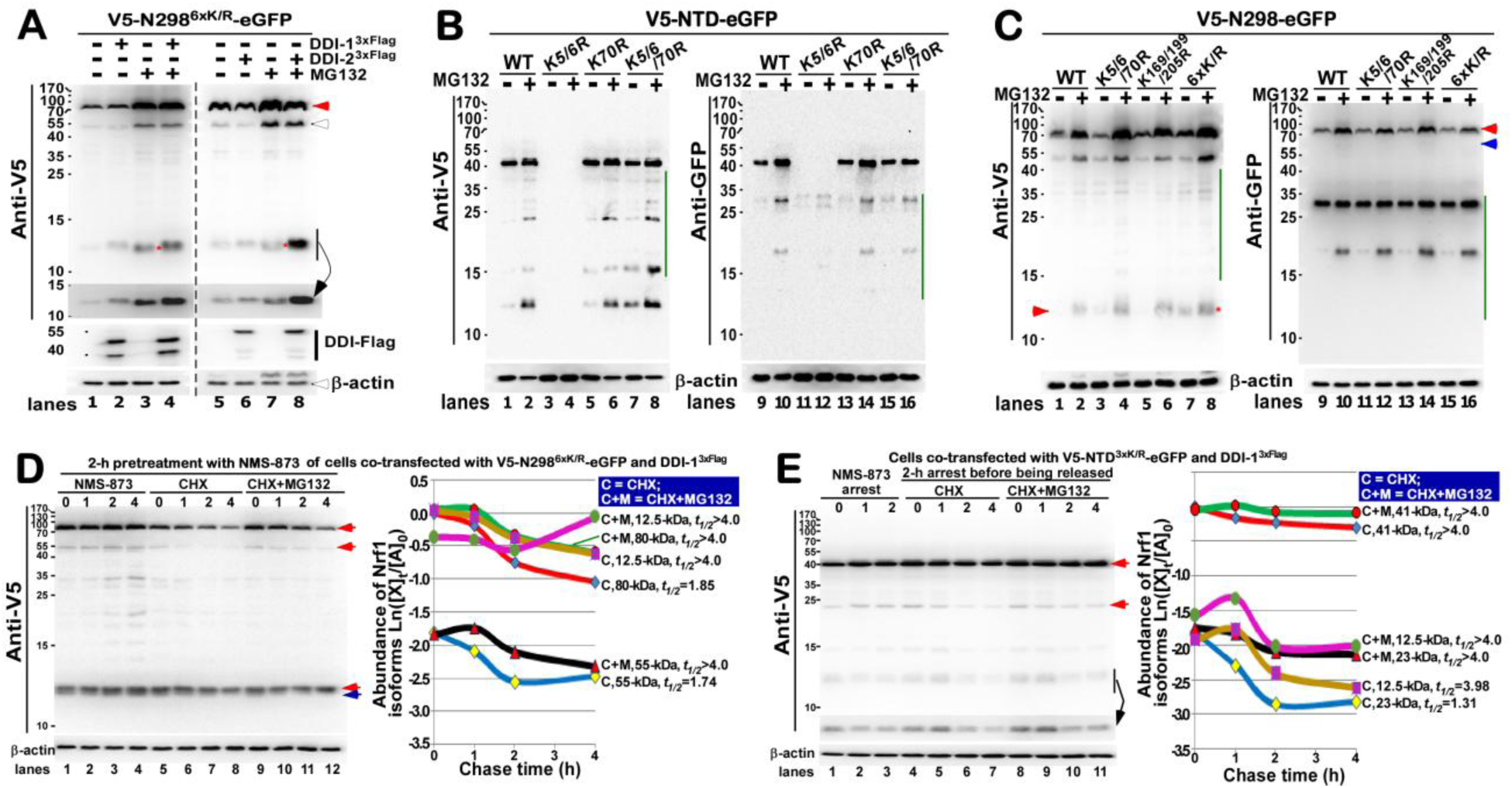
No effects of all putative lysine-based ubiquitination mutants of Nrf1 on the involvement of p97 in the selective proteolytic processing of the client protein by cytosolic DDI-1 protease and proteasomes. (**A**) COS-1 cells expressing V5-N298^6xK/R^-eGFP together with either DDI-1^3xFlag^ or DDI-2^3xFlag^ were treated with MG132 (+ 5 µmol/L) or not (−) for 4 h before being harvested in denatured lysis buffer, followed by Western blotting with V5 or Flag antibodies. (**B,C**) A series of expression constructs for single, double or triple lysine-to-arginine (K/R) mutants of the NTD or N298 portion of Nrf1 attached to eGFP were transfected into COS-1 cells. They were then treated MG132 (+ 5 µmol/L) or not (−) and examined by Western blotting as described above. (**D**, **E**) COS-1 cells were transfected with expression constructs for V5-N298^6×K/R^-eGFP (***D***) or V5-NTD^3×K/R^-eGFP (***E***) in combination with DDI-1^3xFlag^. They were then treated with NMS-873 (10 µmol/L), CHX (50 µg/ml) alone or plus MG132 (5 µmol/L) for indicated lengths of time, before being harvested. Equal amounts of proteins in total lysates were separated by SDS-PAGE containing 15% polyacrylamide, and visualized by Western blotting with V5 antibody. β-actin served as the protein-loading control. The intensity of three major polypeptide bands *arrowed* on each gels was quantified as shown graphically in the *right panels*, with distinct *t_1/2_* values estimated.

Further examinations revealed that generation of the N-terminal ∼12.5-kDa polypeptide from NTD- and N298-fused proteins was not completely abolished, *de facto*, no matter what results were obtained by mapping possibly ubiquitination-based lysine-to-arginine (K/R) mutagenesis of NTD and AD1 (Figures 8B,C and S6). Amongst these mutants, V5-NTD^K5/6/70R^-eGFP caused an obvious increase in abundance of NTD-derived ∼12.5-kDa polypeptide (Figure 8B, *lanes 7,8*). However, almost none of major polypeptides were detected in the case of V5-NTD^K5/6R^-eGFP (*lanes 3,4*); this mutant protein might be unstable, such that it was rapidly degraded to disappear, even in the presence of MG132 (*lanes 11,12*). The notion is also supported by additional putative degradation of all the eGFP-free cases of V5-NTD and mutants (including V5-NTD^K5/6R^) into one major NTD-derived ∼14-kDa and another minor ∼12.5-kDa polypeptides, the latter of which was too faint to be seen (Figure S6A, *lanes 11-18*). By sharp contrast, additional four eGFP-free cases of the V5-N298 and its K/R-indicated mutants gave rise to the full-length protein of ∼45-kDa, along with its N-terminal ∼12.5-kDa and ∼15-kDa polypeptides, the latter two of which seemed similar to those arising from V5-N298-eGFP (Figure S6B, *cf. lanes 11-18 with 10*). Moreover, the proteolytic processing of the intact ∼80-kDa V5-N298-eGFP to removal several polypeptides of between ∼55-kDa and ∼12.5-kDa was not prevented by its K/R mutants (Figure 8C), but yield of these two polypeptides was conversely highlighted, in particular, following MG132 treatment (*lanes 3-8*). Overall, besides NTD, the AD1 region is thus deducible to be required for the proper folding of Nrf1 and its selective proteolytic processing to remove its N-terminal polypeptides, but this event is unaffected by putative ubiquitination of Nrf1. This conclusion is further supported by additional data obtained from a series of single or double K/R mutants and other K/R mutants retaining equivalent lysines in wild-type form (Figure S6, C to F).

Next, we determined whether putative lysine-based ubiquitination of Nrf1 is not a prerequisite for involvement of p97-driven retrotranslocation in selective proteolytic processing of the CNC-bZIP protein by cytosolic proteases. As anticipated, a time-dependent increase in the intact 80-kDa V5-N298^6×K/R^-eGFP, its N-terminal polypeptides of between 55-kDa and 12.5-kDa resulted from arrest of p97-driven retrotranslocation by NMS-873 (Figure 8D, *lanes 1-4*). Subsequently, the recovery of p97 from its inhibitor led to partial repositioning of the ubiquitination-deficient mutant protein and its N-terminal derivates into extra-ER luminal side of membranes, enabling them to gain access to the juxtamembrane proteolytic processing by DDI-1. It just so occurred that the proteolytic processing of the existing V5-N298^6×K/R^-eGFP was accelerated by co-expression of DDI-1 (Figure 8D*, lanes 5-8*). The rapid turnover of its 80-kDa mutant protein was determined with its half-life shortened to be 1.85 h after CHX treatment (Figure 8D, *right graph*), when compared with half-life (*t_1/2_*=2.83 h) of wild-type V5-N298-eGFP (Figure 7E). However, the proteolytic effect of DDI-1 on the processing of V5-N298^6×K/R^-eGFP was partially mitigated by MG132, such that its half-life was prolonged to over 4 h after treatment with CHX (Figure 8E, *lanes 9-12*). More excitingly, a substantial increase in the N-terminally-derived ∼12.5-kDa polypeptide with a half-life prolonged (from 2.13 to over 4.0 h) resulted from co-expression of DDI-1 and V5-N298^6×K/R^-eGFP, but was unaffected by MG132 (Figure 8D, *lanes 5-12*). These indicate that the V5-N298^6×K/R^-eGFP mutant may facilitate its full-length protein to be liberated for the DDI-1-mediated proteolytic processing to remove its N-terminal ∼12.5-kDa and other polypeptides.

Further examinations of V5-NTD^3×K/R^-eGFP revealed that NMS-873 caused a modest increase in the ∼41-kDa full-length protein and N-terminal ∼23-kDa rather than ∼12.5-kDa polypeptides in cells co-expressing DDI-1 (Figure 8E, *lanes 1-3*). After release of p97 from its inhibitor, the existing V5-NTD^3×K/R^-eGFP stability remained unaltered by the presence of DDI-1 alone or plus MG132 (*lanes 4-11*). In addition, the stability of the ∼12.5-kDa, but not ∼23-kD, polypeptides arising from V5-NTD^3×K/R^-eGFP was enhanced by K/R mutant, with a half-life extended from 1.74 h to 3.98 h (Figure 8E), as compared with those of wild-type V5-NTD-eGFP (Figure S5B). Collectively, these demonstrate that distinct effects of p97-driven retrotranslocation and -independent pathways on selective proteolytic processing of Nrf1 fusion proteins by cytosolic proteases are not influenced by the client ubiquitination.

### Topovectorial regulation of Nrf1-fused Gal4D-VP16 transactivation activity and its proteolytic processing

To determine whether the N-terminal proteolytic processing of Nrf1 and its AD1-mediated transactivation activity are monitored by its dynamic membrane-topology within the ER, here we created a series of expression constructs for various lengths of its N-terminal residues 1-298, each of which was attached to the N-terminus of Gal4D-VP16 fusion protein with a C-terminal V5 tag. Theoretically, these resulting fusion proteins were allowed to be anchored within membranes through the TM1-adjoining NHB1 peptide of Nrf1. Subsequently, the CRAC1/2-adjoining NHB2 peptides and AD1 regions, as well as the C-terminally fused Gal4D-VP16 portion, should be co-translocationally positioned on the ER lumen (Figure 9A, *upper cartons*). Only when repositioning of the fused Gal4D-VP16 portion from the luminal side of membranes into the cyto/nuclelasmic side, it may be allowed for transactivation of the Gal4/*UAS*-driven reporter (*lower cartons*). The Gal4-target gene activity may also be promoted if the N-terminal proteolytic processing of Nrf1 by cytosolic proteases renders its fused Gal4D-VP16 to be released from membrane-associated confinements. As anticipated, a membrane-free Gal4D-VP16 factor exhibited a maximum activity of the *UAS-luc* reporter gene to ∼5,975-fold changes (Figure 9C). By contrast, attachment of Gal4D-VP16 to the C-terminus of N65 (i.e. the N-terminal 65-aa of Nrf1) conferred on the resultant N65-Gal4D-VP16 factor with only ∼1,650-fold transactivation activity. Immunoblotting of this C-terminally V5-tagged fusion protein showed a single full-length protein of ∼47-kDa, without any proteolytic polypeptides visualized (Figures 9D and S7A, *lane 2*). Thus, it is inferable that N65-Gal4D-VP16 may act as a membrane-anchored protein entailed with dynamic topologies. This is based on the fact that the N65 (as a membrane-associated half of putative UBL) is protected by membranes against cytosolic protease-mediated proteolysis, and hence its fused Gal4D-VP16 may be restricted in proximity to the membranes, albeit it could also be dislocated into extra-ER compartments.

**Figure 9.**
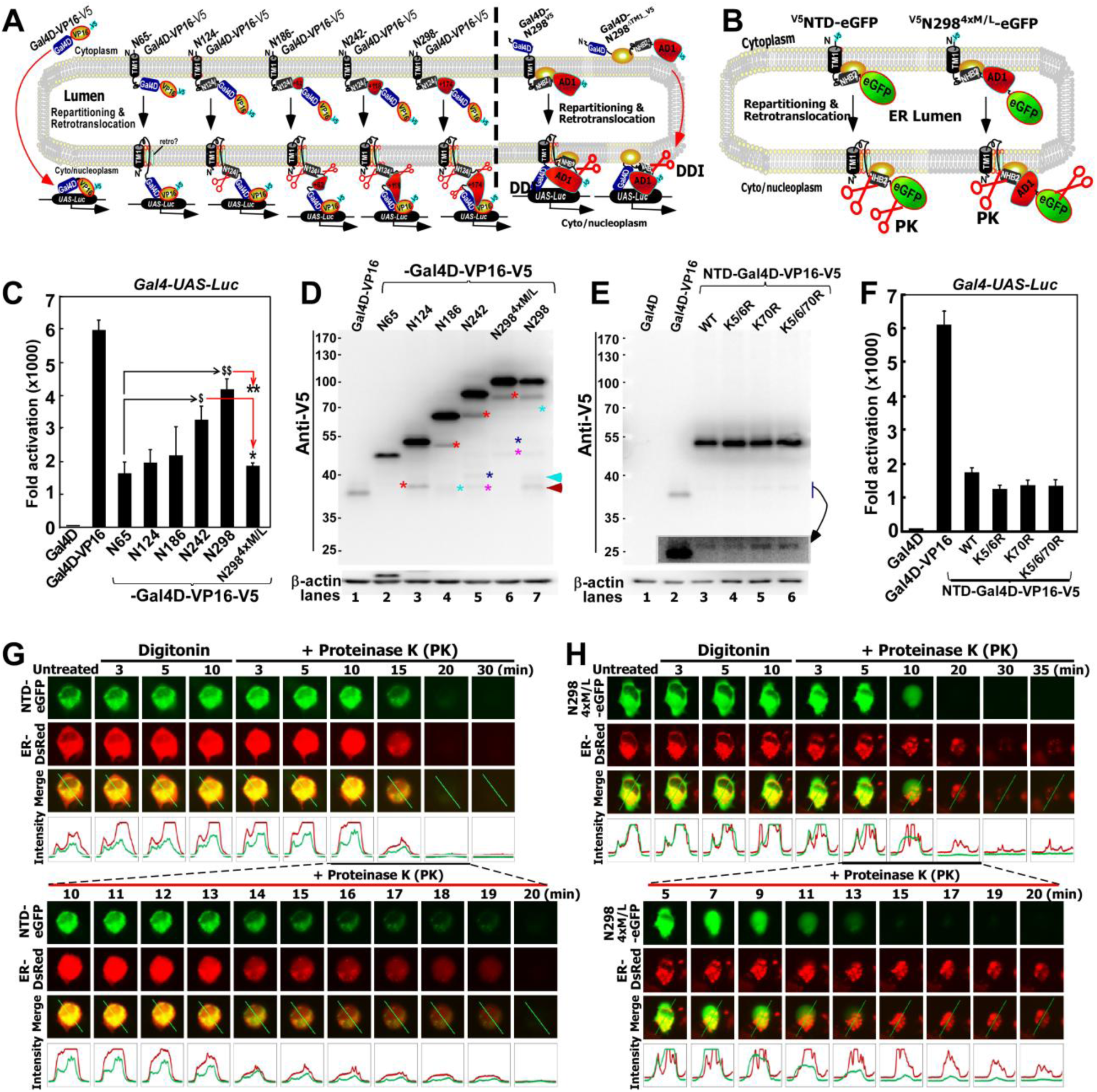
Distinct effects of diverse N-terminal TM1-containing regions of Nrf1 and its mutants on transactivation by its C-terminally-fused Gal4D-VP16 factor occurring after its topovectorial repositioning into the extra-ER compartments. (**A**) A model is proposed to explain the topobiological processes, which control the transactivation activity of N65-, N124-, N186-, N242-N298-fused Gal4D-VP16 factors and their proteolytic processing to yield distinct cleaved proteoforms. The TM1-adjoining portions of these fusion proteins are co-translationally anchored and integrated within ER membranes, whilst their C-terminal Gal4D-VP16 portions are translocated into the lumen. If necessary, the luminal-resident portions will be partially repositioned and retrotranslocated across ER membranes into the cyto/nucleoplasmic side, which leads to the putative proteolytic processing before transactivation of target genes. In contrast, *left panel* shows an additional model to give a explicit interpretation of dynamic membrane-topologies of Gal4D-N298 and its mutant Gal4D-N298^ΔTM1^ (in which Gal4D is attached to the N-terminal end of N298). (**B**) Distinct two models are proposed to interpret dynamic membrane-topologies of the NTD and N298 fusion proteins of Nrf1 with eGFP. (**C**) COS-1 cells had been transfected with each of indicated Nrf1-Gal4D-VP16 factors (1.2 µg), together with *P_TK_UAS/Gal4*-*Luc* (120 ng) and *Renilla* (60 ng) reporter plasmids. Gal4/UAS-driven luciferase gene activity was measured as described for Figure 2A. The data were calculated as a fold change (mean ± S.D, n = 9) over the single Gal4D levels after normalization to the *Renilla* value as an internal control. (**D**) The total lysates of cells expressing N298-Gal4D-VP16 and truncated mutants were resolved by SDS-PAGE containing 10% polyacrylamide and visualized by Western blotting with V5 antibody. (**E, F**) Each of expression constructs for indicated K/R mutants, that had been established on the base of NTD-Gal4D-VP16, alone or in combination with *P_TK_UAS/Gal4*-Luc and *Renilla* reporter plasmids, was co-transfected into COS-1 cells, before total lyates were subjected to Western blotting (***E***) and Gal4/UAS-driven reporter assays (***F***). (**G,H**) COS-1 cells co-expressing either V5-NTD-eGFP or V5-N298^4xM/L^-eGFP, together with the ER-DsRed marker, were subjected to live-cell imaging combined with the *in vivo* membrane protease protection assay. The cells were first permeabilized with 20 μg/ml of digitonin for 10 min, and then co-incubated with 50 μg/ml of proteinase K (PK) about 3 to 35 min. The real-time images were acquired with the Leica DMI-6000 microscopy system. The merged images of V5-NTD-eGFP or V5-N298^4xM/L^-eGFP (*green*) with ER-DsRed (*red*) were placed on *the third row*, whereas changes in the intensity of these signals are shown graphically (*bottom*). The images shown are a representative of at least three independent experiments.

By comparison with N65-Gal4D-VP16, no significant differences in the reporter gene activity were observed from N124-Gal4D-VP16 and N186-Gal4D-VP16 [Figure 9C, N124 (i.e. NTD) and N186 indicate its N-terminal 124-aa or 186-aa regions of Nrf1, respectively]. However, a small fraction of the two fusion proteins was subject to putative proteolytic processing to yield minor cleaved polypeptides of ∼39-kDa and/or ∼51-kDa, besides their full-length proteins of ∼52-kDa and ∼65-kDa, respectively (Figures 9D and S7A, *lanes 3,4*). By contrast, a marked increase in the report gene transactivation by N242-Gal4D-VP16 (containing the N-terminal 242-aa of Nrf1) was measured to ∼3,275-fold (Figure 9C). This was accompanied by a multiple polypeptide ladder composed of between major ∼65-kDa and minor ∼39-kDa, beyond the intact ∼85-kDa N242-Gal4D-VP16 (Figures 9D and S7A, *lane 5*). This suggests that Neh2L facilitates its fusion protein to be partially dislocated from the luminal side of membranes into extra-ER compartments, targeting the protein for its proteolytic processing to generate several digested polypeptides.

A strikingly transactivation activity of Gal4/*UAS-*driven reporter was mediated by an Neh5L-containing fusion protein N298-Gal4D-VP16 to ∼4,190-fold changes (Figure 9C). This was also accompanied by a ladder comprising multiple proteoforms from minor ∼39-kDa to major ∼85-kDa, besides its full-length protein of ∼100-kDa (Figures 9D and S7A, *lane 7*). Conversely, two close proteoforms of ∼39/40-kDa, but not others with higher molecular masses, were almost completely abolished by a mutant of Neh5L (i.e. N298^4xM/L^-Gal4D-VP16, in which all four methionines were changed into leucines) (*lane 6*). However, the reporter gene transactivation activity was not increased, but conversely repressed, by this mutant fusion factor to a so much lower level as being similar to that mediated by N65-Gal4D-VP16 (Figure 9C). Together with the previous data [7, 19], this suggests that partial repositioning of the fusion protein through an Neh5L-triggered mechanism may be restricted by the mutant N298^4xM/L^, due to increases in hydropathicity and aliphaticity of the acidic-hydrophobic amphipathic helix predicted (Figure S7, C *vs* D).

Further mathematic analysis of multiple distinct proteoforms (Figure S7B) unraveled that there exists a dominant proteolytic cleavage site between aa 66-124 (covering a C-terminal half of putative UBL) in Nrf1, even though other regions may also be subject to its proteolytic processing, particularly upon removal of this dominant cleavage site. Moreover, the C-terminally-derived ∼39-kDa polypeptide arising from NTD-Gal4D-VP16 and its transactivation activity appeared to be unaffected by all three K/R mutants (Figure 9, E & F). Another C-terminal ∼85-kDa proteoform arising from N298^4xM/L^-Gal4D-VP16 and its mediated reporter activity were also not significantly altered by those indicated K/R mutants (Figure S7, E & F).

### Dynamic topological movement of V5-NTD-eGFP and V5-N298-eGFP from the ER luminal side of membranes into the cytoplasmic side renders it susceptible to the proteolytic processing by cytosolic proteases

Here, we determined whether N-terminally-tagged Nrf1 fusion proteins have to abide by such rules adhering to dynamic topologies of the original CNC-bZIP protein, which was properly integrated within and around membranes [7, 14, 15, 19]. For this aim, the live-cell imaging of the fusion proteins V5-NTD-eGFP (Figure 9G) and V5-N298^4xM/L^-eGFP (Figure 9H), in combination with *in vivo* membrane protease protection assays, was performed to examine whether Nrf1-fusion proteins are capable of being dislocated from within the lumen to the extra-ER cytooplasmic side of membranes. Experimental cells expressing V5-NTD-eGFP or V5-N298^4xM/L^-eGFP together with the ER/DsRed marker were pre-treated for 10 min with digitonin (to permeabilize cellular membranes) before being challenged with proteinase K (PK) for 3-35 min (still in the presence of digitonin) to digest cytoplasmic proteins. Hence we surmised that if these fusion proteins were transferred from the ER luminal side to the cytoplasmic side of the membrane, it would become vulnerable to digestion by PK. As anticipated, the green fluorescent signals from V5-NTD-eGFP (Figure 9G) and V5-N298^4xM/L^-eGFP (Figure 9H) were superimposed upon the red fluorescent images presented by ER/DsRed. No apparent changes in the intensity of the green signals were observed within 10 min after treatment of the cells first with digitonin and during additional treatment with PK for 10 min (Figure 9G) or 5 min (Figure 9H), when compared to their equivalents from untreated cell states. This observation demonstrates convincingly that the GFP ectope of both fusion proteins was initially translocated into the ER and buried in the lumen, so that it was thus protected by the membrane against PK digestion.

Thereafter, it is interesting to note that the green and red signals from V5-NTD-eGFP and ER/DsRed became gradually fainter as the time of PK treatment was extended from 13 min to 20 min, and both time-lapsed fluorescent signals almost simultaneously disappeared as a result after 20-min PK treatment (Figure 9G, *lower panels*). By contrast, the green signal of V5-N298^4xM/L^-eGFP (Figure 9H) was rapidly weakened after 7-min PK treatment and then gradually became fainter, leading to almost disappearance after 13-min treatment of PK. However, the signal of ER/DsRed was much slowly lapsed as the time of PK treatment was extended to 20 min (Figure 9H*, lower panels*) and the remaining signal appeared to have been retained by 35 min after PK treatment (*upper panels*). Collectively, the difference in these green fluorescence changes observed, irrespective of nuances in these cell states, indicates more rapid dislocation of V5-N298^4xM/L^-eGFP than V5-NTD-eGFP from the ER lumen into extra-ER compartments, whereupon both become susceptible to protease attack. Thereby it is deducible that AD1 is more potent than NTD at controlling the topological repositioning of Nrf1 across membranes into the cyto/nucleoplasm side (Figure 9B). In addition, it should be noted that the V5-NTD-eGFP is retained in the ER as both the fusion protein and ER-DsRed were simultaneously disappeared in 20 min (Figure 9G). This differs from other similar assays (i.e. Figure 9H, refs. 7 & 19) that the signal from ER-DsRed was remaining for at least 30 min after supplementation with PK, suggesting the possible cause for the acceleration of degradation of ER-DsRed.

### A requirement of p97-driven retrotranslocation for selective proteolytic processing of V5-Nrf1-eGFP to remove the client N-terminal portions during maturation of the CNC-bZIP transcription factor

To gain an in-depth understanding of the functional consequences, cells expressing V5-Nrf1-eGFP were pretreated with NMS-873 and subjected to the pulse-chase experiments to determine a role of p97-driven retrotranslocation in the proteolytic processing of the CNC-bZIP protein to remove its N-terminal portions during maturation into an activator. As expected, treatment with NMS-873 caused a time-dependent accumulation of V5-Nrf1-eGFP and its N-terminal polypeptides of between ∼160-kDa and ∼12.5-kDa (Figure 10A, *lanes 1-4*). Following recovery of p97 from its inhibitor, the existing V5-Nrf1-eGFP and its N-terminal derivates was released into the extra-luminal side of membranes, allowing them to be proteolytically processed and degraded by cytosolic proteases (i.e. proteasomes) (*lanes 5-8*). This resulted in the rapid turnover of V5-Nrf1-eGFP, its two major ∼36-kDa and 12.5-kDa polypeptides, which behaved with largely similar half-lives of 0.68 h (=41 min), 0.80 h (=48 min) and 0.67 h (=40 min) after CHX treatment, respectively (Figure 10B, *green curves*). By contrast, their turnover was obviously mitigated by MG132 (Figure 10A, *lanes 9-12*), such that their half-lives were extended to over 4 h (Figure 10B, *blue curves*). As such, a fraction of these proteins was gradually degraded, even in the presence of this proteasomal inhibitor, implying that they may be destructed to disappear by other proteases beyond proteasomes.

**Figure 10.**
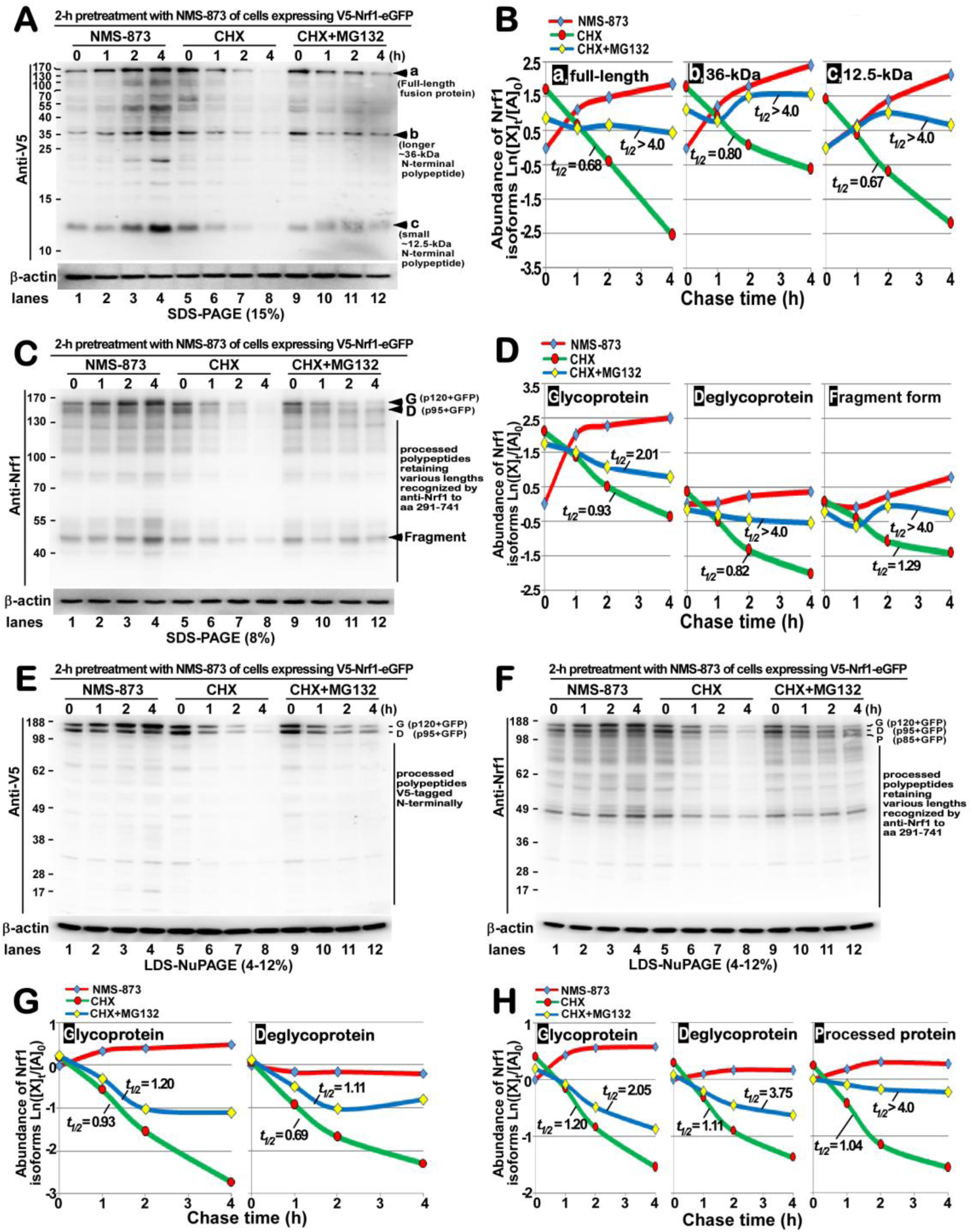
An effect of p97 inhibitors on the topovectorial repositioning and proteolytic processing of V5-Nrf1-eGFP fusion protein to yield distinct isoforms. COS-1 cells expressing V5-Nrf1-eGFP were pretreated with NMS-873 (10 µmol/L) for 2 h, and then transferred in another fresh medium containing either NMS-873 (10 µmol/L) or CHX (50 µg/ml) alone or plus MG132 (5 µmol/L) to continue the culture for distinct lengths of time before being harvested in denatured lysis buffer. The total lysates were subjected to protein separation by SDS-PAGE gels containing 15% polyacrylamide (*A*) or 8% polyacrylamide (*C*), or LDS-NuPAGE gels 4-12% polyacrylamide (*E, G*), followed by visualization by Western blotting with antibodies against Nrf1 or its N-terminal V5 tag. The electrophoretic locations of the intact V5-Nrf1-eGFP and derived isoforms were indicated. The single letters *G*, *D* and *P* represent glycoprotein, deglycoprotein and processed mature protein, respectively. In addition, the relative intensity of indicated protein bands (*A, C, E, F*) was quantified and shown graphically (*B, D*, *G*, *H*), with distinct *t_1/2_* values estimated, respectively.

The parallel experiments were carried out by employing either SDS-PAGE gels containing 8% polyacralymide (Figure 10C) or NuPAGE gels containing 4-12% polyacralymide (Figure 10E). The intact V5-Nrf1-eGFP was represented by two major bands of glycoprotein (i,e. p120 Nrf1 + 28.5-kDa GFP) and deglycoprotein (i.e. p95 Nrf1 + 28.5-kDa GFP). They were visualized by antibodies against Nrf1 (Figure 10C) or its N-terminal V5 tag (Figure 10E) and also identified by assaying *in vitro* deglycosylation reactions (Figure S8). Further examinations revealed that arrest of p97 by NMS-873 led to an incremental enhancement in the glycoprotein of V5-Nrf1-eGFP, but not its deglycoprotein (Figure 10 C to E, *lane 1-4*). This demonstrates that the intact glycoprotein of V5-Nrf1-eGFP is accumulated in the lumen, whereas its deglycoprotein is released in extra-luminal compartments and therefore unaffected by the p97 inhibitor (Figure 10D). Subsequently, a recovery of p97 for the retrotranslocation conferred the existing glycoprotein of V5-Nrf1-eGFP and its deglycoprotein to be rapidly turned over (Figure 10E), with their half-lives estimated to be 0.93 and 0.69 h after CHX treatment, respectively (Figure 10G). The turnover of both proteins was modestly postponed by MG132, so that their half-lives were slightly prolonged to similar 1.20 and 1.11 h (Figure 10E & G).

Further pulse-chase experiments with anti-Nrf1 immunoblotting of cell lysates (Figures 10F and S8B) convincingly demonstrated that inhibition of p97-driven retrotranslocation by NMS-873 caused the intact V5-Nrf1-eGFP to be sequestered in the ER and thus resulted in a concomitant accumulation of its glycoprotein in the luminal side of membranes. Upon recovery of p97 from being arrested by NMS-873, the client glycoprotein of V5-Nrf1-eGFP was dynamically repositioned into the extra-ER cyto/nucleoplasmic side. Therein, this glycoprotein was allowed for deglycosylation and proteolytic processing to generate various lengths of proteoforms. Together with the previous data [7, 12, 19], these indicate that deglycoprotein of V5-Nrf1-eGFP and its N-terminally-derived polypeptides are unstable, due to being rapidly degraded by proteasomes and other proteases (Figure 10H). However, it is fortunate that the maturation process of Nrf1 enables it to become an active processed CNC-bZIP transcription factors.

### The proteolytic processing of Nrf1 to generate a mature ∼85-kDa CNC-bZIP transcription factor is determined by p97-driven retrotranslocation pathway

To provide a better explanation of the functional consequence from the multistage post-translational processing of Nrf1 to generate a mature CNC-bZIP transcription factor, Nrf1-expressing cells were pretreated with NMS-873 and then subjected to the time-course analysis to further elucidate the protein processing steps. As excepted, western blotting of cell lysates with Nrf1-specific antibodies (Figure 11A) and its C-terminally-tagged V5 antibodies (Figure 11B) showed that abundance of the 120-kDa full-glycoprotein (and ∼105-kDa partial-deglycoprotein) was incremented along with increasing time of NMS-873-arrested p97 (Figure 11C). By contrast, this p97 inhibitor did not lead to any obvious changes in both the 95-kDa deglycoprotein of Nrf1 and its N-terminally-truncated 85-kDa isoform (Figure 11, A to C).

**Figure 11.**
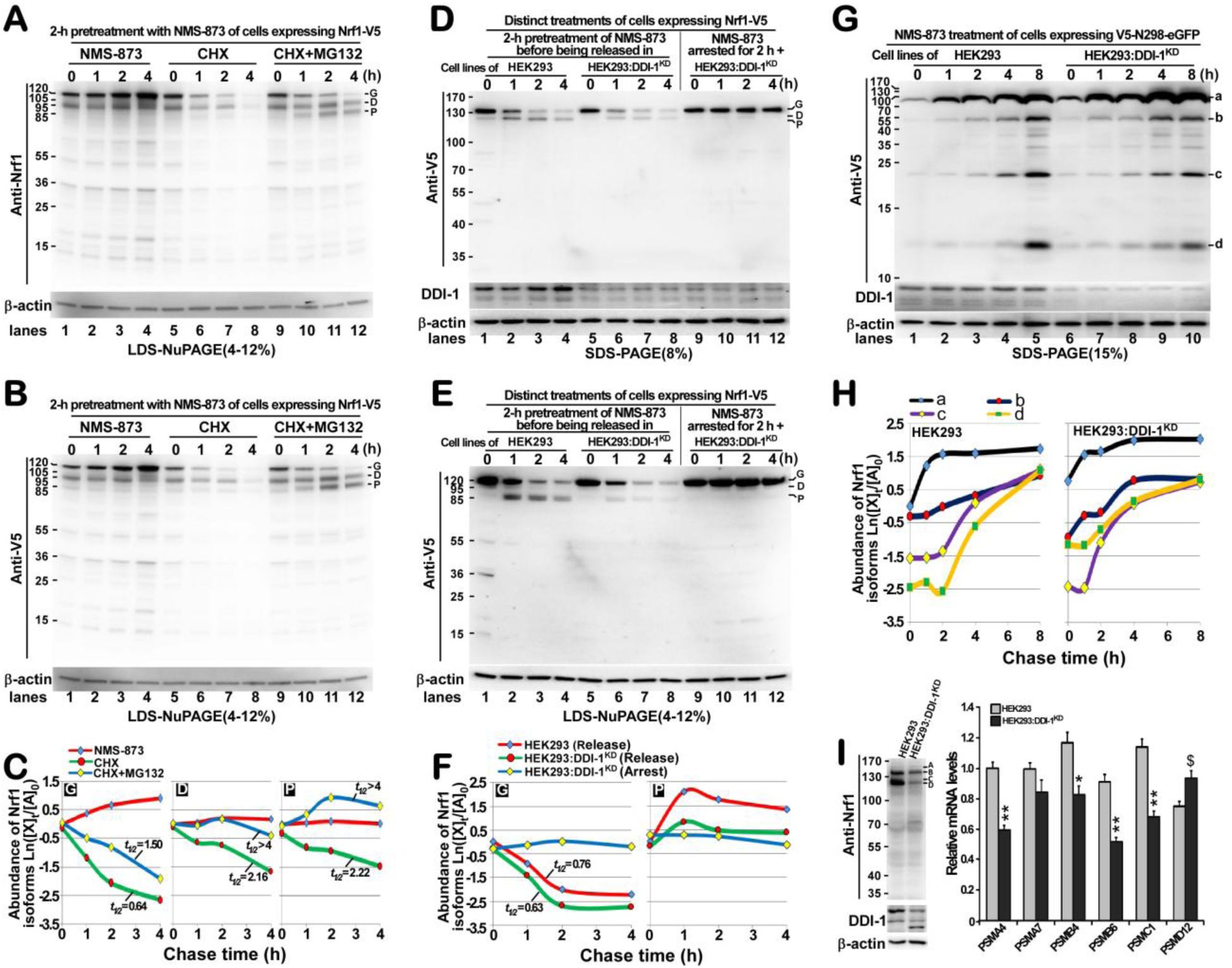
DDI-1-mediated proteolytic processing of Nrf1 into a mature CNC-bZIP transcription factor is determined by p97-driven retrotranslocation and -independent pathways. (**A** to **C**) Nrf1-expressing COS-1 cells were pretreated with NMS-873 (10 µmol/L) for 2 h, and then transferred in another fresh medium containing either NMS-873 (10 µmol/L) or CHX (50 µg/ml) alone or plus MG132 (5 µmol/L) for continue the culture for additional lengths of times as indicated. The total lysates were subjected to protein separation by 4-12% LDS-NuPAGE gels and visualization by immunoblotting with antibodies against Nrf1 (***A***) or its C-terminal V5 tag (***B***). The resultant electrophoretic locations of Nrf1 glyprotein, deglycoprotein and processed mature protein were indicated by single letters ***G, D*** and ***P*** respectively. (**D** to **F**) Wild-type HEK293 and/or DDI-1-difficient HEK293:DDI-1^KD^ cell lines were transfected with an expression construct for Nrf1-V5 and then treated with each of indicated chemicals alone or in distinct combinations. Subsequently, the lyastes were resolved in either SDS-PAGE gels containing 8% polyacrylamide (***D***) or LDS-NuPAGE gels containing 4-12% polyacrylamide (***E)***, followed by immunoblotting with V5 antibody. (**G,H**) Similar experiments was performed in both HEK293 and HEK293:DDI-1^KD^ cell lines that had been transfected with V5-N298-eGFP before being analyzed as described above. Of note, the relative intensity of protein bands as indicated in the panels (***B,E,G***) was quantified as shown graphically in the panels (*C,F,H*). (**I**) *Left panel* shows Western blotting of HEK293 and HEK293:DDI-1^KD^ with antibodies against Nrf1 or DDI-1. *Right panel* shows real-time qPCR analysis of Nrf1-target genes (encoding distinct 26S proteasomeal subunits) such as *PMSA4*, *PSMA7*, *PSMB4*, *PSMB6*, *PSMC1* and *PSMD12*.

Upon the recovery of p97-driven retrotranslocation from its inhibitor, the luminally-sequestered glycoprotein of Nrf1 was enabled for dynamic repositioning to dislocate extra-ER cyto/nucloplasmic side of membranes. Therein, the 120-kDa Nrf1 glycoprotein was deglycosylated to become a deglycoprotein of ∼95-kDa. The deglycoprotein is, transient, though restricted to membranes, because it is further subject to the proteolytic processing by cytosolic proteases so as to remove its N-terminal membrane-anchored portions. Thus, the proteolytic processing of Nrf1 led to generation of an mature ∼85-kDa CNC-bZIP transcription factor, as a functional consequence. This notion is evidenced by the fact that treatment of NMS-873-pretreated cells with CHX resulted in dramatical effects on the abundance and stability of the existing 120-kDa Nrf1 glycoprotein, its derivates 95-kDa deglyprotein and 85-kDa processed protein (Figure 11A, B, *lanes 5-8*). Their distinct half-lives were determined to be 0.64 h (=38 min), 2.16 h (= 130 min) and 2.22 h (=133 min) after CHX treatment, respectively (Figure 11C). Notably, the rapid turnover of Nrf1 120-kDa glycoprotein was modestly ameliorated by MG132 (*lanes 9-12*), which was defined with its half-life slightly prolonged to 1.50 h (= 90 min). By contrast, the stability and abundances of its 95-kDa deglycoprotein and particularly 85-kDa processed proteins were significantly augmented by this proteasomal inhibitor, along with both half-lives extended to over 4 h of the experimental time (Figure 11C). Interestingly, there existed a time-dependent decrease in the existing 120-kDa Nrf1 glycoprotein, even in the presence of proteasomal inhibitor (Figure 11A,B, *lanes 9-12*). This was coherently accompanied by distinct increases in yield of both the 95-kDa Nrf1 deglycoprotein and its processed 85-kDa isoform. The increase is not only attributable to the proteasomal-limited degradation and is also relevant to the feedback response required for the selective proteolytic processing of Nrf1 to yield a mature active transcription factor, as consistent with other previous reports [16, 18, 20].

### DDI-1-mediated proteolytic processing of Nrf1 into a mature active CNC-bZIP factor is monitored by p97 during the client retrotranslocation into the extra-ER cyto/nucleoplasmic side

To elucidate the significant role of p97 for the retrotranslocation of Nrf1 in its proteolytic processing by DDI-1, a specific cell line HEK293:DDI-1^KD^ has been established by CRISPR/Cas9-mediated genome editing of *DDI-1* to be killed deadly. Following release of p97 from the arrest by NMS-873 in HEK293:DDI-1^KD^ cells, a time-lapsed decrease in abundance of Nrf1 glycoprotein was observed (Figure 11D,E, *lanes 5-8*), but this was almost not accompanied by a putative increment in its deglycoprotein and processed protein. By contrast, albeit wild-type HEK293 cells also showed a gradual decrease in the 120-kDa Nrf1 glycoprotein, the latter form was concomitantly replaced by an increase in its processed 85-kDa isoform (*lanes 1-4*). In addition, a roughly similar stability of Nrf1 glycoprotein was determined with its half-lives of 0.63 and 0.76 h, respectively, in HEK293:DDI-1^KD^ and wild-type HEK293 cell lines (Figure 11F). The rapid turnover of Nrf1 glycoprotein was, however, almost completely prevented by NMS-873 in HEK293:DDI-1^KD^ cells (lanes *9-12*). Collectively, these data demonstrate that the processed ∼85-kDa protein of Nrf1 is generated from the DDI-1-mediated proteolytic processing of a transient ∼95-kDa Nrf1 deglycoprotein; the latter arises from deglycosylation of intact ∼120-kDa glycoprotein. This successive event occurs after retrotranslocation of Nrf1 glycoprotein by p97 into the extra-ER cyto/nucleoplasmic side of membranes.

Surprisingly, the rapid turnover of those extra-ER cytoplasmically-localized Nrf1 isoforms is neither prevented nor ameliorated by deficiency of DDI-1 in HEK293:DDI-1^KD^ cells. This finding indicates additional requirements of proteasomes and other proteases for the Nrf1 processing into an active CNC-bZIP transcription factor. Therefore, It is reasonable that, during the mature processing of Nrf1, various lengths of its N-terminal UBL-adjoining portions should be progressively proteolytically removed by DDI-1 and other proteases besides proteasomes. Such selective proteolytic processing of Nrf1 occurs, however, only after some of potential degrons and cleavage sites within its UBL-adjoining regions would be repositioned and exposed to the enzymatic active centers of cytosolic proteases on the cyto/nucleoplasmic side. Intriguingly, the blockage of p97-driven retrotranslocation by NMS-873 resulted in a time-dependent increment of intact 80-kDa V5-N298-eGFP, along with its N-terminally-derived polypeptides of between 55-kDa and 12.5-kDa (Figure 11G,H). By contrast with wild-type cells, HEK293:DDI-1^KD^ cells showed a further accumulation of intact V5-N298-eGFP, but this was accompanied by significant decreases in yield of its N-terminal polypeptides (*lanes 6-10*). Together, these indicate that potential degrons and cleavage sites within the N-terminal membrane-protective portion of Nrf1 may partially repositioned by p97-driven retrotranlocation and -independent pathways, gaining a local access to cytosolic proteases involved in the juxtamembrane proteolysis.

Western blotting of HEK293:DDI-1^KD^ lysates revealed deficiency of DDI-1 significantly diminished expression of endogenous Nrf1 protein-C/D to much low levels, but its longer intact protein-A/B was almost unaffected (Figure 11I, *left panel*). Further quantitative PCR analysis unraveled marked decreases in the mRNA expression of Nrf1-target genes encoding proteasomal subunits, such as *PSMA4, PSMA7, PSMB4, PSMB6* and *PSMC1*, but not *PSMD12* in HEK293:DDI-1^KD^ cells, when compared with those obtained from HEK293 cells (Figure 11I, *right panel*). This indicates that DDI-1 is also required for the endogenous Nrf1 processing into a mature active CNC-bZIP transcription factor before regulating cognate target genes.

## DISCUSSION

To provide a better understanding of topovectorial processes that regulate juxtamembrane proteolysis (RJP) of Nrf1 (Figure 12), experimental evidence has been herein presented revealing that: (i) portions of Nrf1-, its NTD- and N298-fused proteins are dynamically repartitioned from the ER luminal side of membranes into the extra-ER side, whereupon they are allowed for the selective proteolytic processing; (ii) the proteolytic processing of Nrf1 is predominantly determined by its TM1-associated NHB1 peptide, and finely tuned by its CRAC sequence situated between NHB1 and NHB2 (which together could be refolded into a membrane-regulated UBL module, Figure S1); (iii) the N-terminal UBL-adjoining regions of Nrf1 are progressively proteolytically truncated by cytosolic proteases (e.g. DDI-1/2 and proteasomes) in close proximity to the extra-luminal interface of ER membranes; (iv) an N-terminal 12.5-kDa polypeptide of Nrf1 arises from the proteolytic processing of its NHB2-adjoining region (aa 66-124), albeit other regions may be subjected to progressively proteolytic processing; (v) the mature processing of Nrf1, as a functional consequence, contributes to transcriptional regulation of cognate target genes; and (vi) both p97-driven retrotranslocation and -independent pathways are required for the topovectorial processing of Nrf1 into a mature active ∼85-kDa CNC-bZIP transcription factor.

**Figure 12.**
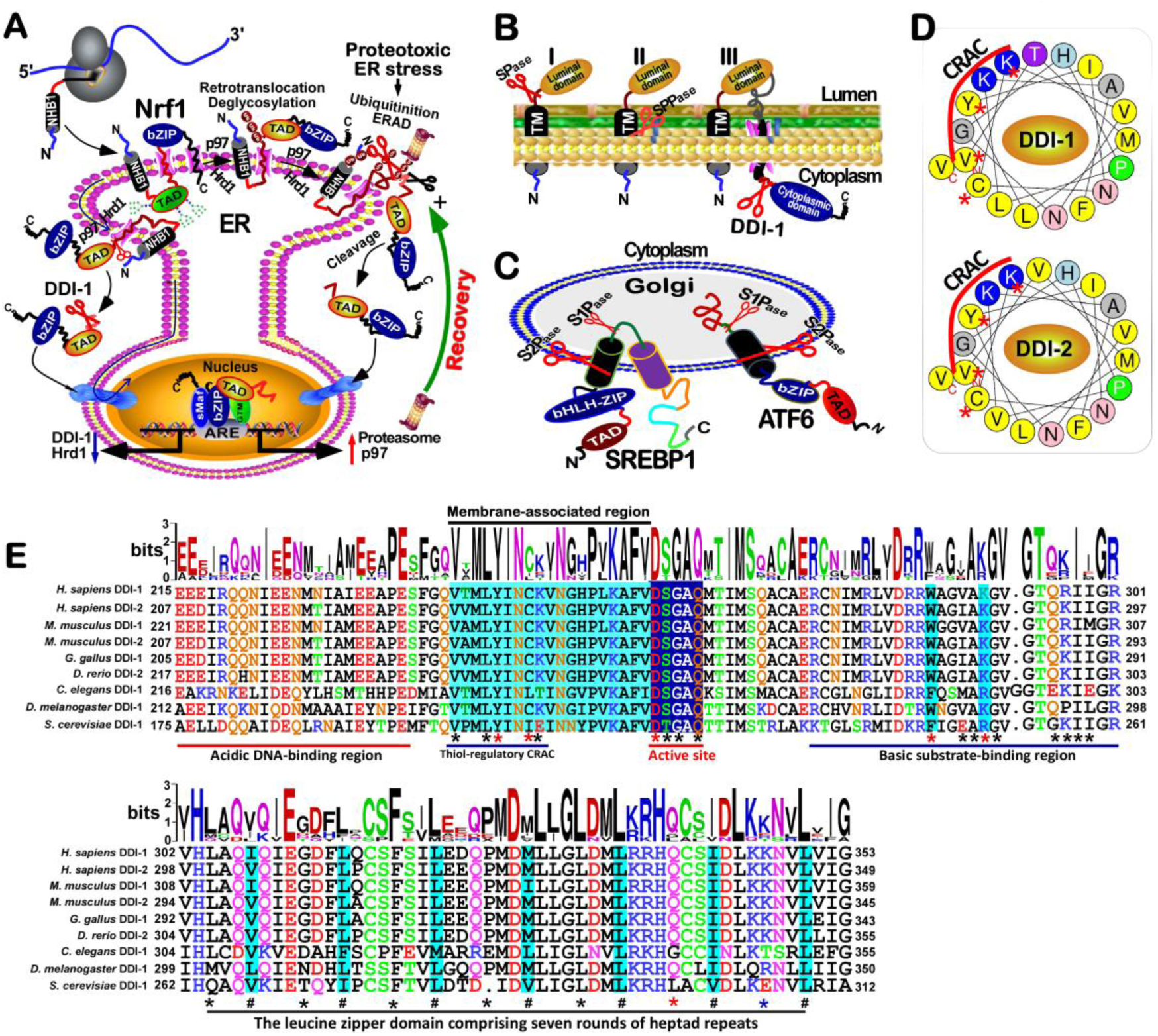
The regulated juxtamembrane proteolysis (RJP) of Nrf1 is distinctive from other mechanisms accounting for the processing of signal peptides and the regulated intramembrane proteolysis (RIP) of ATF6 and SREBP1. (**A**) A model is proposed to provide a better understanding of topovectorial mechanisms that control dynamic moving of Nrf1 in and out of the ER and its juxtamembrane proteolytic processing into a mature CNC-bZIP transcription factor that regulates target genes. (**B**) Three models are illustrated to give an explicit explanation of their distinctions. *Model I* shows the proteolytic processing of an ER-targeting signal peptide by SPase (sign al peptidase); *Model II* indicates the regulated intramembrane proteolysis by SPPase (signal peptide peptidase); and *Model III* displays the regulated juxtamembrane proteolysis of Nrf1 by DDI-1 (or other unidentified proteases), enabling relevant membrane-proteins to be released from the membrane confinements during their maturation. (**C**) As consistent with the above *Models I* and *II*, the regulated intramembrane proteolysis (RIP) accounts for the proteolytic processing of SREBP1 and ATF6 by Site-1 protease (SIPase, acting as an equivalent to SPase) and Site-2 protease (S2Pase, acting as an equivalent to SPPase). (**D**) A CRAC-adjoining region immediately close to the active site of DDI-1/2 may refold into a basic-hydrophobic amphipathic helix, enabling the enzymatic center to lie on the cytoplasmic plane of membrane bilayer lipids and thus facilitate the juxtamembrane proteolysis of Nrf1. (**E**) These nine amino acid sequences of DDI-1 and DDI-2 from distinct species were aligned. This aligned region covers the enzymatic active sites of DDI-1/2 and their flanking regions for its DNA-binding, membrane-associated, and substrate-binding, along with the adjacent leucine zipper domain. Furthermore, the enzymatic active center within DDI-1/2 is flanked immediately by its N-terminal membrane-bound CRAC-adjoining motif and its C-terminal substrate-binding regions. The CRAC-adjoining region may refold into a basic-hydrophobic amphipathic helix, whilst the basic-enriched substrate-binding region and adjacent leucine zipper domain of DDI-1/2 should be allowed for a putative interaction with Nrf1.

### Distinct topovectorial processes of Nrf1 dictate dislocation of its fusion proteins from the ER lumen into extra-ER cyto/nucleoplasms where they are proteolytically processed before transactivating target genes

The proper membrane-topological folding of Nrf1 within and around the ER is determined by its NHB1-associated TM1 peptide in cooperation with other amphipathic regions (e.g. TMi, TMp and TMc) [7, 13, 14, 19]. This notion is further supported by live-cell imaging of V5-NTD-eGFP and V5-N298^4xM/L^-eGFP, combined with *in vivo* membrane protection assay against proteolytic digestion by proteinase K. To give an explicit explanation, a topobiological model was proposed and illustrated (Figures 3H, 9B and 12A). The TM1-adjoining NHB1 peptide of Nrf1 is integrated in an N_cyto_/C_lum_ orientation within ER membranes, whilst the CRAC1/2-adjacent NHB2 and AD1 regions, along with the C-terminally-fused GFP, are partitioned on the ER luminal side. By contrast, the N-terminally-tagged V5 ectope was hence positioned to face on the cytoplasmic side. Subsequently, dynamic repositioning of the luminal-resident portions into the cytoplasmic side makes it susceptible to protease attack. The retrotranslocation of V5-NTD-GFP or V5-N298-GFP to move out of the ER is monitored by several amphipathic regions, such as CRAC1/2-adjacent NHB2, DIDLID/DLG and Neh5L within AD1 ([7, 19] and this study). This is further evidenced by other experiments, revealing that aa 81-90 (covering an N-terminal half of NHB2), aa 171-186 (covering DIDLID/DLG), and aa 125-170 (i.e. the N-terminal PEST1-adjoining one-third of AD1) are required for the correct topological folding of intact V5-N298-GFP and its protein stability. This conclusion is drawn from the fact that deletion of each of these three regions from the wild-type leads to rapid disruption of their corresponding full-length mutant proteins, so as to disappear and also be replaced by multiple degraded proteoforms.

In order to verify the conceptual statement that topovectorial regulation of NTD- or N298-fused Gal4 proteins determines dislocation of their chimeric factors into extra-ER cyto/nucleoplasmic compartments, gaining an access to cognate target genes and transactivating them, two distinct strategic sets of expression constructs for different lengths of Gal4-based fusion proteins have been created. Firstly, attachment of Gal4D to the N-terminus of N298 (Gal4D-N298^V5^) and various lengths of its N-terminally TM1-truncated mutants (e.g. Gal4D-N298^ΔTM1_V5^) results in at least two completely different topological folding models as proposed (Figure 9A). Based on the current knowledge of membrane-topology [26–28], it is predicted that the basic Gal4D portion of Gal4D-N298^V5^ is retained on the cytoplasmic side of membranes, whilst AD1 of N298 should be transiently translocated into the ER lumen. Only after the luminally-localized AD1 element is retrotranslocated across membranes into the cyto/nucleoplasmic side, it could serve as a *bona fide* functional domain to mediate transactivation of Gal4/UAS-driven reporter gene. By contrast, all those N-terminally TM1-truncated mutants of with N298 cause the resultant fusion factors (e.g. Gal4D-N298^ΔTM1_V5^) to be not integrated within the ER, such that their AD1-mediated activity to transactivate Gal4 reporter gene expression is no longer restrained by this organelle. This is further supported by additional evidence showing that negative regulation of Gal4D-N298^V5^ by NTD is conversely disinhibited by removal of TM1-adjoining NHB1 alone or in combination with CRAC1/2-adjacent NHB2 peptides, which causes obvious two-stage stepwise increases in AD1-mediated transactivation of the Gal4 reporter gene (Figure 2A). Interestingly, the increase at the second stage is postulated to be attributable to being released from a TM-independent inhibitory confinement, albeit CRAC1/2-adjacent NHB2 peptide is enabled for binding to cholesterol-enriched membranes.

Secondly, the notion that topovectorial regulation of NTD- or N298-fused Gal4D-VP16 proteins dictates their dislocation into extra-ER subcellular compartments has also been validated through another strategic way. As illustrated (Figure 9A), several sandwiched proteins have been created by C-terminal attachment of the Gal4D-VP16 chimeric ectope to N298 and various lengths of C-terminally-truncated mutants. Theoretically, it is inferred that all these TM1-containing Gal4D-VP16 sandwiched proteins should be integrated within the ER, whilst Gal4D-VP16 chimeras is co-translocationally positioned on the luminal side of membranes and thus buried in the lumen, so that it is unable to transactivate Gal4-target reporter gene. Conversely, the reporter gene transactivation would occur upon dislocation of Gal4D-VP16 chimeras from the luminal side of membranes into the cyto/nucleoplasmic side. This deduction has been experimented by Gal4/*UAS-Luc* reporter assays, showing that its transcriptional activity is mediated by each of those sandwiched proteins including N65-Gal4D-VP16, NTD-Gal4D-VP16, N186-Gal4D-VP16, N242-Gal4D-VP16, N298-Gal4D-VP16 (Figure 9C), albeit their transactivation abilities are much lower than the value measured from a membrane-free Gal4D-VP16 chimeras. This demonstrates that these artificial membrane-bound transcription factors are predominantly restricted with the ER through their TM1-adjoining NHB1, in cooperation with CRAC1/2-adjacent regions. As such, they are partially dislocated into extra-ER compartments and subjected to the proteolytic processing to yield multiple cleaved proteoforms, before transactivating target gene expression. Further comparison of N298-Gal4D-VP16 and N298^4xM/L^-Gal4D-VP16 deciphers that the former transactivation activity is significantly suppressed by the mutant. This is due to that this mutant can block the in-frame translation into a free protein of Gal4D-VP16. In addition, intact Neh5L-driven retrotranslocation of its fusion factor could also be restricted by its mutant, with enhanced hydropathicity and aliphaticity of the helix folded (Figure S7D).

### Different topovectorial organizations of Nrf1 dictate selective proteolytic processing of its fusion proteins by cytosolic proteases to yield multiple proteoforms including its N-terminal 12.5-kDa polypeptide

Although Nrf1 is co-translationally translocated in the ER lumen, none of the luminal-resident proteases (e.g. signal peptidase and Site-1 protease, Figure 12B,C) have been hitherto identified. Thereby, a conclusion is reached that Nrf1 is not proteolytically processed, but rather N-glycosylated, on the ER luminal side of membranes [12–15]. However, upon dislocation of Nrf1 into the extra-ER cyto/nucleoplasmic side [29], it can be allowed for deglycosylation and proteolytic processing in order to yield multiple cleaved proteoforms [7, 19]. This functional processing is executed by other cytosolic proteases (e.g. DDI-1/2) [24, 25] rather than 26S proteasomes [18, 21, 22]. The conclusion is also supported by our evidence provided herein, albeit detailed mechanisms by which distinctive doses of proteasomal inhibitors cause dual opposing effects on Nrf1-target gene expression are not well understood.

Interestingly, two roughly parallel trend lines are illustrated (Figure S7B), which are defined by molecular masses of various N-terminal TM1-containing portions of Nrf1 that fused with the Gal4D-VP16 chimeras and their major processed polypeptides. From this, it is deduced that a dominant proteolytic cleavage site exists at NHB2-adjoining peptide of between aa 66-124 (covering the C-terminal half of UBL), albeit other adjacent regions may also be proteolytically processed particularly upon removal of this dominant cleavage site. This finding appears to be further supported by other evidence obtained from putative proteolytic processing of Gal4D/Nrf1^1-298^ and those progressively NTD-lengthened deletion mutants till Gal4D/Nrf1^107-298^ (Figures 2B and S2). All these fusion proteins give rise to a major processed polypeptide of ∼39-kDa estimated close to the molecular mass of the full-length Gal4D/Nrf1^188-298^. This processing event was, however, prevented by additional two mutants Gal4D/Nrf1^120-298^ and Gal4D/Nrf1^170-298^, even in the presence of proteasomal inhibitors. Notably, among all those proteoforms visualized with antibodies against their C-terminal V5 ectope, the processed ∼39-kDa polypeptide is postulated to result from a putative protease cleavage occurring within endopeptide bonds of between aa 108-187.

In order to precisely determine the selective proteolytic processing of Nrf1 through its NTD alone or plus AD1 within N298, a series of doubly-tagged fusion proteins of either NTD or N298 have been made by using N-terminal V5 (VSVg or StrepII) tags and C-terminal GFP (Figures 3 and S3). It is of critical importance to notice that all these doubly-tagged proteins should be properly folded in accordance with the correct topological orientation of NTD or N298 within and around ER membranes [12–14]. This is reasoned by the fact that they have been engineered to abide by the positive-inside rule and are also defined by net charge differences between both N-terminally and C-terminally flanking regions of the hydrophobic TM1 core spanning membranes [26–28]. The resulting evidence has been provided here revealing that these doubly-tagged chimeric proteins are proteolytically processed to yield multiple degraded polypeptides of between 70-kDa and 12.5-kDa, one of which is represented by the major N-terminal 12.5-kDa polypeptide of Nrf1.

### Yield of the N-terminal 12.5-kDa polypeptide from the proteolytic cleavage of Nrf1 within its UBL-adjoining peptides is determined by its membrane-topovectorial folding and repositioning into cyto/nucleoplasms

As described above, the N-terminal 12.5-kDa and other longer polypeptides of Nrf1 have been shown to derive primarily from the selective proteolytic processing of NTD or N298 at UBL-adjoining NHB2 peptides within distinct fusion contexts. However, these N-terminal polypeptides are still yielded, but not abolished, even upon deletion of some NHB2-adjoining peptides between aa 81-90 (i.e. an N-terminal half of NHB2), aa 125-170 (i.e. the N-terminal one-third of AD1) and aa 171-186 (i.e. the DIDLID/DLG element). All three mutant proteins should be, *bona fide*, rapidly degraded to disappear and replaced by multiple degraded forms. This implies that aa 81-186 of Nrf1 are required for the correct membrane-topological folding of intact chimeric proteins and their stability.

Further mapping of internal deletion mutants within NTD and AD1 revealed that yield of the N-terminally-derived 12.5-kDa polypeptide is required for the presence of aa 11-22, acting as the core hydrophobic h-region of the TM1-containing NHB1 peptide anchored within membranes. This demonstrates that the proteolytic processing event occurs in close proximity to the ER and depends on the TM1 anchor within membranes. More interestingly, removal of either CRAC1/2-adjoining peptide (aa 31-80) or the entire NHB2 peptide (aa 81-106) conferred on the resulting mutants to yield a minor fast-migrating polypeptide close to 12.5-kDa. This indicates that the N-terminal 12.5-kDa polypeptide of Nrf1 is yielded by its proteolytic cleavage on the C-terminal border of the NHB2-adjoining peptides. Similar N-terminal polypeptides of ∼12.5-kDa was weakened by loss of aa 107-186 or 280-298 (covering Neh5L), but no changes in its electrophoretic mobility were observed. Therefore, it is deducible that yield of the N-terminal 12.5-kDa polypeptide of Nrf1 is also monitored by repositioning of its selective proteolytic cleavage site between aa 107-186 (Figure 1A). This processing is defined by dynamic repartitioning of NHB2-flanking DIDLID/DLG and Neh5L regions from the luminal side of membranes into extra-ER cyto/nucleoplasms. This topovectorial event facilitates the N-terminal proteolytic processing of Nrf1 to give rise to multiple proteoforms.

Intriguingly, yield of the N-terminal 12.5-kDa polypeptide of Nrf1 was unaffected, though being endowed with a slightly faster mobility, by loss of the C-terminal two-fifths of NHB2 (in V5-N298^Δ96-106^-eGFP), but significantly prevented rather than completely abolished by additional two tripeptide mutants of within the above same region (i.e. V5-N298^Δ103-105^-eGFP V5-N298^W103A/L104A^-eGFP). By contrast, another two mutants V5-N298^Δ100-102^-eGFP and V5-N298^N101A/A102L^-eGFP only caused an enhancement in the slightly fast-migrating polypeptide close to 12.5-kDa. These seemingly paradoxical findings, together with the previous reports [7, 19, 25, 29], have led us to surmise that selective proteolytic processing of Nrf1 subjected to efficient cleavage is much likely to occur within the C-terminal ^103^WLV^105^ tripeptide of NHB2 and adjacent peptide sequences (Figures 1A and 12A). Notably, this efficient processing should depend on distinct local topologies of NHB2-adjoining regions in different contexts surrounding membranes, which dictate the *bona fide* repositioning of them from the ER luminal side of membranes into cyto/nucleoplasmic compartments.

Moreover, it is quite baffling that abundances of intact V5-N298-eGFP and its N-terminal 12.5-kDa polypeptide are obviously increased by inhibition of p97-driven retrotranslocation. It is inferable that some N-terminal portions, i.e. CRAC1/2-adjacent NHB2 peptides, of Nrf1 are retained on the ER luminal side within a certain time extended by arrest of p97, and thus transiently protected by membranes against cytosolic proteolytic degradation. However, the proteolytic processing of V5-N298-eGFP to remove its N-terminal 12.5-kDa polypeptide is not suppressed, but conversely promoted, by blockage of p97-feuled dislocation of the client protein into extra-ER compartments. This suggests that there exists a not-yet-identified p97-independent mechanism (e.g. the putative membrane-flipping translocation of cholesterol-enriched microdomains associated with the CRAC peptide of Nrf1).

### The putative ubiquitination of UBL-adjoining NTD and AD1 is not a necessary prerequisite for the selective proteolytic processing of Nrf1 (and its fusion proteins) to yield multiple proteoforms

Distinct topovectorial events enable Nrf1 (and longer TCF11) to be selectively processed in different tempo-spatial subcellular locations. It is specifically modified in a variety of post-translational fashions, including *N*-glycosylation, *O*-GlcNAcylation, deglycosylation, phosphorylation, ubiquitination, degradation and proteolytic cleavage. These results in the yield of multiple distinct proteoforms (i.e. 120-kDa, 95-kDa, 85-kDa, 55-kDa, 46-kDa, 36-kDa and 25-kDa estimated by NuPAGE), which exert different and even opposing abilities to mediate expression of target genes [1, 38]. Besides glycosylation of Nrf1 and TCF11 on the ER lumen and ensuing deglycosylation on the extra-ER cyto/nucleoplasms [7, 13, 15, 19, 39], both are also ubiquitinated by Hrd1-, β-TrCP- and Fbw7-mediated pathways in cyto/nucleoplasmic compartments. In turn, this ubiquitin-proteasomal degradation is controlled by Nrf1/TCF11 *via* an ER-associated degradation (ERAD) feedback loop [16–18, 20, 37]. These facts convincingly demonstrate that the recovery expression of proteasomal subunits is transcriptionally regulated by Nrf1 that is dominantly triggered by limited proteasomal inhibition. Notably, a key to the recent debate on this issue is whether Nrf1 is proteolytically processed and consequently activated by the proteasome-dependent and -independent pathways [18, 21, 22]. This debate has indirectly reflected that the mechanism underlying dual opposing effects of proteasomal inhibitors on Nrf1-mediated proteasomal gene expression remains elusive [18, 19, 22]. Recently, proteasomal dysfunction by its inhibitors was considered to result in activation of Nrf1 (and its homologue Skn-1), *per se*, by DDI-1/2-mediated proteolytic processing of the CNC-bZIP protein [21, 22, 24, 25].

In essence, selective proteolytic processing of Nrf1, as for general degradation or specific cleavage, is dictated by topovectorial locations of both its potential degrons and cleavage sites within distinct contexts, particularly its N-terminal UBL-adjoining portions ([7, 19] and this study). For example, only upon retrotranslocation of Neh2L (aa 156-242) from the ER luminal side of membranes into extra-ER cytoplasmic compartments, this region is allowed to serve as a *bona fide* functional degron targeting Nrf1 (or its fusion proteins such as Gal4D/Nrf1^170-243^) for the cytosolic ubiquitin-proteasomal degradation. However, some incompletely-degraded proteoforms (including the N-terminal 12.5-kDa polypeptide) are enhanced by dose-dependent inhibitors of proteasomes rather than calpains. Further examinations of Nrf1^Δ1-156^, Neh2^Nrf2^:Nrf1^Δ1-156,^ Nrf2, and Nrf2^Neh2L^ have revealed that, like the Neh2 domain of Nrf2, the Neh2L of Nrf1 has a capability to target for relevant protein degradation (Figure S9A). The functionality of Neh2L may be elicited only upon retrotranslocation and dislocation into extra-luminal side of membranes, which is distinguishable from that of Neh2 at different ubiquitination status. Moreover, comparison of deletion mutants from within Neh2L-flanking or Neh5L-containing regions (to yield V5-N298^Δ261-279^-eGFP, V5-N298^Δ243-260^-eGFP and V5-N298^Δ280-298^-eGFP) unravels that a loss of the Cdc4 phosphodegron (CPD, ^267^LLSPLLT^273^) enables the resultant mutant protein (together with a major cleaved polypeptide) to be conferred with a relative slower electrophoretic mobility than equivalents examined from other two cases. In addition, the protein stability might be monitored by topological repositioning of the PEST1 degron (aa 141-170), because it enables Gal4D/Nrf1^81-181^ to be targeted for degradation within the Gal4D portion.

Intriguingly, the proteolytic processing of Gal4D/Nrf1^1-181^, by comparison with Gal4D/Nrf1^81-181^, resulted in a multiple polypeptide ladder, and similarly various ladders were found in Gal4D/Nrf1^1-298^ and also in a series of progressively NTD-lengthened deletion mutants up to yield Gal4D/Nrf1^107-298^. However, not a similar polypeptide ladder was examined in additional cases of N65-, N124- and N186-Gal4D-VP16. Therefore, it is inferable that the N-terminal 80-aa region of Nrf1 has a potent ability to act as a partial functional degron, possibly depending on the dynamic topological folding of such distinct strategic fusion proteins within and around the ER. This appears to be consistent with the notion that CRAC1/2-adjoining aa 31-80 region of Nrf1 is required for its turnover through the Hrd1-mediated ERAD pathway [7, 17, 19]. Based on these findings, together with an alignment of multiple amino acid sequences (Figure S1), it is reasoned that upon dislocation of Nrf1 (and its fusion proteins) into the extra-ER cyto/nucleoplasms, most of its NTD could also be refolded into a membrane-regulated UBL module [32, 33]. Particularly, when the essential ER-anchored region of Nrf1 is disrupted, the remaining portions of mutant proteins would be released from membranes and thus be not protected by membranes. Consequently, those refolded UBL-containing proteins might be rapidly proteolytically degraded by 26S proteasomes (Figure S9C). This assumption is also supported by additional evidence that NTD can act as a potential UBL degron targeting its fusion proteins with Nrf2 (Figure S9B) or Gal4D (Figures 2B and S2) to the proteasomal degradation pathway.

Recently, a group reported that ubiquitination of Nrf1 is required for its ensuing proteolytic cleavage besides proteasomal degradation [18, 22]. However, mapping of K/R mutagenesis has revealed that putative lysine-based ubiquitination of UBL-adjoining NTD and N298 regions is not a necessary prerequisite for the selective proteolytic processing of Nrf1 (and its fusion proteins) to yield multiple processed proteoforms (Figures 8 and S6). In addition, we have also found that either NTD- or N298-fused proteins were electrophoretically resolved with a slow mobility of higher molecular weights than that of their original full-length forms (Figures 3 and S10). This implies that these higher molecular mass proteins may be raised by protein oligomerization or aggregation, besides ubiquitination, through the N-terminal portion of Nrf1. Further experiments demonstrate that such putative oligomerization of Nrf1 is also influenced by distinct changes in redox states and pH values of changing intracellular microenvironments (Figure S10), but the underlying mechanism needs to be further elucidated.

### The retrotranslocation of Nrf1 by p97 determines its juxtamembrane proteolysis by cytosolic DDI-1 and proteasomes to remove its N-terminal portions during its maturation into an active CNC-bZP factor

Several lines of experimental evidence have been presented as illustrated for a proposed model (Figures 3H, 12A). This aims to giving a better explicit explanation of topovectorial mechanisms that control dynamic repositioning of Nrf1 by p97-driven retrotranslocation and -independent pathways into the extra-ER cyto/nucloplasmic side of membranes and the juxtamembrane proteolysis of the CNC-bZIP protein by cytosolic DDI-1 proteases and/or 26S proteasomes. This process is collectively referred to as RJP in this study. Consequently, the proteolytic processing of Nrf1 leads to generation of multiple distinct isoforms, including a mature active ∼85-kDa CNC-bZIP factor. The maturation is accompanied by the selective cleavage of within its NHB2-adjoining peptides and proteolytic processing of AD1 around PEST1 and Neh2L degrons in order to remove various lengths of the N-terminal polypeptides between ∼12.5-kDa and ∼55-kDa.

Distinct processing steps of Nrf1 are determined by dynamic repositioning of its UBL-containing NTD and AD1 into the extra-ER cyto/nucleoplasms, allowing the protein to be selectively processed by cytosolic DDI-1/2 and 26S proteasomes. This is supported by our work showing that, while p97-driven retrotranslocation is arrested by its inhibitor NMS-873, the N-linked glycoprotein of Nrf1, as well its NTD- or N298-fusion proteins, was concomitantly sequestered in the lumen of ER and hence predominantly protected by membranes against cytosolic proteolytic processing, such that yield of its processed proteoforms, including its mature protein of ∼85-kDa, was diminished. Conversely, upon recovery of p97 from its inhibitor, the luminal-arrested Nrf1 glycoprotein is immediately released by p97-driven retrotranslocation pathway to enter the extra-ER cyto/nucleoplasms. Thereafter, Nrf1 is allowed for deglycosylation and proteolytic processing by DDI-1/2 and 26S proteasomes to yield a mature active CNC-bZIP factor regulating transcriptional expression of target genes. This notion is also further evidenced by additional data obtained from DD1-deficient cells. Together with the data from other groups [18, 29], these results convincingly demonstrate that p97 executes a essential function for the retrotranslocation in the proteolytic processing of Nrf1. However, such an important role of p97 in the RJP of Nrf1 is unaffected by its client ubiquitination-deficient K/R mutants. As such being in the case, portions of a small fraction of Nrf1 (and its fusion proteins) are also subjected to their additional repositioning through p97-independent pathways, i.e. CRAC-associated cholesterol-enriched membrane translocation. This also leads to dynamic exposure of those potential degrons and cleavage sites (within its UBL-adjoining regions) to the enzymatic active center of cytosolic DDI-1 and other proteases. Consequently, the selective proteolytic cleavage of Nrf1 enables most of its C-terminal CNC-bZIP-adjoining domains to be released from membranes, whilst portions of its N-terminal membrane-bound polypeptides may be partially protected on some occasions.

Notably, the putative RJP of Nrf1 is completely distinctive from the previous known mechanisms accounting for the processing of signal peptides [40, 41] and the regulated intramembrane proteolysis (RIP) of ATF6 and SREBP1 [23] (Figure 12, A to C). Specifically, the RJP of Nrf1 is evidenced by the fact that the topovectorial processing of the ER-anchored protein in order to yield distinct isoforms occurs on the extra-ER cytoplasmic side in closer proximity to ER membranes. Conversely, none of similar putative processed N-terminal polypeptides are determined to arise from all the TM1-disrupted proteins of Nrf1 and relevant fusion proteins, though they are unable to be anchored within ER membranes so as to gain a free access to cytosolic proteases (Figures 5B and S9). This is just a matter of objective fact, supporting that the RJP of Nrf1 should occur in the vicinity of the cytoplasmic surface of ER. Further bioinformatics reveals that the RJP of Nrf1 by DDI-1/2 proteases is much likely to depend on the juxtamembranous location of its enzymatic active site. This notion is supported by additional fact that the active center of DDI-1/2 is flanked immediately by its N-terminal membrane-bound CRAC motif and its C-terminal substrate-binding regions (Figure 12E). The CRAC-adjoining region of DDI-1/2 is predicted to refold into a basic-hydrophobic amphipathic helix (Figure 12D), enabling it to lie topologically on the cytoplasmic plane of membrane lipid bilayers. By contrast, the basic-enriched substrate-binding region of DDI-1/2 and its adjacent leucine zipper domain of DDI-1 (Figure 12E) could be allowed for a putative interaction with the bZIP region of Nrf1. Whatever happens in such detail, our evidence also unravels that the proteolytic processing of endogenous Nrf1 precursors by DDI-1 and other protease leads ultimately to the generation of an active ∼85-kDa CNC-bZIP factos, as a remarkable functional consequence, to regulate target genes. Conversely, deficiency of DDI-1 results in severe impairments in the mature processing of Nrf1 and transcriptional expression of its cognate genes.

### Concluding remarks

Here, we have thoroughly elucidated topovectorial mechanisms that control dynamic movement of Nrf1 in and out of the ER lumen, as well as the selective proteolytic processing of the protein to remove distinct lengths of within its NTD (most of which is refolded as UBL) and adjacent AD1. The mature processing of Nrf1 leads to generation of an active ∼85-kDa CNC-bZIP factor, after its N-terminal portions are truncated. Amongst processed isoforms, a major N-terminal 12.5-kDa polypeptide of Nrf1 is inferable to arise from the selective protease-mediated cleavage within its NHB2-adjoining region (on the C-terminal border of putative UBL). By contrast, other longer N-terminal UBL-containing isoforms are yielded from the proteolytic processing of Nrf1 within its AD1 around three putative degrons PEST1, Neh2L and CPD. Thus, the susceptibility of Nrf1 (and its fusion proteins) to proteolytic cleavage is determined by dynamic repositioning of its NTD and AD1 from the ER luminal side of membranes into extra-ER cyto/nucleoplasmic compartments. The repositioned degrons and cleavage sites of Nrf1 within these two domains are coming into their own *bona fide* functionality, in order to enable the protein to be selectively processed by cytosolic proteases DDI-1/2 and degraded *via* 26S proteasomes. Such retrotranslocation and dislocation of Nrf1 to release from ER membranes is dominantly driven by p97-dependent pathways, but the role of p97 in the RJP of Nrf1 is unaffected by the client ubiquitination-deficient mutants of Nrf1. Furthermore, potential ubiquitination of UBL-adjoining regions in Nrf1 is not a prerequisite necessary for involvement of p97 in the selective proteolytic processing of the CNC-bZIP protein.

## EXPERIMENTAL

### Chemicals, antibodies and other reagents

All chemicals were of the highest quality commercially available. The 26S proteasomal inhibitor (MG132) and Calpain inhibitors (ALLN, CII and CP) were purchased from Sigma-Aldrich (St Louis, MO, USA). Another proteasomal inhibitor bortezomib (BTZ) were obtained from ApexBio (USA). Tunicamycin and thapsigargin were from Sangon Biotech (Shanghai, China). Cholesterol, POPC (1-palmitoyl-2-oleoyl-sn-glycero-3-phosphocholine), DPPC (1,2-di palmitoyl-sn-glycero-3-phosphocholine) and palmitoyl-sphingo-myelin, PSM (N-palmitoyl-D-erythro-sphingosyl phosphorylcholine) were from Avanti Polar Lipids (Alabaster, AL,USA). The concentrations of other different phospholipids were determined by the Bartlett method [42] and for cholesterol gravimetrically (MT5, Mettler-Toledo, Columbus, OH, USA). The primary antibodies against each tag, e.g. V5 (Invitrogen, Shanghai, China), GFP (Millipore, Tumecula, CA), Flag (Beyotime Biotechnology, Shanghai, China), Strep II (Beijing Ray Antibody Biotech, Beijing, China), VSVg (Abcam, Shanghai, China) and Ub epitope (Cell Signaling Technology, Bedford, MA, USA) were here employed, besides the first antibody against β-actin and various secondary antibodies were obtained from ZSGB-BIO (Beijing, China).

### Cell culture, transfection and Luciferase reporter assay

Experimental cells [1.5 ×10^5^, i.e. COS-1 cells and HEK293 were maintained as described [7, 19]] were seeded into each of 35-mm culture dishes. After the cells reach 60% or 80% confluence, they were transfected with either siRNA or expression constructs in a mixture with the Lipofectamine 3000 (Invitrogen) for 7-h. In brief, COS-1 cells were co-transfected with a reporter gene *P_TK_UAS/Gal4-Luc,* along with Renilla luciferase reporter (*pRL-CMV*) or *pcDNA4/HisMax/lacZ* encoding β-galactosidase (β-gal), each of which served as an internal control for transfection efficiency, and together with expression constructs for Gal4D-Nrf1, Nrf1-Gal4D-Vp16 or their mutants. At 16-h after transfection, the reporter gene activity was measured by using dual luciferase reporter assay (E1910 from Promega) according to the manufacturer’s instruction.

### Plasmids and siRNAs

Each sequence of cDNAs encoding V5, Strep II and VSVg was inserted into the XhoI/KpnI sites of the *peGFP-N2* vector, in which the translation start codon ATG of eGFP was mutated to CTG insofar as to abolish generation of free GFP. Then sandwiched fusion proteins V5-Nrf1-GFP, StrepII-Nrf1-GFP and VSVg-Nrf1-GFP were engineered by inserting indicated nucleotide sequences encoding Nrf1, its truncated or mutants into the KpnI/XbaI sites. A series of expression constructs for Gal4D/Nrf1 and its truncated mutants were created by ligating various lengths of cDNA fragments encoding N-terminal portions of Nrf1, to the 3′-end of Gal4D cDNA contained in the *pcDNA3.1 Gal4D-V5*, through the BamHI/EcoRI site. Another series of Nrf1-Gal4D-Vp16 plasmids were also constructed by inserting various lengths of codons for N65, N124, N186, N242 and N298 (each of which contains the N-terminal 65, 124, 186, 242 and 298 aa of Nrf1) into the KpnI/XbaI sites of the *pcDNA3.1/Gal4D-Vp16/V5His B*, which was created by subcloning cDNA of Gal4D-Vp16 (Clontech) into the XbaI/SacII site of *pcDNA3.1/V5His B*. In addition, several expression constructs for Nrf2 and relevant chimeric proteins were described previously [11]. All oligonucleotide primers were synthesized by TSINGKE Biological Technology (Chengdu, China), and the fidelity of all the cDNA products used were confirmed by sequencing. Moreover, small interfering RNAs (siRNAs) targeting against the human DDI-1 or DDI-2 coding regions were designed (siDDI-1#1, 5′-CAUGAAUAUAGCGAUAGAA-3′; #2, 5′-GUCAUGGAUUCAGGACGAA-3′; #3, 5′-CAAGUGAC GAUGCUCUACA-3′; siDDI-2#4, 5′-GUGCCCAGAUGACUAUCAU-3′; #5, 5′-GAGAUAUGUUGCUGGCCAA-3′; and #6, 5′-CUCAUCUCCUGGAGAAA U-3′), which were biosynthesized by BIOTEND (Shanghai, China). Another scrambled sequence 5′-UUCUCCGAACGUGUCACG-3′ (without bioinformatic homology with any genes analyzed in Genbank) was employed as an internal negative control.

### Establishment of DDI-1-specific knockout cell line

The CRISPR/Cas9-mediated genome-editing system was employed to establish a DDI-1-specific knockout cell line on the base of HEK293 (thus called HEK293:DDI-1^KD^). The sequence of single guide RNA (gRNA) for targeting DDI-1 specifically (5′-CACCGTGTACTGCGTGCGGA-3′) was designed through the CRISPR-direct (http://crispr.dbcls.jp/) and then inserted into the pCAG-T7-cas9+gRNA-pgk-Puro-T2A-GFP vector (Viewsolid Biotech, Beijing, China). Then, the positive clones of HEK293 cells transfected with the gRNA-engineered vector were selected by their resistance to puromycin (2.5 mg/mL) and confirmed by Western blotting, before being experimented.

### Quantitative real-time PCR analysis

Either HEK293 or HEK293:DDI-1^KD^ cell lines that had been transfected with indicated siRNAs or not were subjected to RNA isolation by using the RNAsimple Kit (Tiangen Biotech CO. Beijing, China). Subsequently, 500 ng of total RNAs was added in a reverse transcription reaction to generate the first strand of cDNA (using Revert Aid First Strand Synthesis Kit from Thermo). The synthesized cDNA was used as the template for qPCR, together with the GoTaq® qPCR Master Mix (from Promega), which was deactivated at 95°C for 10 min, and then amplified by 40 reaction cycles of 15 s at 95°C and 30 s at 59°C. The final melting curve was added to examine the quality of amplification, whereas β-actin or RPL13A mRNA levels served as an internal control. All the forward and reverse primers of indicated genes were shown in Table S1.

### Western blotting

COS-1 cells were transfected with distinct expression constructs for 7 h, and then allowed for a recovery culture for 16 h in a fresh DMEM. The cells were treated with indicated chemicals for 4 h before being harvested in denatured lysis buffer (0.5% SDS, 0.04 mol/L DTT, pH 7.5, containing 1 tablet of cOmplete protease inhibitor EASYpacks per 10 ml buffer). Immediately, the lysates were denatured at 100°C for 10 min, sonicated sufficiently, and then diluted with 3× loading buffer (187.5 mmol/L Tris-HCl, pH 6.8, 6% SDS, 30% Glycerol, 150 mmol/L DTT, 0.3% Bromphenol Blue) before re-boiled at 100°C for 5 min. Thereafter, equal amounts of protein extracts were subjected to separation by SDS-PAGE or LDS-NuPAGE. The resolved proteins were transferred onto the PVDF membranes, followed by blockage by incubation with 5% non-fat milk for 1 h, before overnight incubation with each of the primary antibodies at 4°C. On some occasions, some blotted membranes were rinsed for 30 min with stripping buffer before being re-probed with an additional primary antibody against β-actin, as an internal control, to verify equal loading of proteins in each of electrophoretic wells. The PageRuler^™^ Prestained protein ladder (Thermo, catalog No. 26616) was used to identify molecular masses of standard proteins separated during electrophoresis.

### Purification of Nrf1 and its CRAC peptides

Expression constructs for the mouse full-length Nrf1 (aa 1-741) and its smaller isoform Nrf1β (aa 292-741) were described previously [35, 36]. Briefly, COS-1 cells, that have reached 70% confluence (7**×**10^6^ cells), were transfected by electroporation of 10 μg DNA for these C-terminally 6**×**His-tagged expression constructs for Nrf1 or Nrf1β in OPTIMEM. The cells were grown for 24-h in DMEM and lysed by freeze-thawing cycles. Then, interested proteins were purified with affinity chromatography [43]. Furthermore, two 28-aa peptides CRAC^F78W^ and CRAC^F78W+Y65/77S^ of Nrf1 were commercially synthesized, and purified (>85% purity) by Genscript (Piscataway, NJ), before both were dissolved in phosphate buffered saline with 10% glycerol.

### Intrinsic tryptophan emission measurements

Putative interactions of the CRAC peptides with lipid membranes were here studied by monitoring the tryptophan fluorescence emission spectra upon addition of vesicles. The dry lipids were dissolved in phosphate buffered saline and then sonicated with a probe sonifier (Branson 250). The size of the small vesicles has been measured to be ∼45 nm in diameter, by using a light scattering instrument (Malvern Zetasizer Nano-SZEN1600, Malvern Instruments, Worcestershire, UK). The tryptophan fluorescence of both CRAC^F78W^ and the CRAC^F78W+Y65/77S^ peptides (2.0 μmol/L) was measured before and after addition of POPC, DPPC or PSM vesicles, or each of their combinations with 25 mol/L % cholesterol to a final lipid concentration at 0.5 mmol/L (with a 250:1 ratio of lipid to peptide). The measure of tryptophan fluorescence (at 295 nm excitation) was conducted between 310 and 450 nm at 37°C on a Cary Eclipse (Varian, Palo Alto, CA), by using spectrofluorometer with excitation bandwidth at 5 nm and emission bandwidth at 5 nm. The emission intensities are corrected for the increasing light scattering of the vesicles added.

### Surface plasmone resonance (SPR) spectroscopy

Membrane-peptide interactions of the CRAC peptides with distinct vesicles were also examined at 23°C with the BioNavisSPR Navi 200 instrument (BioNavis Ltd, Ylöjärvi, Finland). Such vesicles including POPC, DPPC or PSM with or without 15 mol% cholesterol in PBS were immobilized on a gold sensor chip. The sensor chip surface consists of acarboxymethylated dextran matrix with preimmobilized lipophilic groups (C10) and the vesicles can be captured at the surface non-covalently. In experimentation, freshly prepared vesicles were immobilized on the sensor chip at a flow rate of 20 µl/min in PBS (pH 7.4). In order to remove any multilamellar aggregates from the surface, 200 µl of NaOH (50 mmol/L) was injected at a flow rate of 20 µl/min; this resulted in a stable baseline corresponding to the lipid monolayer linked to the chip surface. In addition, 200 µl of BSA (0.1 mg/µl in PBS) as a negative control was injected to confirm complete coverage of the non-specific binding sites. The peptide binding was measured by observing the changes in the SPR angle as 200 µl of the peptide analyte flowed over the immobilized vesicles with a flow rate of 20 µl/min. The sensor chip was regenerated using 200 µl of 20 mmol/L CHAPS at a flow rate of 20 µl/min for 2 times. The response was monitored as a function of time at 23°C. The experiments were conducted in triplicate and the bindings were repeatable at least twice.

### Statistical analysis

Statistical significance of changes in Gal4/*UAS*-driven reporter gene activity and other gene expression was determined using either the Student’s *t*-test or Multiple Analysis of Variations (MANOVA). The data are shown as a fold change (mean ± S.D), each of which represents at least 3 independent experiments that were each performed triplicate.

## Acknowledgments

The study was supported by the National Natural Science Foundation of China (key programs 91129703 & 91429305) awarded to Prof. Yiguo Zhang (University of Chongqing, China), and in part funded by Chongqing University postgraduates′ innovation project (No. CYB15024) awarded to Mr. Lu Qiu. This work was also partially funded by Sigrid Jusélius Foundation and Magnus Ehrnrooth Foundation awarded to Dr. Peter Mattjus (Åbo Akademi University, Finland).

## Author contributions

Y.X. constructed most of the plasmids, performed all Western blotting except those indicated, measured some lucifurase assays, collected the data, and prepared drafts of most figures. J.H. fulfilled the purification of Nrf1 and CRAC peptides, together with SPR spectroscopy and intrinsic tryptophan emission measurement. Z.F. carried out the live-cell imaging experiments. S.H. established HEK293:DDI-1^KD^ cell lines and completed relevant work. M.W. and L.Q. had done the qPCR and repeat lucifurase assays. Z.Z. created some expression constructs and finished bioinformatic analysis. P.M. together with Y.Z. analyzed and discussed the data for Figure 1, and helped to revise this manuscript. Y.Z. designed this study, constructed an essential portion of expression plasmids, measured lucifurase assays, analyzed all the data, prepared all figures, wrote and revised the paper.

