## Supplementary Materials for "Topovectorial mechanisms control the juxtamembrane proteolytic processing of Nrf1 to remove its N-terminal polypeptides during maturation of the CNC-bZIP factor"

### Figure S1

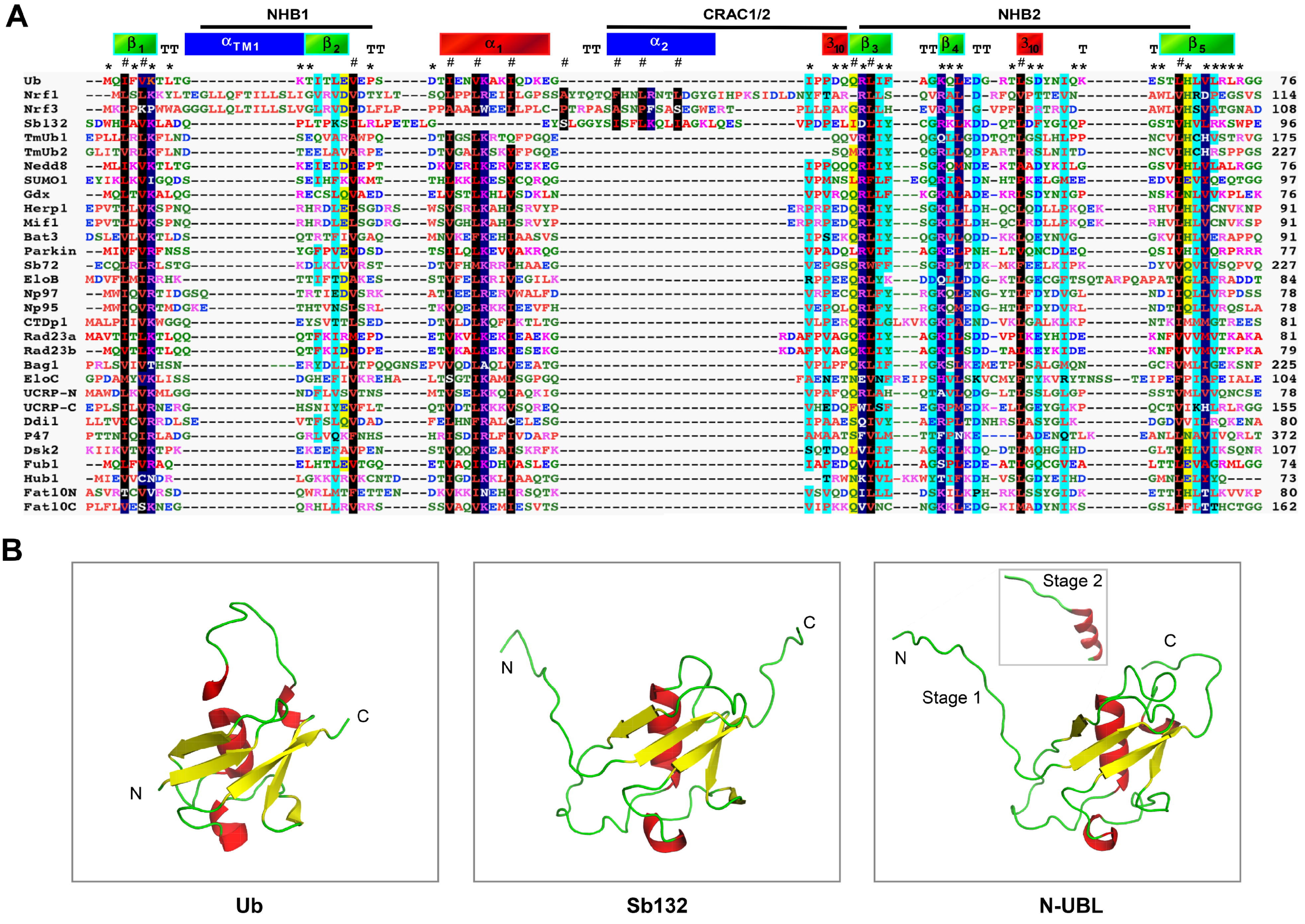


**Figure S1. Multiple amino acid sequence alignment of NTD in Nrf1 with 30 distinct UBL proteins.**
**(A)** Within NTD of mouse Nrf1, its amino acids 1 to 114 [covering three major motifs NHB1, CRACs and NHB2] were aligned with similar regions of known ubiqintin-like (UBL) proteins using the T-Coffee. Besides those structurally conserved regions within UBL domains, the N-terminal 114-aa region of Nrf1 also contains a unique TM1 α-helix ($\alpha_{TM1}$) within the NHB1 signal sequence and additional $\alpha_2$ (similar to Sb132 only) within CRACs; this region of Nrf1 could be thereby allowed to fold into a membrane-regulated UBL domain only after being released from the confined membranes. The symbol (#) indicates those residues buried in the globular β-grasp Ub superfold as ββαββαβ, the star (*) represents some residues in the hot-spots, and the letter T shows the secondary structural turns. The consensus residues are placed on distinct shaded backgrounds. **(B)** A proposed three-dimensional structure of the N-terminal UBL (N-UBL, *right*) of Nrf1 was modeled by the PyMOL program and Discovery Studio 2.5 software on the template of homology with SB132 (ID: 1X1M from PDB, *middle*) or Ub (ID: 2GBM, *left* ). It is important to note that such structural variations are caused by membrane-associated folding and membrane-released refolding of both the N-terminal amphiphathic TM1 α-helix ($\alpha_{TM1}$) and $\alpha_2$ regions, because they could span across amphipathic membrane lipids or lie flat on the plane of the membrane bilayers, in addition to a possibility that they also exist as a loop in a non-liposoluble microenvironments. The abbreviations of UBL proteins used herein include: Bag1, Bcl-2 binding athanogene-1; Bat3, HLA-B-associated transcript 3; CTDp1, UBL domain- containing CTD phosphatase 1; DDI-1 (and also Ddi1), DNA-damage inducible protein homolog 1; Dsk2, dominant suppressor of kar1 (homologous with human proteasome ligase interaction component 1, hPLIC); EloB, transcription elongation factor B polypeptide 2 (Elongin B, a component of the multi-subunit VHL E3 complex); EloC, transcription elongation factor B polypeptide 1 p15; Gdx, UBL protein from chromosome Xq28; Fat10, F-adjacent transcript-10 comprising two UBL domains (called ubiquitin D); Fub1, failure of ureteric bud invasion protein; Herp1, homocysteine-responsive ER-resident UBL domain member 1 protein; Hub1, homologous to ubiquitin 1, known as UBL5 in mammals); Mif1, methyl methanesulfonate-inducible fragment protein 1 (also called Herp1); Nedd8, neural precursor cell expressed developmentally down-regulated protein 8 (also called RUB, related to ubiquitin); Np97 and Np95, nuclear zinc finger protein 97 and 95, respectively; parkin, an E3 ligase containing an N-terminal UBL; p47, NSFL1 (p97) cofactor; Rad23, radiation-sensitive mutant 23 of UV excision repair proteins, and its human homolog called hHR23 (which forms a complex with PNGase for coupling of deglycation with degradation); Sb132, a bone marrow stromal cell UBL protein (BMSC-UBP); Sb72, dendritic cell derived UBL protein DC-Ubp; SUMO1, small ubiquitin-related modifier-1; TmUb, transmembrane UBL protein; and UCRP, ubiquitin cross-reactive protein precursor, also called interferon-stimulated gene product 15-kDa (ISG15).

### Figure S2

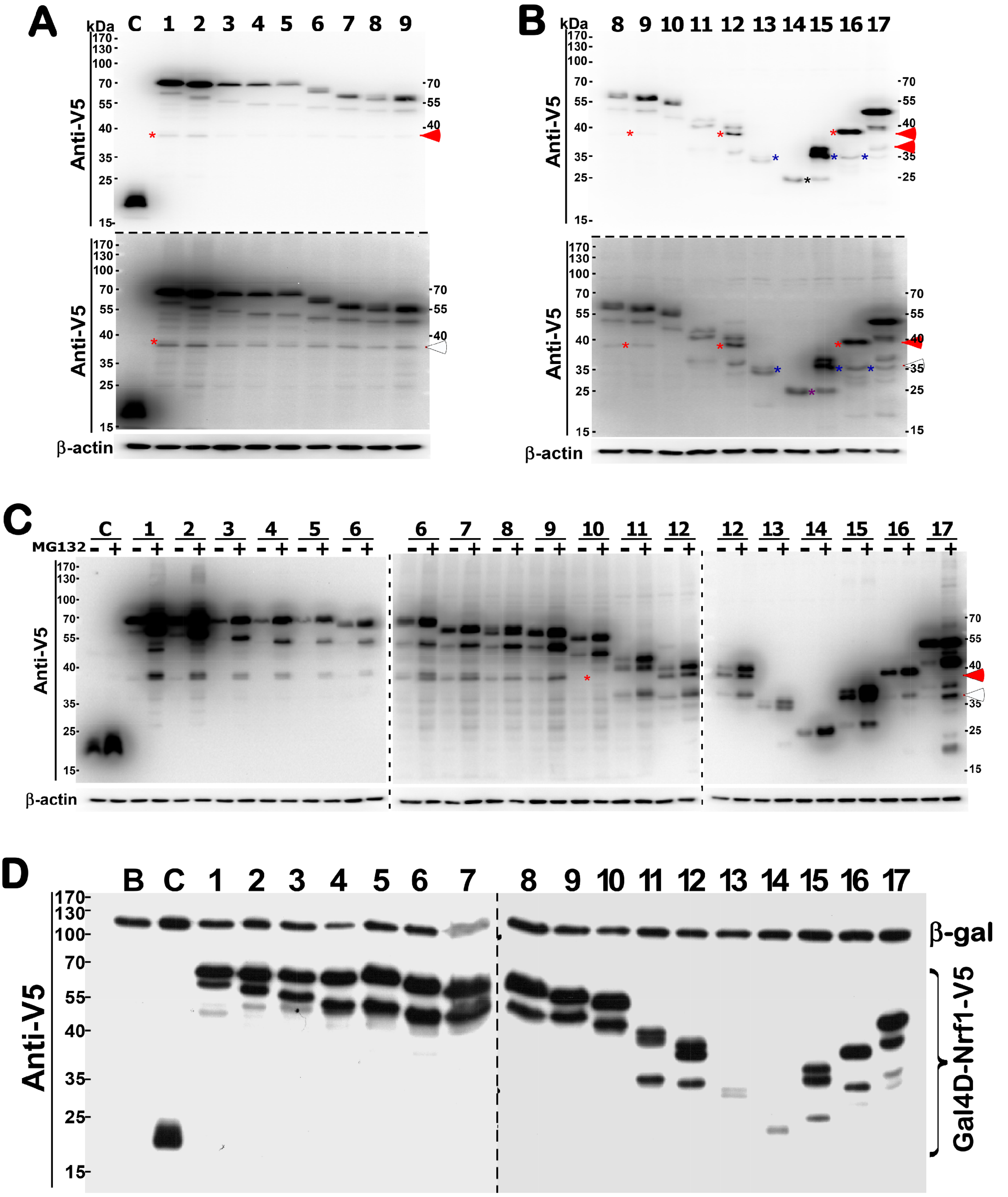


**Figure S2. Multiple proteoforms arising from the NTD and AD1 portions of Nrf1 fusion protein with Gal4D.**
(A to C) COS-1 cells that had been transfected with each (2 μg of DNA) of indicated expression constructs were treated with MG132 (+ 5 μmol/L) or not (–) for 4 h before being harvested in denatured lysis buffer. Equal amounts of proteins in total lysates were separated by SDS-PAGE containing 10% polyacrylamide, and then visualized by Western blotting with antibodies against V5 tagged C-terminally, whereas β-actin served as the protein-loading control. Similar images acquired from the exposure of Western blots for a relative short time are shown on the top (***A***,***B***). (D) These transfected cell lysates were also subjected to protein isolation by SDS-PAGE containing 8% polyacrylamide, followed by Western blotting with anti-V5 antibody, with β-gal used as an internal protein-loading control. Of note, the work as shown in (***D***) was carried out in Scotland, UK, whereas other repeat experiments (***A*** to ***C***) finished in China. A resultant nuance in these results presented should be attributable to different lysis buffer and/or experimental settings in between two countries.

### Figure S3

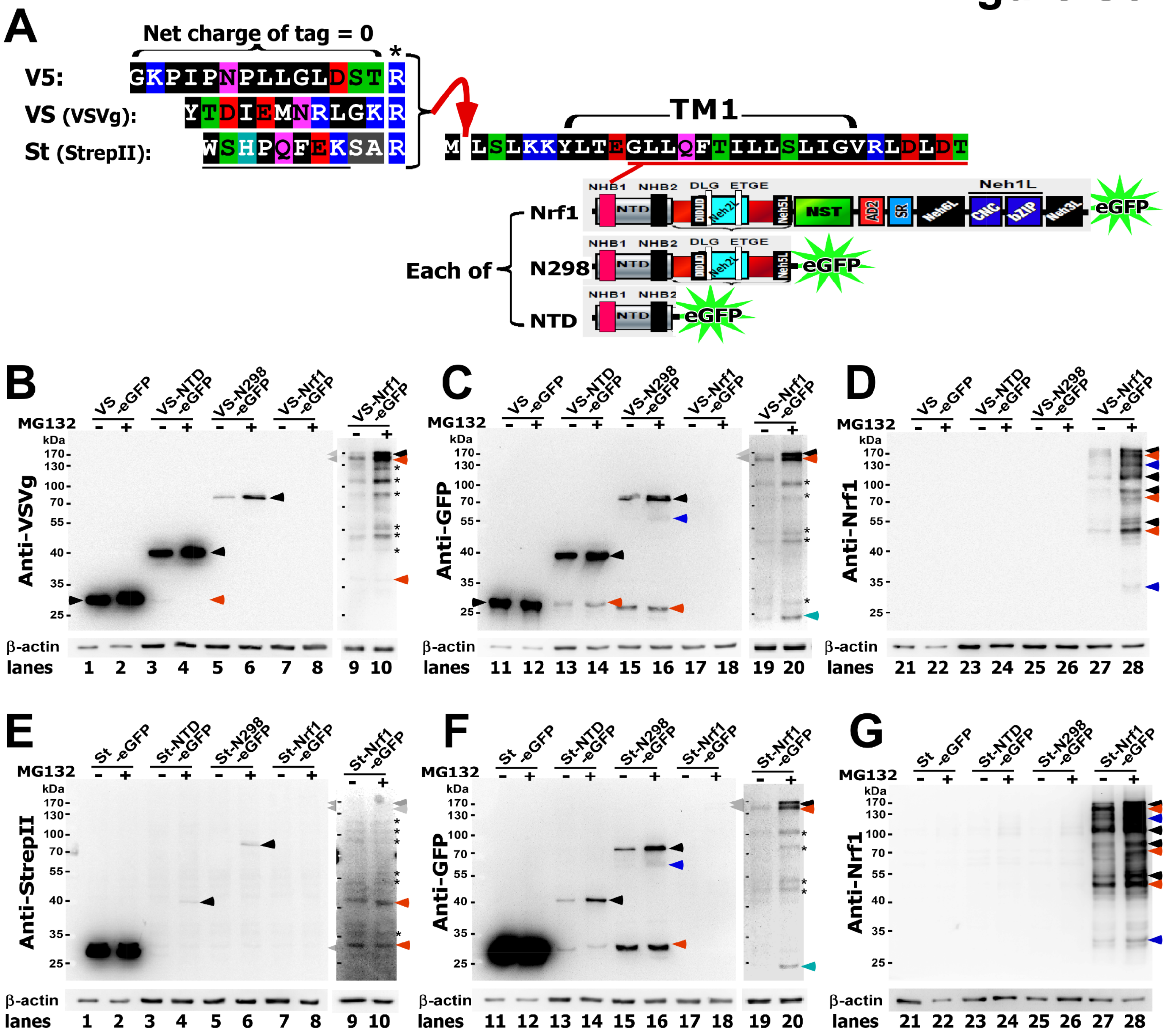


**Figure S3. Multiple polypeptides arising from distinct doubly-tagged Nrf1, its NTD and N298 proteins.**
(**A**) Schematic of the full-length Nrf1, its NTD and N298 within distinct doubly-tagged fusion contexts. Each of net zero charged tags such as V5, VSVg (VS) and StrepII (St) was attached to the N-terminal end of Nrf1 or its truncated mutants, and each C-terminal end of these fusion proteins was also associated with an eGFP ectope. (**B** to **G**) Each of expression constructs for indicated fusion proteins was transfected into COS-1 cells and then allowed for a 16-h recovery before being treated with MG132 (+ 5 µmol/L) or not (–) for 4 h. Subsequently, total lysates were subjected to identification of multiple polypeptides by SDS-PAGE containing 10% polyacrylamide and immunoblotting with antibodies against each of tags VSVg (***B***), StrepII (E) GFP (***C,F***) or Nrf1 (***D,G***). In addition, samples expressing VSVg-Nrf1-eGFP or StrepII-Nrf1-eGFP were repeated and then visualized, after Western blots were exposed for a relative longer time as shown (on *lanes 9, 10, 19 and 20*).

### Figure S4

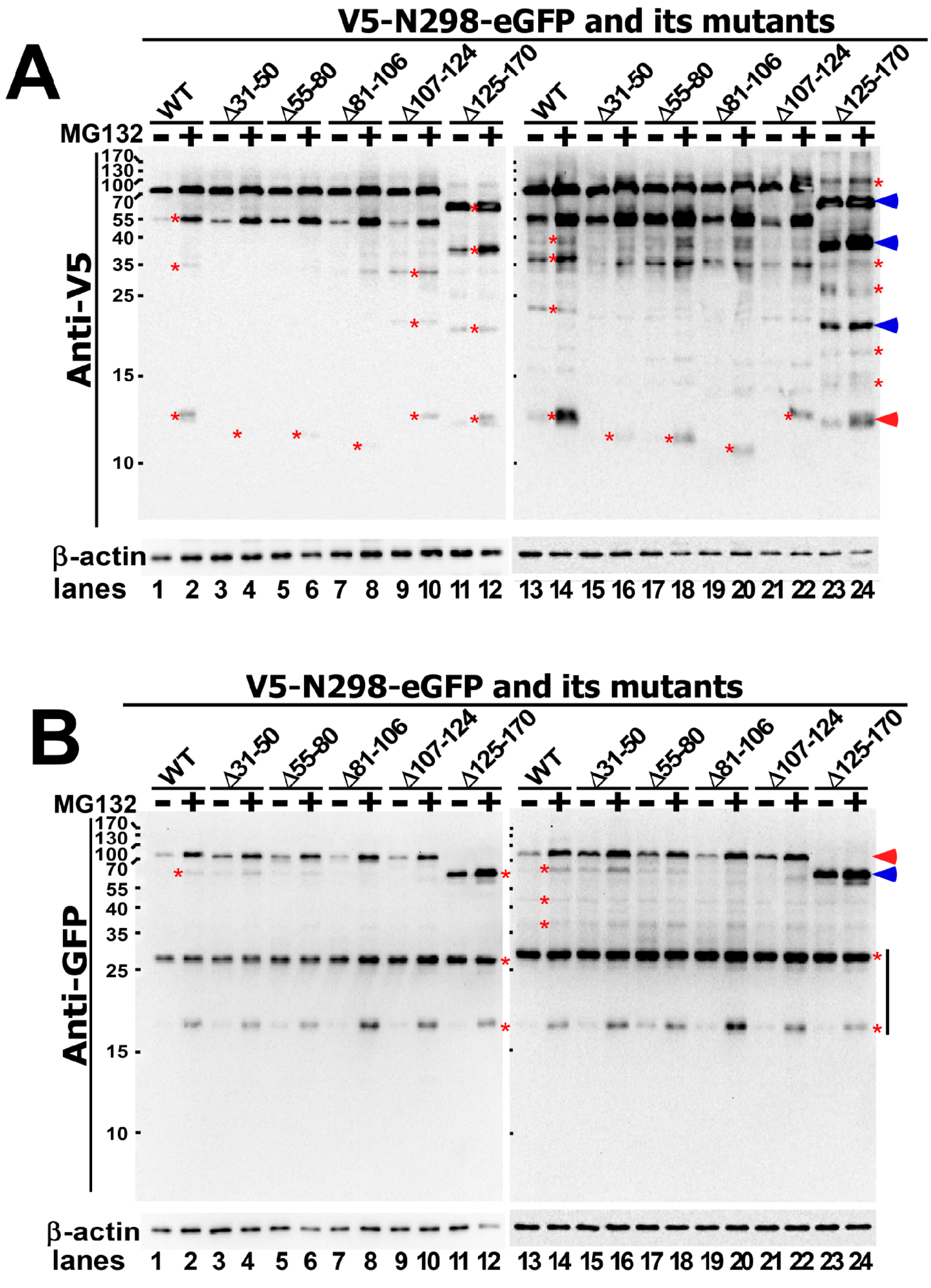


**Figure S4. Distinct effects of CRAC-adjoining peptides on the processing of V5-N298-eGFP.**

COS-1 cells expressing V5-N298-eGFP and its mutants were treated with MG132 (+ 5 μmol/L) or not (–) for 4 h before being harvested. Equal amounts of sample proteins were loaded and analyzed by Western blotting with antibodies against V5 (***A***) or GFP (***B***), as described in the legend of main Figure 5 (B and C).

### Figure S5

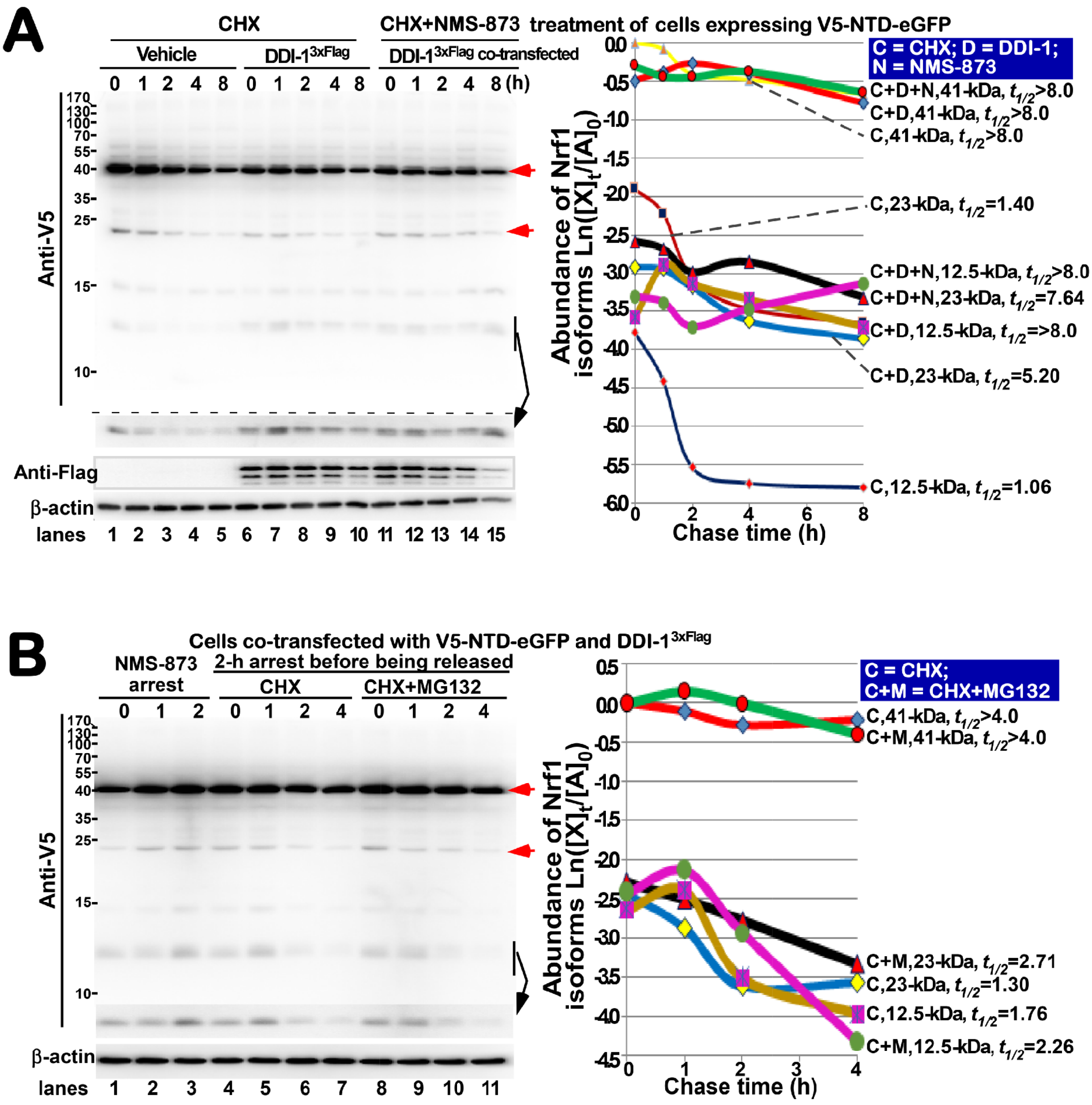


**Figure S5. Selective proteolytic processing of V5-NTD-eGFP by DDI-1 and proteasomes was monitored by p97-driven retrotranlocation and -independent pathways.**

COS-1 cells that had been transfected with an expression construct for V5-NTD-eGFP alone or in combination with DDI-1$^{3xFlag}$, were treated with NMS-873 (10 μmol/L) and/or CHX (50 μg/ml) (***A***), or in combination with MG132 (5 μmol/L) (***B***) for indicated lengths of time, before being harvested. Equal amounts of proteins of total lysates were separated by SDS-PAGE containing 15% polyacrylamide, and visualized by Western blotting with V5 antibody. DDI-1 was identified by its Flag antibody, whilst β-actin served as an internal protein-loading control. The intensity of three major polypeptide bands *arrowed* on each gels was quantified and shown graphically in the *right panels* with distinct $t_{1/2}$ values estimated, respectively.

### Figure S6

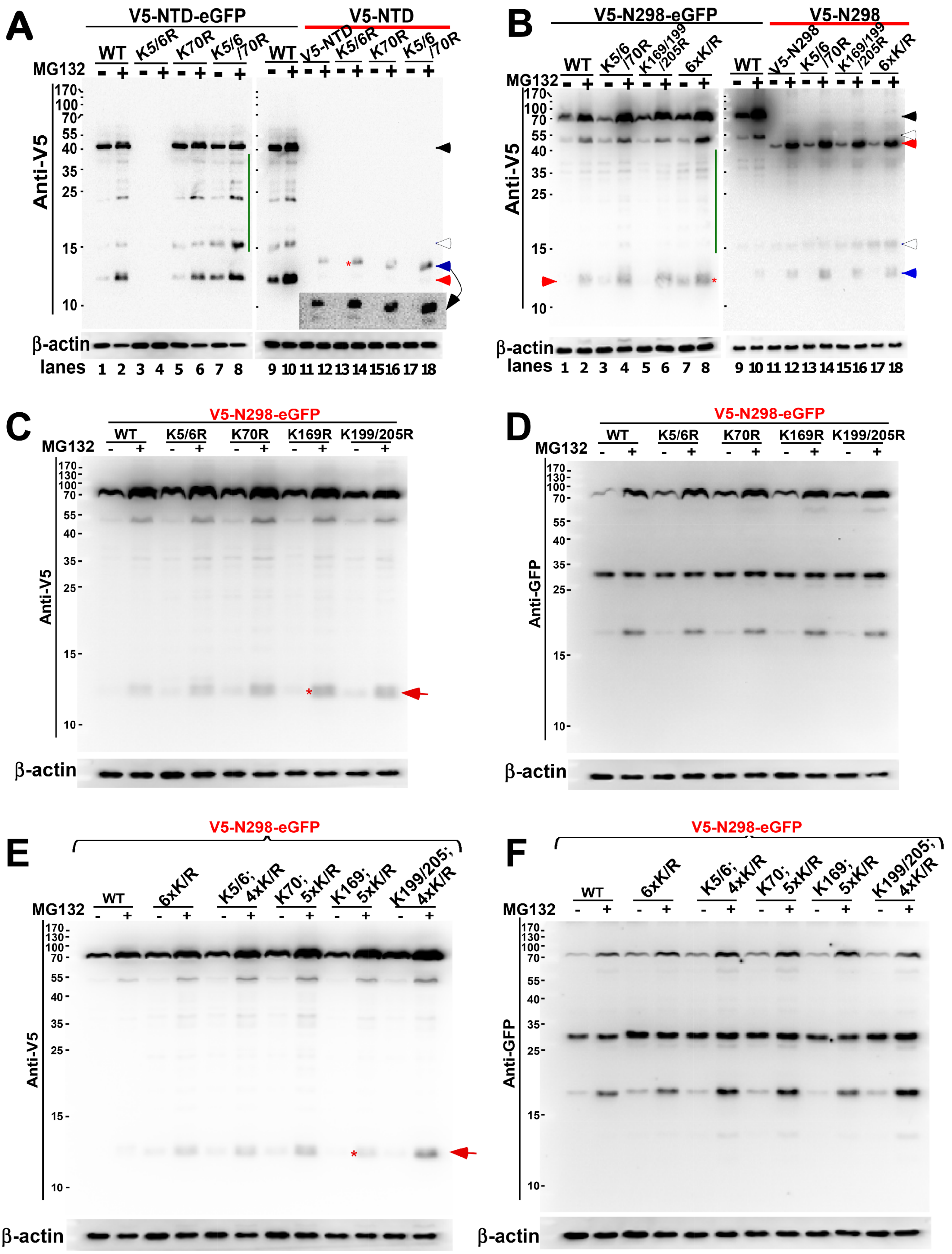


**Figure S6. No effects of all K/R mutations on the putative proteolytic processing of relevant chimeric proteins.**
**(A,B)** A series of expression constructs for single, double or triple lysine-to-arginine (K/R) mutants of the NTD (***A***) or N298 portion (***B***) of Nrf1, which were attached to eGFP (*left panel*) or without eGFP tagged (*right panel*), were transfected into COS-1 cells. They were treated MG132 (+ 5 μmol/L) or not (–) and examined by Western blotting with V5 antibodies. **(C to F)** Each of expression constructs for the indicated K/R mutants of V5-N298-eGFP (aiming to totally or partially prevent the putative lysine-based ubquintination of the N298 portion) was transfected into COS-1 cells for 7 h and then treated with MG132 (+ 5 μmol/L) or not (–) for 4 h, before being harvested in denatured lysis buffer. Equal amounts of protein samples were subjected to separation by SDS-PAGE containing 15% polyacrylamide and identification by Western blotting with distinct antibodies against V5 (***C,E***) or GFP (***D,F***). β-actin served as an internal protein-loading control. The data shown were representative of at least three independent experiments.

### Figure S7

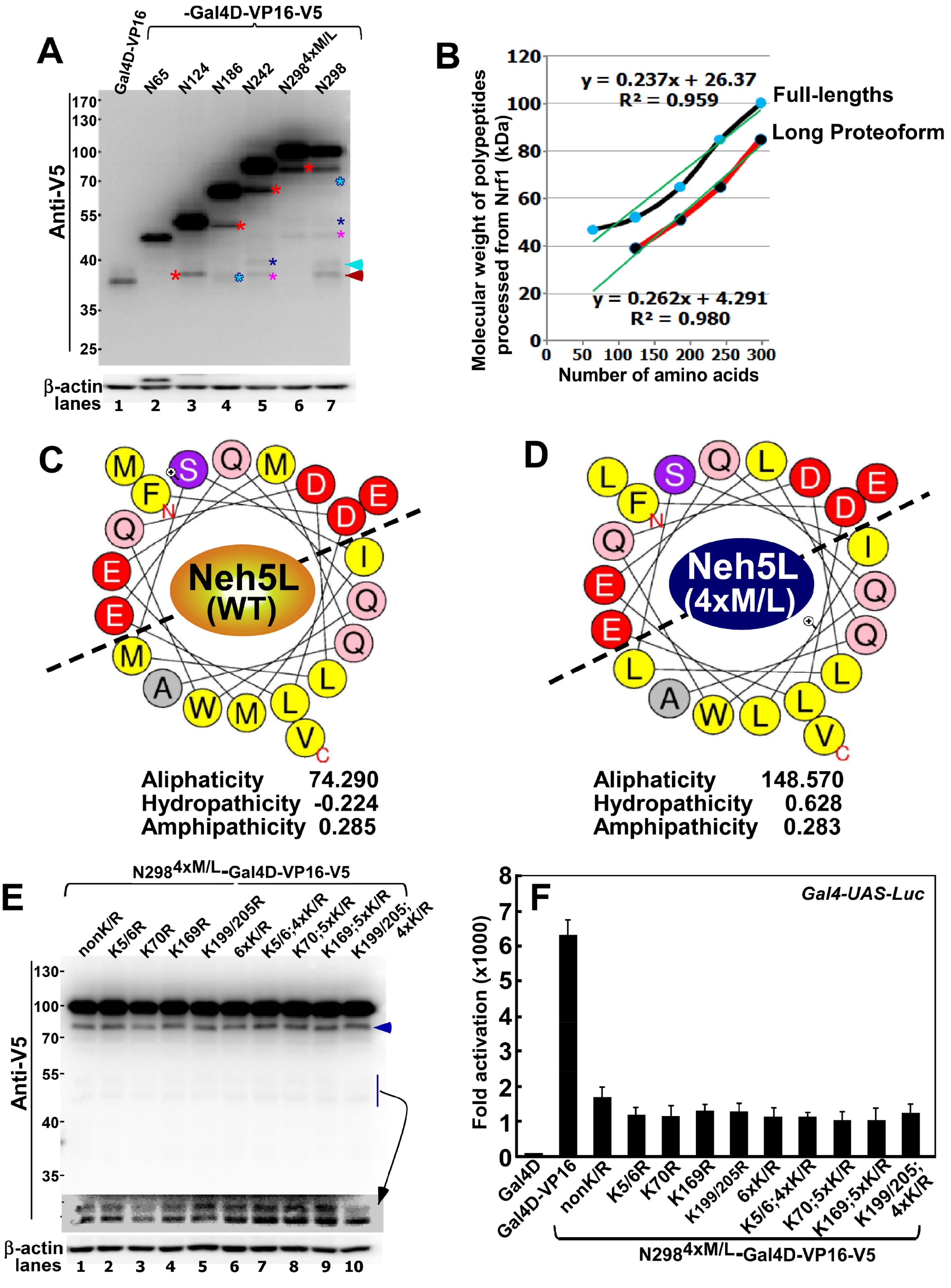


**Figure S7. Different proteoforms arising from V5-N298-eGFP and other Gal4D-VP16 fusion proteins.**

(**A**) A series of TM1-containing fusion proteins with the Gal4D-VP16 chimeras, that were allowed for expression in COS-1 cells, were visualized by Western blotting with V5 antibody. Different lengths of interested polypeptides arising from the putative proteolytic processing of these fusion proteins were indicated by *asterisk* (*) and *arrow* (←). (**B**) The number of the N-terminal amino acids of Nrf1 within distinct fusion contexts seems to be directly proportional to distinct molecular weights estimated of intact polypeptides and their processed major longer proteoforms, both of which are shown in two roughly parallel lines. (**C**) A putative interaction of wild-type Neh5L (with its core amino acid sequence $^{279}$FDLEQQWQD LMSIMEMQAMEV$^{299}$) with amphipathic membrane bilayers enables it to topologically fold into an acidic-hydrophobic amphipathic helix, which could lie flat on the plane of membrane and if required, spin or flip across membranes. (**D**) a mutant sequence ($^{279}$FDLEQQWQDL_L_SI_L_E_L_QA_L_EV$^{299}$) of the core Neh5L subdomain, in which all four original methionines were mutated into leucines (*underlined* ) insomuch as to abolish the in-frame translation of Nrf1, could be allowed to topologically fold into another acidic-hydrophobic amphipathic helix with more hydropathicity and aliphaticity than those predicted from the wild-type. (**E,F**) Each of expression constructs for indicated K/R mutants, that had been established on the base of N298$^{4xM/L}$-Gal4D-VP16 (retaining additional mutation of all four methionines into leucines within its Neh5L region), alone or in combination with $P_{TK}$*UAS/Gal4*-Luc and *Renilla* reporters, were co-transfected into COS-1 cells, in order to assay relevant protein expression (***E***) and Gal4/UAS-driven reporter gene activity (***F***).

### Figure S8

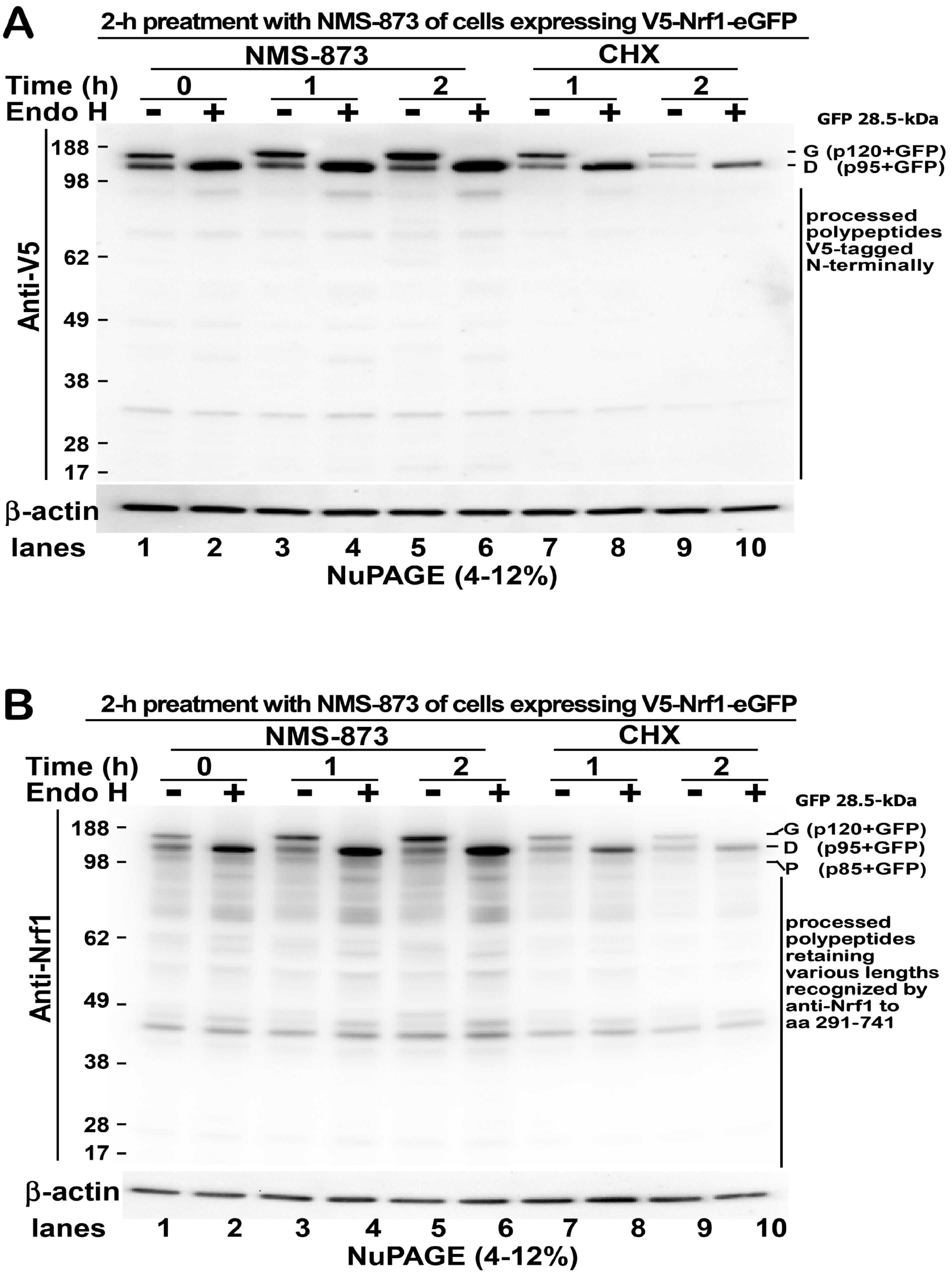


**Figure S8. Distinct isoforms arising from deglycosylation reactions with lysates of cells expressing V5-Nrf1-eGFP.**

An expression construct for V5-Nrf1-eGFP (in which the full-length Nrf1 is fused with the N-terminal V5 tag and the C-terminal eGFP) were transfected into COS-1 cells for 7 h, and then allowed for a 16-h recovery from transfection. Subsequently, the cells were pretreated with NMS-873 (10 μmol/L) for 2 h and the transferred in another fresh medium containing either NMS-873 (10 μmol/L) or CHX (50 μg/ml) to continue the culture for distinct lengths of time, before being harvested in denatured lysis buffer. The total lysates were allowed for deglycosylation reactions with Endo H for 4 h at 37℃, prior to being resolved by 4-12% NuPAGE gels and visualized by Western blotting with antibodies against Nrf1 (***B***) or its N-terminally-tagged V5 ectope (***A***). The electrophoretic locations of the glycoprotein, deglycoprotein and processed protein were indicated by single letters *G, D* and *P,* respectively.

### Figure S9

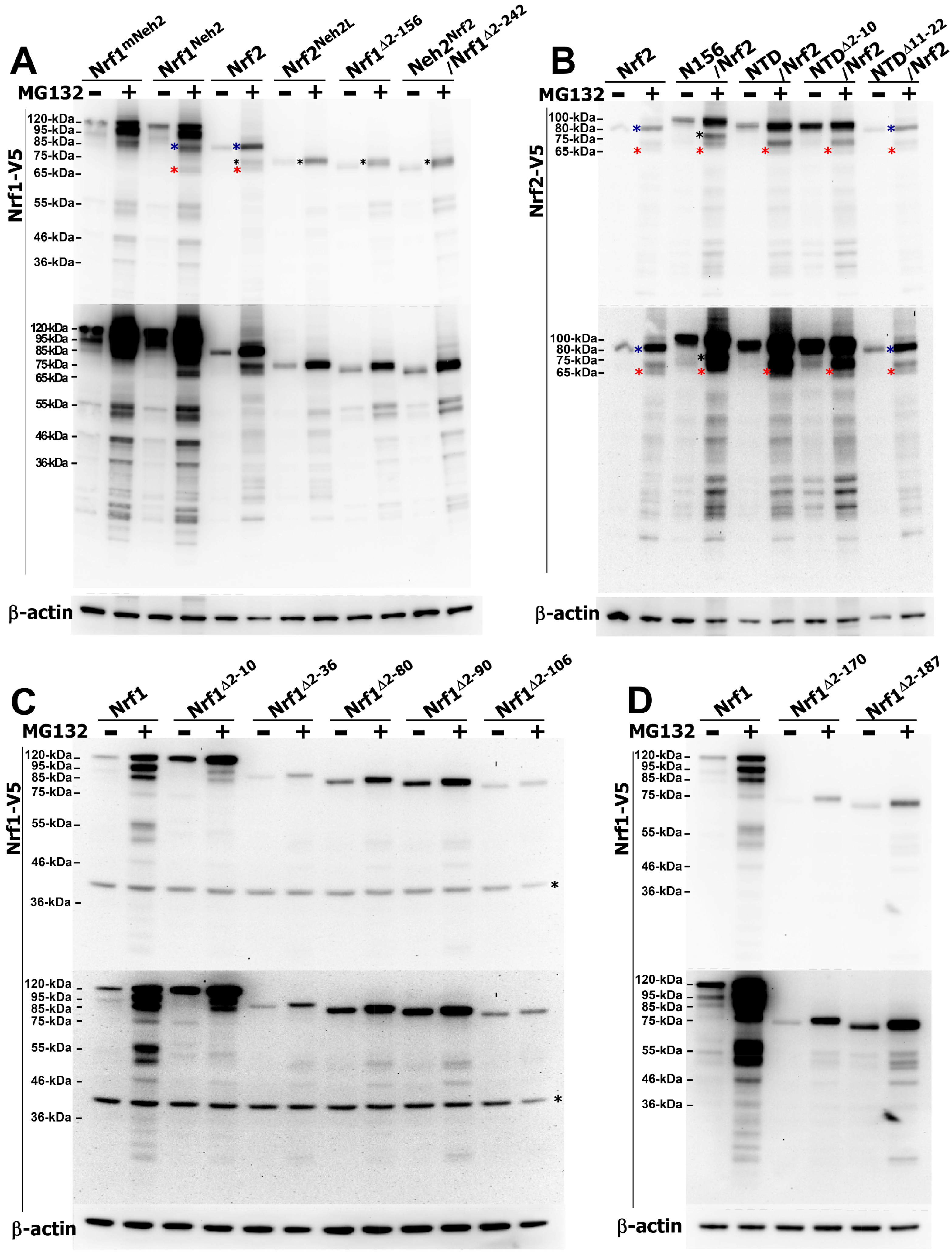


**Figure S9. Identification of functional NTD and Neh2L regions contributing to protein stability.**

(**A, B**) COS-1 cells were transfected for 7-h with each of expression constructs for $Nrf1^{Neh2}$ and $Nrf1^{mNeh2}$ [both contain $Neh2^{Nrf2}$ and mimicked Neh2 (i.e. mNeh2), respectively], $Nrf2^{Neh2L}$ (containing $Neh2L^{Nrf1}$), $Neh2^{Nrf2}/Nrf1^{\Delta2\text{-}242}$, or other indicated chimeric proteins [as described in the reference by Zhang, Y., Ren, Y., Li, S., and Hayes, J. (2014). Transcription factor Nrf1 is topologically repartitioned across membranes to enable target gene transactivation through its acidic glucose-responsive domains. PLoS ONE *9*, e93456], and then allowed for a 16-h recovery, before being treated with MG132 (+ 5 μmol/L) or not (–) for additional 4-h and subsequently were harvested in denatured lysis buffer. The total lysates were separated by 4%-12% NuPAGE gel and visualized with immunoblotting with V5 antibody. As compared with wild-type Nrf1 pattern, the proteoform expression of $Nrf1^{mNeh2}$ appeared to be unaffected by mNeh2 (A), whereas Neh2 enabled the chimeric $Nrf1^{Neh2}$ to express an extra ~65-kDa polypeptide similar to the equivalent derived from wild-type Nrf2 in the presence of MG132. In fact the full-length Nrf2 protein was exhibited to be ~85-kDa, but substitution of its Neh2 domain with the equal length of Neh2L from Nrf1 gave rise to a single $Nrf2^{Neh2L}$ chimeric polypeptide of ~75-kDa, which is similar to the molecular masses of $Nrf1^{\Delta2\text{-}156}$ and $Neh2^{Nrf2}/Nrf1^{\Delta2\text{-}242}$. These indicate that Neh2, but not Neh2L, enables Nrf2 to display a high molecular mass of ~85-kDa that is attributed to ubiquitination of this domain dependent on their chimeric contexts, and also triggers the putative proteolytic processing of its cognate Nrf2 or chimeric $Nrf1^{Neh2}$ to yield ~65/75-kDa degraded proteoforms. More intriguingly, the N-terminal portions (N156, NTD, $NTD^{\Delta2\text{-}11}$) of Nrf1 was predicated to play a role for the membrane-regulated UBL domain (Fig. S1), such that the abundance of ~65/75-kDa degraded proteoforms was increased in MG132-treated cells that had been transfected with a expression construct for relevant chimeric N156/Nrf2, NTD/Nrf2 or $NTD^{\Delta2\text{-}11}$/Nrf2. Within such chimeric protein, Nrf2 cannot be glycosylated *per se*. (**C, D**) In parallel experiments, total lysates of COS-1 cells that had been transfected with an expression construct for Nrf1 or its N-terminally truncated mutants were examined as described above. Molecular masses of multiple polypeptides expressed in the transfected cells were estimated. As shown in (***C***), when the n-region of TM1 was deleted to yield $Nrf1^{\Delta2\text{-}11}$, its 120-kDa protein could be folded into an improper membrane-topology or in an ill-natured orientation so as to be buried in the ER lumen and protected by membranes against cytosolic protease attack, leading to its accumulate, as accompanied by decreases in the expression of other multiple proteoforms. Upon progressively increased deletions of the N-terminal TM-containing portions from within Nrf1 (***C*** & ***D***), all the resulting mutants gave rise to a major protein band of between ~85-kDa and ~75-kDa, along with no or less abundance of short isoforms. This demonstrates that such Nrf1 mutants lacking its N-terminal TM1-adjoining signal peptde cannot be protected by ER membranes and thus become more susceptible to proteolytic degradation. Relative to other N-terminally truncated mutants such as $Nrf1^{\Delta2\text{-}80}$ and $Nrf1^{\Delta2\text{-}90}$ (in both the essential UBL is partially disrupted), $Nrf1\triangle^{2\text{-}36}$ (in which most of UBL is retained but not monitored by membranes), $Nrf1^{\Delta2\text{-}106}$ (in which its PEST1 sequence could be exposed as a degron) and $Nrf1^{\Delta2\text{-}170}$ (in which most of Neh2L is exposed to act as a degron) thus appeared to be unstable even in the presence of MG132, suggesting that they might be rapidly degraded through proteasome-independent proteolysis pathways.

### Figure S10

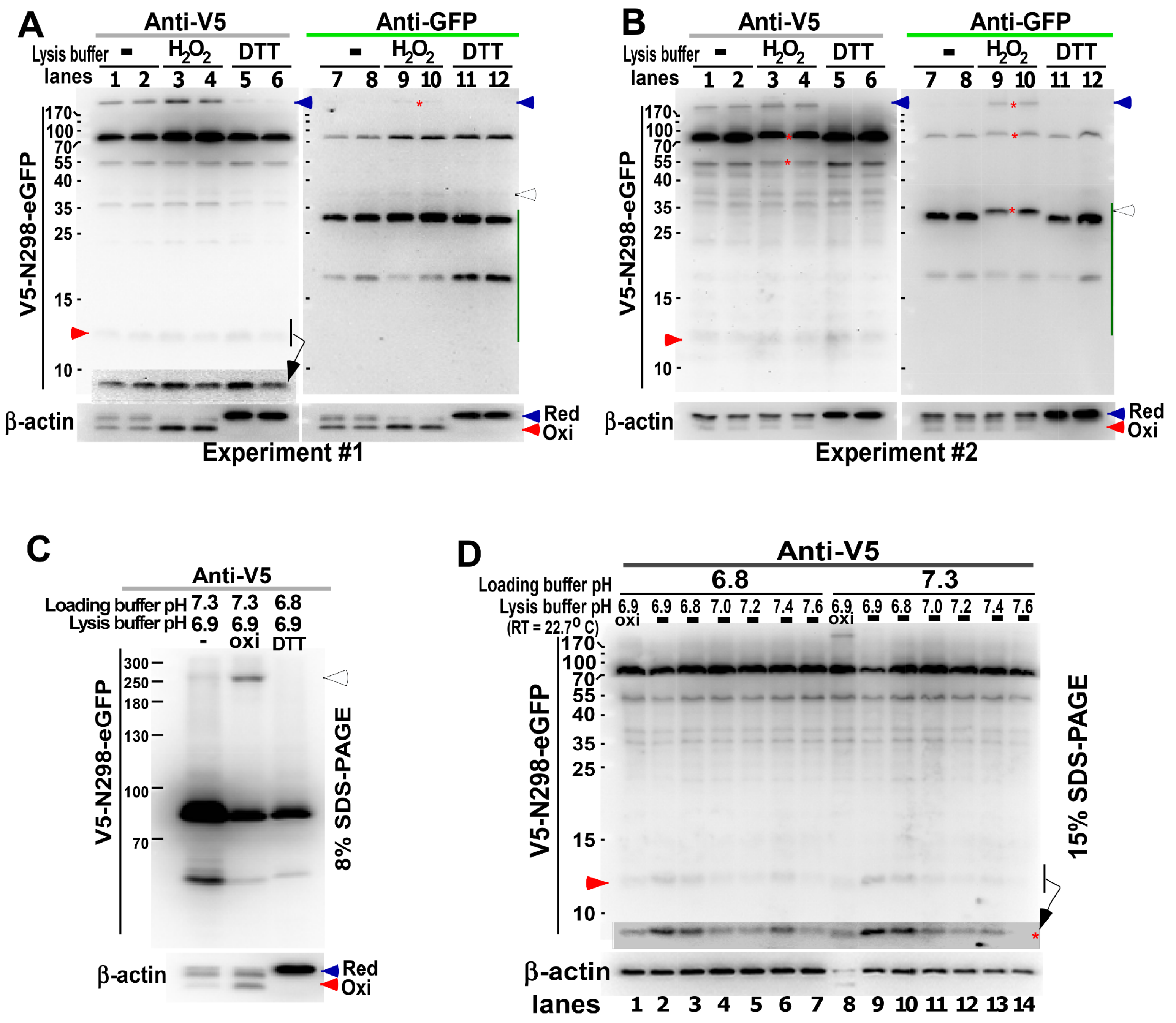


**Figure S10. Formation of several high molecular-weight Nrf1 proteoforms is affected by changes in both redox states and pH values in the micro-environments.**

Total lysates of COS-1 cells expressing V5-N298-eGFP were incubated for 15 min with vehicle (–), $H_2O_2$ (5 mmol/L) or DTT (5 mmol/L) in lysis buffer at distinct pH values, before being visualized by Western blotting with distinct antibodies against V5 or GFP. In addition, some contrasting images were cropped from their originals (as indicated by arrows and/or asterisks).

**Redox electrophoresis** was employed to examine redox-modified Nrf1 or its fusion proteins. COS-1 cells were transfected with an expression construct containing a fragment of cDNA encoding the N-terminal 298 aa of Nrf1. After a 18-h recovery, the cells were harvested in a lysis buffer (5% SDS, 10% NP-40, 1 mmol/L PMSF, pH 7.5). Each sample of cell lysates was split into three parts, that were sonicated in the following respective reaction with vehicle, 5 mM $H_2O_2$ and 5 mM DTT before being immediately denatured for 10 min at 100℃, and then reduced for 5 min with a DTT-free 3× loading buffer (187.5 mmol/L Tris-HCl, pH 6.8, 6% SDS, 30% Glycerol, 0.3% Bromphenol Blue) at 100℃. These resulting reactants were subjected to electrophoresis and Western blotting analysis.

**The results revealed that formation of several high molecular-weight Nrf1 proteoforms is dependent on optimal redox states and pH values in the micro-environments.** Slow electrophoretic mobility of V5-NTD-eGFP and V5-N298-eGFP displays with several higher molecular weights than their original masses. This implies that both may also be oligomerized or aggregated, besides ubiquitinated, probably through the NTD and AD1 portions of Nrf1 (Fig.3E). Herein, we examined whether such mobility of these higher molecular weight forms is affected by changes in either redox states or pH values. Of note, one major slowly-migrated polypeptide of ~260-kDa (which may arise from oligomerization or aggregation of Nrf1 in the fusion context) was highlighted upon oxidative incubation of V5-N298-eGFP with 5 mmol/L of $H_2O_2$ (Fig. S10***A***, *lanes 3, 4*). Conversely, they almost completely disappeared after another fraction of the same samples was allowed for incubation with DTT, a reducing agent (*lanes 5, 6*). In additional experimental conditions, such incubation with oxidizing, rather than reducing, agents caused a slower electrophoretic migration of the ~260-kDa polypeptide, along with the original full-length ~80-kDa protein of V5-N298-eGFP and its processed N-terminal ~55-kDa polypeptide (Fig. S10***B***, *lanes 3, 4*). The C-terminal ~28.5-kDa polypeptides seemed similar to free GFP in size (*lanes 9, 10*), implying that an oxidative reaction may also occur within the epitope of GFP *per se*. Further examinations revealed that the presence of the ~260-kDa polypeptide in oxidative conditions was optimized at pH values (of 6.9 in lysis buffer and 7.3 in loading buffer) (Fig. S10,***C*** and ***D***). Also, the N-terminal ~12.5-kDa polypeptide was obviously enhanced at pH 6.8-6.9 in lysis buffers (Fig. S5, *lanes 2, 3, 9, 10*), but disappeared at pH 7.6 in lysis buffer plus pH 7.3 in loading buffer (*lane 14*). By contrast with an increase in the N-terminal ~55-kDa polypeptide arising from V5-N298-eGFP, its N-terminal ~12.5-kDa polypeptide appeared to be almost unaffected by either oxidizing or reducing agents (Fig. S10***A***). This is also supported by the finding that β-actin was also separated into both oxidative and reduced isoforms.

**Table S1 Pairs of forward and reverse primers used for qPCR herein**

| Gene | Primer (5' → 3') |
|---|---|
| *DDI-1* | F: AGAGTTTTGGACAAGTGACGATGC<br>R: CAAGCCTGGCTCATAATGGTCATC |
| *DDI-2* | F: GTATAGCTGTGCCTGGCACATC<br>R:ATTGTCCAA GCCCTGAGGAGATG |
| *PSMA4* | F: GTCCCTTTGGTGTTTCATTGCTG<br>R: CTTCCATCCCCCGTAATTTCCAC |
| *PSMA7* | F: ATTAAGCTGGTGATCAAGGCACTC<br>R: GGATTGATCTCGCCTCATGACAG |
| *PSMB4* | F: AGATGAACCCTTTGTGGAACACC<br>R: AGCCAAGTATGCACCATAACCAG |
| *PSMB6* | F: TATGGGGGGTATGATGGTAAGGC<br>R: CAGACACTCTTCCTTGGTCATGC |
| *PSMC1* | F: CTGATGATGTAACCCTGGACGAC<br>R: CGTTCTCTTAAGGCCATCAGACC |
| *PSMD12* | F: TATCTGTTGATGAGTCCGAAGCC<br>R: GATCCTTGGGTCTCTGGAAGTTG |
| *RPL13A* | F: CATAGGAAGCTGGGAGCAAG<br>R: GCCCTCCAATCAGTCTTCTG |
| β-*actin* | F: CATGTACGTTGCTATCCAGGC<br>R: CTCCTTAATGTCACGCACGAT |
